# Fast and flexible joint fine-mapping of multiple traits via the Sum of Single Effects model

**DOI:** 10.1101/2023.04.14.536893

**Authors:** Yuxin Zou, Peter Carbonetto, Dongyue Xie, Gao Wang, Matthew Stephens

## Abstract

We introduce mvSuSiE, a multi-trait fine-mapping method for identifying putative causal variants from genetic association data (individual-level or summary data). mvSuSiE learns patterns of shared genetic effects from data, and exploits these patterns to improve power to identify causal SNPs. Comparisons on simulated data show that mvSuSiE is competitive in speed, power and precision with existing multi-trait methods, and uniformly improves over single-trait fine-mapping (SuSiE) performed separately for each trait. We applied mvSuSiE to jointly finemap 16 blood cell traits using data from the UK Biobank. By jointly analyzing the traits and modeling heterogeneous effect sharing patterns, we discovered a much larger number of causal SNPs (>3,000) compared with single-trait fine-mapping, and with narrower credible sets. mvSuSiE also more comprehensively characterized the ways in which the genetic variants affect one or more blood cell traits; 68% of causal SNPs showed significant effects in more than one blood cell type.

## Introduction

Genome-wide association analyses (GWAS) have been performed for thousands of traits and have identified many genomic regions associated with diseases and complex traits [1–4]. Many statistical fine-mapping methods have been developed to prioritize putative causal SNPs for a single trait [5–16], but much fewer methods are available to finemap multiple traits simultaneously. A simple strategy to finemap multiple traits is to finemap each trait separately, then integrate the results *post hoc*. However, integration of results is not straightforward; for example, it is difficult to say whether signals identified in different single-trait analyses correspond to the same underlying causal SNP. Further, analyzing each trait independently is inefficient in that it cannot exploit the potential for increased power of a multivariate analysis [17]. Therefore, it is desirable to finemap the traits simultaneously—that is, to perform *multi-trait fine-mapping*.

Although several methods have been developed for multi-trait fine-mapping [18–26] (Table 1), these methods have important practical limitations. For example, several methods are computationally impractical for more than a small number of traits, and most methods make restrictive assumptions about how SNPs affect the traits, such as that the effects of causal SNPs are uncorrelated among traits. These assumptions are easily violated in fine-mapping applications; for example, in the blood cell traits considered in this paper, some genetic effects are specific to subsets of the traits (e.g., red blood cell traits). There are also several methods developed for the problem of colocalization of two traits (e.g., [27–30]), which has different analysis aims, but overlaps with multi-trait fine-mapping.

**Table 1.** Overview of available statistical methods for multi-trait fine-mapping. Sample runtimes were obtained by running on data sets with *J* = 5,000 SNPs, *N* = 250,000 individuals (only relevant to methods that do not accept summary data), and *R* = 2 or 20 traits. When possible, the upper limit on the number of causal SNPs, *L*, was set to 10. In our tests, PAINTOR ran for a very long time when allowing 3 or more causal SNPs, so we set *L* = 2. (This was without the “MCMC” option, because at the time of our trials the “MCMC” option produced unreasonable results.) moloc was computationally impractical with more than 4 traits. See the Supplementary Note for further details and explanation of the table columns. ^§^MFM is for multiple case-control traits with a shared set of controls. ^†^flashfm’s properties depend on the single-trait fine-mapping method; to illustrate, we used FINEMAP [10]. flashfm with FINEMAP was limited to at most 5 traits. (Another flashfm interface allows up to 6 traits.) ^‡^Calculation of CSs is trivial when limiting to at most 1 causal SNP.

| Method | upper limit on number of causal SNPs | data accepted |  |  | allows correlated traits | models effect sharing | sample runtimes |  | software | version |
| --- | --- | --- | --- | --- | --- | --- | --- | --- | --- | --- |
|  |  | summary | sufficient | CSs |  |  | 2 traits | 20 traits |  |  |
| mvSuSiE | user-specified | yes | yes | yes | yes | yes | 41 s | 2 min | R | 9f28916 |
| flashfm<sup>†</sup> [20] | 10 | yes | yes | yes | yes | yes | 5 min | – | R | 0.0.0.9000 |
| MT-HESS [18] | no limit | no | no | no | yes | yes | >1 day | – | R | 1.99 |
| BayesSUR [21] | no limit | no | no | no | yes | yes | 7 h | – | R | 2.0-1 |
| msCAVIAR [22] | user-specified | yes | no | no | no | yes | >1 day | – | command-line | 0.1 |
| CAFEH [23] | user-specified | yes | yes | yes | no | no | 20 s | 37 s | Python | 1.0 |
| PAINTOR [19] | user-specified | yes | no | no | no | no | 30 min | – | R | 3.1 |
| MFM<sup>§</sup> [24] | user-specified | no | no | yes | no | no | – | – | R | 0.2-1 |
| HyPrColoc [25] | 1 | yes | no | yes<sup>†</sup> | no | no | <1 s | <1 s | R | 1.0 |
| moloc [26] | 1 | yes | no | yes<sup>†</sup> | no | no | <1 s | – | R | 0.1.0 |

Here we introduce mvSuSiE, a fast and flexible method for multi-trait fine-mapping. The name “mvSuSiE” evokes its origins as an extension of the Sum of Single Effects (SuSiE) model [13] to the multivariate analysis setting. In particular, mvSuSiE combines the SuSiE model with ideas from [31] to learn, in a flexible way, the patterns of shared genetic effects among traits. mvSuSiE automatically adapts to the patterns of effect sharing in the particular traits being analyzed, making it widely applicable to fine-mapping any set of related traits. We also leverage ideas from [16] to allow for the analysis of summary statistics generated from a genetic association study, which are often more accessible than individual-level data [32, 33]. mvSuSiE is computationally practical for jointly fine-mapping many traits in “biobank scale” data sets. We demonstrate its effectiveness compared with existing methods in simulations and by fine-mapping 16 blood-cell traits in 248,980 UK Biobank samples.

## Results

### Methods overview

Consider fine-mapping *R* traits in a region containing *J* SNPs (or other biallelic loci). For each individual *i* = 1,…,*N*, let *y*_*ir*_ denote trait *r* measured individual *i*, and let *x*_*ij*_ denote the genotype of individual *i* at SNP *j*, encoded as the number of copies of the minor allele. We perform multi-trait fine-mapping using the following multivariate linear regression model:

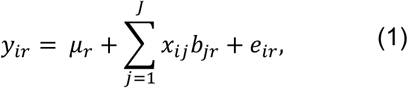

where *μ*_*r*_ reflects the mean of trait *r, b*_*jr*_ is the effect of SNP *j* on trait *r*, and the *e*_*ir*_’s are normally distributed error terms (which may be correlated among the *R* traits). Within this regression model, we frame fine-mapping as a “variable selection problem”: most SNPs are assumed to have no effect on any trait—that is, most effects *b*_j*r*_ are zero—and the goal of multi-trait fine-mapping is to identify which SNPs have a non-zero effect on which traits, and to assess uncertainty in these inferences. (For brevity, we use the term “causal SNP” to mean a SNP with a non-zero effect.) Our mvSuSiE method achieves this goal by extending the *Sum of Single Effects* (SuSiE) model [13] to the multivariate setting. By extending ideas from [16], mvSuSiE can perform fine-mapping using either individual-level data (genotypes and phenotypes) or summary data (e.g., LD matrix and marginal *z*-scores); see Methods and the Supplementary Note for details.

Among existing approaches to fine-mapping, mvSuSiE is most closely related to CAFEH [23], which also extends SuSiE to perform multi-trait fine-mapping. Both CAFEH and mvSuSiE inherit much of the simplicity and benefits of single-trait SuSiE. Like SuSiE, both mvSuSiE and CAFEH require the user to specify an upper bound, *L*, on the number of causal SNPs in a region, and are robust to this upper bound being larger than needed. And both methods exploit SuSiE’s simple fitting procedure, Iterative Bayesian Stepwise Selection (IBSS) [13]. IBSS is similar to standard forward stepwise selection, but improves on it (i) by using Bayesian computations to take into account uncertainty in which SNPs are selected at each step and (ii) by iterating through selection events to allow errors in initial selections to be corrected as fitting progresses. However, mvSuSiE also improves on CAFEH in two key ways:

a. mvSuSiE uses a flexible prior distribution—specifically, a mixture of multivariate normal distributions, as in [31]—to model effect sharing patterns across traits. Further, the parameters of this prior are estimated from the data, allowing mvSuSiE to adapt to each data set. This flexible approach allows for different causal SNPs that show different patterns of association; for example, in analyses of blood cell traits (below), mvSuSiE learns that some SNPs affect primarily red blood cell (erythrocyte) traits, some affect primarily white blood cell (leukocyte) traits, and some affect both, or a subset of one or the other. In contrast, CAFEH assumes a less flexible and less adaptive prior in which causal effects are independent across traits.
b. mvSuSiE allows for correlations in measurements among traits, with these correlations again being estimated from the data. In contrast, CAFEH assumes measurements are independent across traits, which is an inappropriate assumption in association studies involving correlated traits.

For (a), estimating the prior distribution from the data involves combining information across many causal SNPs from many regions, which is an additional step compared with standard single-trait fine-mapping analyses. This additional step can be avoided by using a simpler fixed prior (see Supplementary Note), but at potential loss of power.

We also introduce novel ways to summarize the inferences from multi-trait fine-mapping. Again, this builds on SuSiE, which summarizes single-trait results by reporting, for each SNP, a “posterior inclusion probability” (PIP) quantifying the probability that the SNP is causal, and by reporting “credible sets” (CSs) [7, 13] that are designed to capture, with high probability, at least one causal SNP. Informally, each CS represents an independent association signal in the data, and the size of a CS (*i*.*e*., the number of SNPs in the CS) indicates how precisely one can pinpoint the causal SNP underlying this signal. For multi-trait analyses, it may seem natural to report PIPs and CSs separately for each trait. However, this raises thorny issues: for example, if the reported CSs for two traits overlap, do these represent the same signal with a single underlying causal SNP, or different signals with multiple causal SNPs? To avoid these problems, we separate inference into two questions.

#### First question

Which SNPs are causal for *at least one trait*? This question is answered by *cross-trait PIPs and CSs* that summarize the inferences across all traits.

#### Second question

For each causal SNP (*i*.*e*., CS) identified, which traits does it affect? This is answered by computing a *trait-wise* measure of significance, the *local false sign rate* (*lfsr*) [31, 34], for each SNP in each trait. (A small *lfsr* indicates a high confidence in the *sign* of the effect.) Because SNPs in a CS are typically in high LD, their trait-wise *lfsr* values are typically similar, and it is convenient to use a single number, the *average lfsr*, as a trait-wise measure of significance of each CS. If the *average lfsr* for trait *r* is small, this indicates high confidence in the sign of the effect—that is, a small posterior probability that the true effect is zero or that its estimated sign is incorrect—and we say the CS is “significant for trait *r*.”

In summary, the reported results from a mvSuSiE analysis are the *cross-trait PIPs and CSs* together with *trait-wise* measures of significance (*lfsr*) for each SNP and each CS in each trait. Fig. 1 summarizes the mvSuSiE analysis workflow for a typical genetic association study.

**Figure 1.**
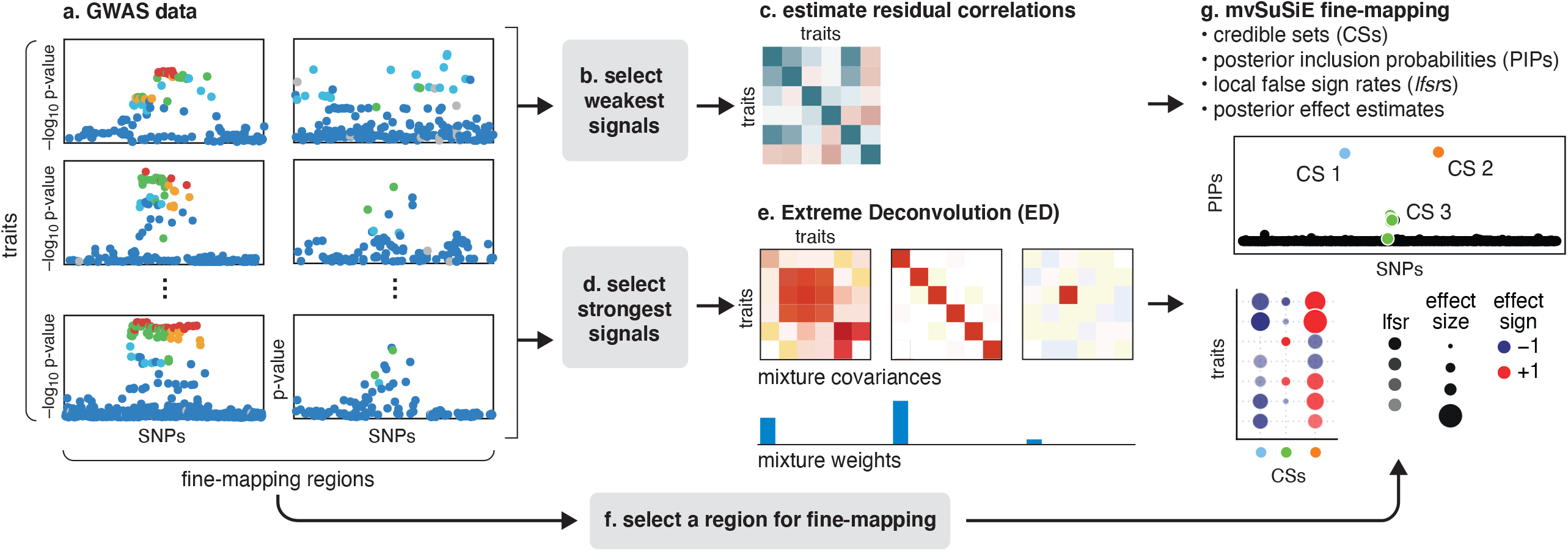
Overview of multivariate fine-mapping using mvSuSiE. mvSuSiE accepts as input *R* traits and SNP genotypes measured in *N* individuals, and *M* target fine-mapping regions. Alternatively, mvSuSiE-RSS accepts SNP-level summary statistics (a) computed from these data. The weakest SNP association signals are extracted from these data (b), which are used in (c) to estimate correlations in the trait residuals. Separately, the strongest association signals are extracted (d) to estimate effect sharing patterns (e) using Extreme Deconvolution (ED) [35]. Finally, the effect-sharing patterns estimated by ED, together with the estimated weights, are used to construct a prior for the unknown multivariate effects, and this prior is used in mvSuSiE to perform multivariate fine-mapping simultaneously for all SNPs within a selected region (f, g). Steps f and g are repeated for each fine-mapping region of interest. The key mvSuSiE outputs are: a list of credible sets (CSs), each of which is intended to capture a causal SNP; a posterior inclusion probability (PIP) for each SNP giving the probability that the SNP is causal for at least one trait; *average local false sign rates* summarizing the significance of each CS in each trait; and SNP-wise posterior effect estimates on each trait. For example, if a region contains 3 causal SNPs, mvSuSiE will, ideally, output 3 CSs, each containing a true causal SNP, with the *average lfsr* indicating which traits are significant for each CS. See Methods for definitions.

### Evaluation in simulations using UK Biobank genotypes

We compared mvSuSiE with existing multi-trait fine-mapping methods and a single-trait fine-mapping method, SuSiE [13, 16], in simulations. Among available multi-trait fine-mapping methods (Table 1), MT-HESS [18] and BayesSUR [21, 36, 37] are similar to mvSuSiE in features and modeling assumptions, but are computationally impractical for large fine-mapping data sets. msCAVIAR [22] shares the ability of mvSuSiE to model effect sharing, but is designed for analyzing data from multiple studies, and therefore makes modeling assumptions that are less appropriate for analyzing multiple traits. MFM [24] is another multi-trait fine-mapping method, but is specific to multiple case-control traits with a shared set of controls. Therefore, we focussed our comparisons on CAFEH [23] which can handle large multi-trait fine-mapping data sets. We also compared with flashfm [20] and PAINTOR [19] on smaller fine-mapping data sets with two traits.

To make our simulations reflective of current large-scale genomic data sets, we obtained imputed genotype data from the UK Biobank [38] and simulated quantitative traits with 1–5 simulated causal SNPs in each fine-mapping region. We simulated from a variety of effect-sharing patterns, with effect sizes scaled to roughly reproduce the distributions of *z*-scores observed in genome-wide association analyses of complex traits from UK Biobank data. The fine-mapping regions were drawn from autosomal chromosomes and varied in size (0.4–1.6 Mb), number of SNPs (1,000–5,000 SNPs), and LD patterns.

We simulated traits under two scenarios:

a. “Trait-specific + Shared Effects,” in which SNP effects on 20 independent traits were either specific to one trait, or shared among traits in simple ways (e.g., equal effects on a pair of traits and no effect on the remaining traits);
b. “Complex Shared Effects,” in which SNP effects on 16 correlated traits were generated from a variety of sharing patterns derived from the UK Biobank blood cell traits.

To compare with PAINTOR and flashfm, we also simulated smaller data sets with 2 independent traits and shared effects.

We compared methods in their detection of cross-trait causal SNPs—in which we define a cross-trait causal SNP as one that affects at least one trait—and trait-wise causal SNPs. We assessed the performance of both SNP-wise measures (e.g., PIPs) and credible sets (CSs) for these tasks. In practice, we recommend focusing on CS-based inferences (Fig. 2c, d) rather than SNP-wise measures (Fig. 2a, b) because the CSs account for uncertainty in the causal SNP due to LD. (In our multi-trait fine-mapping of blood cell traits below, we focussed on CS-based inferences.) However, not all competing methods provide comparable CS-based inferences (e.g., CAFEH does not provide trait-wise CSs), so for completeness and to allow comparisons with other methods, we also evaluated performance of SNP-wise significance measures (Fig. 2a, b).

**Figure 2.**
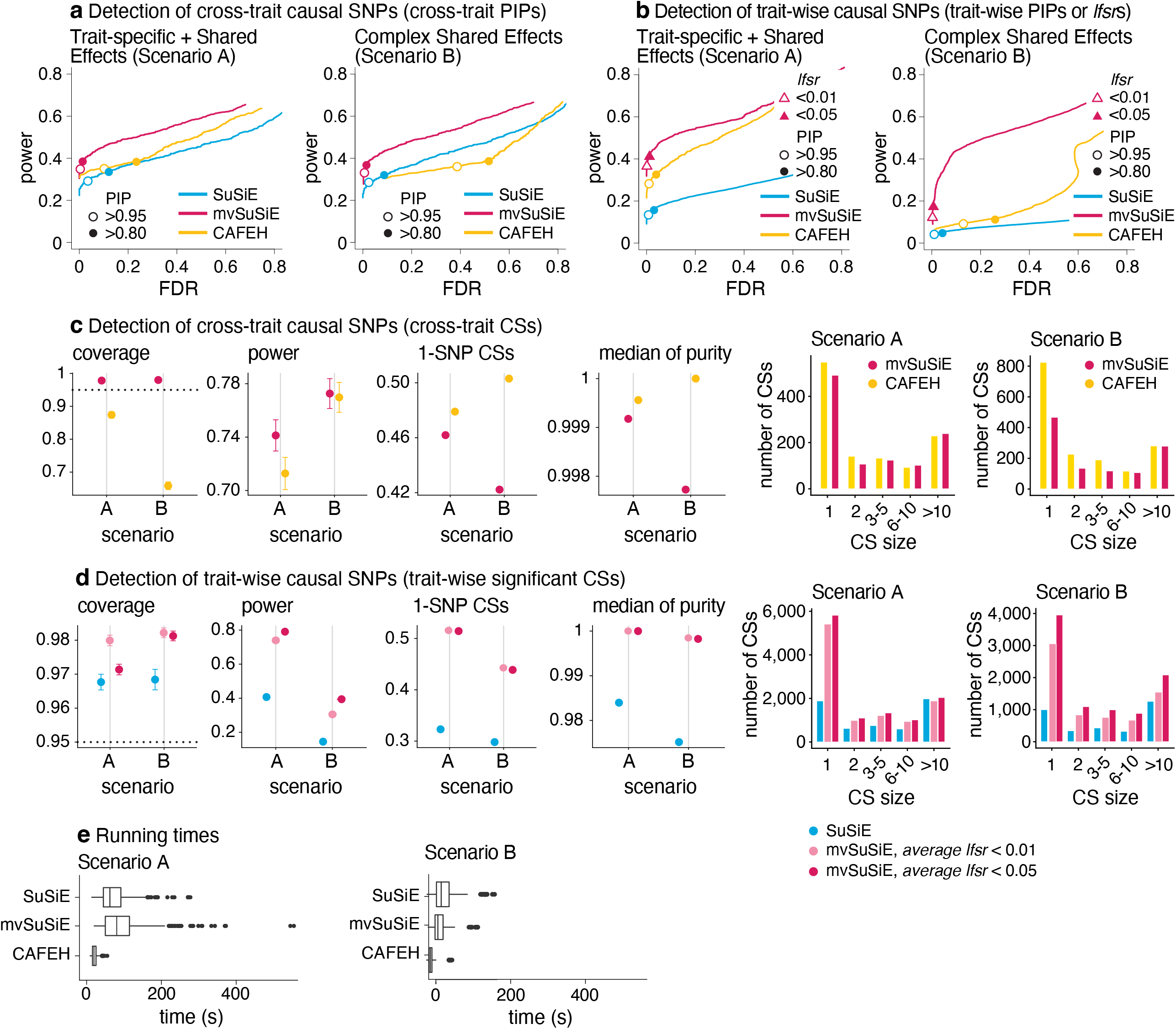
Comparison of fine-mapping methods in simulated data. Parts a and b show power vs. FDR in identifying (a) cross-trait or (b) trait-wise causal SNPs, using SNP-wise measures. In each scenario, FDR and power were calculated by varying the measure threshold from 0 to 1 (*n* = 600 simulations). FDR = FP/(TP + FP) and power = TP/(TP + FN), where FP, TP, FN, TN denote, respectively, the number of false positives, true positives, false negatives and true negatives. The specific SNP-wise measures used in a are PIP (mvSuSiE, CAFEH), max-PIP (SuSiE); in b, PIP (SuSiE), *min-lfsr* (mvSuSiE) and study PIP (CAFEH). In a and b, power and FDR at specific thresholds are indicated by the circles and triangles. Note that the thresholds for the different methods are not equivalent or comparable; results are shown at these thresholds for illustration only. See also Supplementary Table 1. Parts c and d evaluate the 95% CSs from the *n* = 600 simulations using the following metrics: *coverage*, the proportion of CSs containing a true causal SNP; *power*, the proportion of true causal SNPs included in at least one CS; the proportion of CSs that contain a single SNP (“1-SNP CSs”); and *median purity*, in which “purity” is defined as the smallest absolute correlation (Pearson’s *r*) among all SNP pairs in a CS. Histograms of CS sizes (number of SNPs in a 95% CS) are given for each scenario. Target coverage (95%) is shown as a dotted horizontal line. Error bars show 2 times the empirical s.e. from the results in all simulations. Part e summarizes runtimes (*n* = 600 simulations); the SuSiE runtimes are for running SuSiE independently on *all* traits. The box plot whiskers depict 1.5 times the interquartile range, the box bounds represent the upper and lower quartiles (25th and 75th percentiles), the center line represents the median (50th percentile), and points represent outliers. Note that SuSiE analyzes each trait independently and therefore is not included in c. CAFEH does not provide trait-wise CSs and therefore is not included in d.

In all our comparisons, mvSuSiE improved power, coverage and resolution (purity, proportion of 1-SNP CSs) over the SuSiE single-trait analyses (Fig. 2a, b, d; *n* = 600 simulations). The greatest gains were in Scenario B, where mvSuSiE had the advantage that it accounted for correlations among traits. Comparing CAFEH and single-trait SuSiE in SNP-wise inferences, CAFEH improved performance in Scenario A but performed slightly less well in detecting causal SNPs in Scenario B (Fig. 2a, b). CAFEH also produced poorly calibrated PIPs in Scenario B (Supplementary Fig. 1, Supplementary Table 1); for example, CAFEH at a seemingly stringent “study PIP” threshold of 0.95 resulted in an FDR of 0.13 that was much higher than mvSuSiE at an *lfsr* threshold of 0.05 (FDR = 0.0065). This also illustrates the difficulty of setting comparable thresholds for the different quantities outputted by different methods; therefore, following common practice in statistical fine-mapping papers, we presented results using power-FDR curves to sidestep this difficulty.

Comparing CSs (Fig. 2c, d), CAFEH improved the purity of the CSs and the proportion of 1-SNP CSs, but these improvements were tempered by CAFEH’s reduced power and coverage, particularly in Scenario B. A partial explanation for these results is that Scenario B contradicts CAFEH’s assumptions of independent traits and independent causal effects. In support of this explanation, when we forced mvSuSiE to make the same independence assumptions as CAFEH, mvSuSiE’s performance was reduced and the PIPs were also poorly calibrated (see the “random effects prior” and “independent traits” results in Supplementary Figures 1–3).

These results illustrate the benefits of having a flexible model that can adapt to different fine-mapping scenarios by learning effect-sharing patterns from the data (Supplementary Figures 2, 4–7). This flexibility comes at a computational cost—CAFEH was consistently faster than mvSuSiE (Fig. 2e, Supplementary Table 2)—but mvSuSiE was still fast enough to handle the largest fine-mapping data sets we considered.

We also compared mvSuSiE with CAFEH, PAINTOR and flashfm in a variety of simpler fine-mapping data sets simulated in a similar way to above but with only two traits (Supplementary Figures 8–15). Even when the traits were simulated independently in accordance with PAINTOR’s modeling assumptions, PAINTOR had much lower power to detect causal SNPs than both SuSiE and mvSuSiE (Supplementary Fig. 8a). Both flashfm and mvSuSiE improved power over the SuSiE single-trait analyses, but mvSuSiE achieved much greater gains in power (Supplementary Figures 8–15). mvSuSiE also had considerably lower computational cost than PAINTOR and flashfm (Supplementary Fig. 9, Supplementary Table 2). The performance of CAFEH in these simpler simulations was similar to mvSuSiE except when the two traits were highly correlated (Supplementary Figures 10, 11).

In summary, these simulations demonstrate the benefits of mvSuSiE as an efficient and flexible multi-trait fine-mapping method. In particular, mvSuSiE consistently increased power to detect causal SNPs, improved precision (reduced CS size) compared with fine-mapping each trait separately, and was the only method that provided both cross-trait and trait-wise significance measures.

### Multi-trait fine-mapping of blood cell traits from UK Biobank

To illustrate mvSuSiE in a substantive application, we fine-mapped blood cell traits using data from the UK Biobank [38]. Previous analyses of these data include association analyses [39, 40] and single-trait fine-mapping [41, 42], but multi-trait fine-mapping using mvSuSiE has the potential to improve power and precision of fine-mapping. Multi-trait fine-mapping is also better for answering questions about shared genetic effects—which SNPs affect which traits—and hence provide insights into the underlying biology.

Focusing on a subset of 16 blood cell traits (Supplementary Table 3), we performed standard PLINK association analyses [43] with *n* = 248,980 UK Biobank samples for which all 16 traits and imputed genotypes were available (Methods). We included covariates such as sex and age, as well as genotype principal components to limit spurious associations due to population structure. From the results of these association analyses, we obtained 975 candidate genomic regions for fine-mapping (Supplementary Table 4). We then applied the mvSuSiE analysis pipeline to these 975 candidate regions (Methods). To understand the benefits of a multi-trait fine-mapping, we also ran SuSiE on the same regions, separately for each trait.

### Genetic relationships among blood traits inform discovery of multi-trait causal SNPs

From the 975 candidate regions, mvSuSiE identified 3,396 independent causal signals (95% cross-trait CSs). The median size of a CS was 7 SNPs. Among these CSs, 726 contained just one SNP (“1-SNP CS”); therefore, mvSuSiE identified 726 high-confidence candidate causal SNPs (PIP > 0.95; Supplementary Table 5). Several of these 1-SNP CSs (36) were not identified in *any* of our single-trait (SuSiE) analyses, underscoring the benefits of combining evidence across genetically related traits. Reassuringly, 496 of the 726 SNPs were also identified as high-confidence causal SNPs (PIP > 0.95) in the single-trait analyses of [42], and 145 of 726 overlapped with [41].

The number of CSs significant in each trait (*average lfsr* < 0.01) ranged from 370 (basophil percentage) to 1,423 (platelet count), and the number of 1-SNP CSs ranged from 108 to 335 (Fig. 3c). (Note that 10 of the 3,396 CSs were not significant in any traits at *average lfsr <* 0.01.) Notably, mvSuSiE increased fine-mapping discovery and resolution compared to SuSiE single-trait fine-mapping: the number of trait-wise significant CSs increased, on average, 2.2-fold compared with SuSiE, and the number of trait-wise significant 1-SNP CSs increased, on average, 3.5-fold (Fig. 3c).

**Figure 3.**
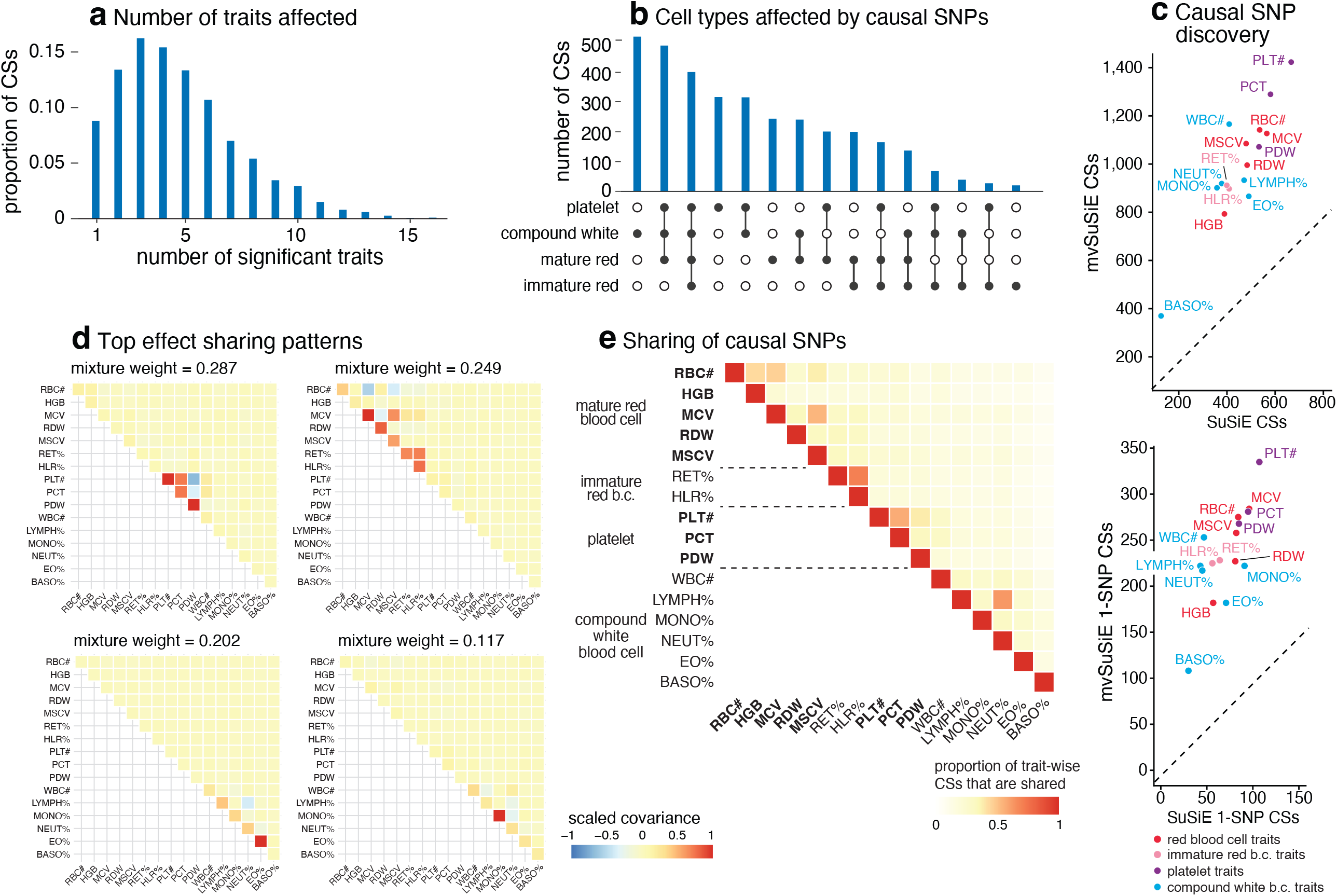
mvSuSiE fine-mapping and primary effect sharing patterns in UK Biobank blood cell traits. Parts a, b and e give summaries of the 3,396 mvSuSiE CSs identified from the 975 candidate fine-mapping regions: (a) number of significant (*average lfsr* < 0.01) traits in each CS; (b) significant traits in CSs grouped by blood cell-type subsets; (e) pairwise sharing of significant CSs among the traits. In e, for each pair of traits we show the ratio of the number of CSs that are significant in both traits to the number of CSs that are significant in at least one trait. (c) Number of CSs and 1-SNP CSs for each trait identified by SuSiE and mvSuSiE (after removing CSs with purity less than 0.5). In c, each mvSuSiE count is the number of mvSuSiE CSs or 1-SNP CSs that are significant (*average lfsr* < 0.01) for the given trait. (d) Covariance matrices in the mvSuSiE data-driven prior capturing the top sharing patterns (these are the covariance matrices with the largest mixture weights in the prior). The covariance matrices were scaled separately for each plot so that the plotted values lie between –1 and 1. See Supplementary Fig. 5 for the full set of 15 sharing patterns.

The fine-mapped SNPs from mvSuSiE were generally slightly more enriched for genomic regulatory annotations than those for SuSiE (Supplementary Fig. 16), providing indirect support for the additional mvSuSiE findings being driven by real signals rather than false positives. For example, the mvSuSiE-fine-mapped SNPs had an enrichment odds ratio of 11.9 for being an eQTL compared to 9.7 from SuSiE. We also analyzed enrichment of the fine-mapped SNPs for accessible chromatin regions in hematopoietic cell-types [41] (Supplementary Figures 17, 18 and Supplementary Tables 6, 7). Similar to [42], both the SuSiE and mvSuSiE results showed some of the expected enrichments such as enrichment of SNPs affecting platelet-related traits for open chromatin in platelet-producing megakaryocytes.

mvSuSiE improved discovery and resolution over single-trait analysis by learning and exploiting patterns of shared (and not shared) genetic effects from the data. In these data, the most prominent learned patterns involved strong sharing of effects amongst traits for the same blood cell type (Fig. 3d). However, many other patterns were also identified (Supplementary Fig. 5), including both trait-specific and broad effects, suggesting that SNPs can affect blood cells in a wide variety of ways, presumably reflecting a wide variety of underlying biological mechanisms. By applying mvSuSiE with a prior that incorporates these learned sharing patterns, we obtained a genome-wide summary that underscores the diversity of genetic effects on blood cell traits (Fig. 3a, b, d). Genetic effects are more commonly shared among traits within the same blood cell type as one might expect (Fig. 3e), but SNPs affecting multiple blood cell types are also common (Fig. 3b).

### Multi-trait fine-mapping reveals highly heterogeneous genetic determination of blood traits

To illustrate the potential for mvSuSiE to help dissect complex genetic association signals, we examine four example blood cell trait loci in more detail (Fig. 4).

**Figure 4.**
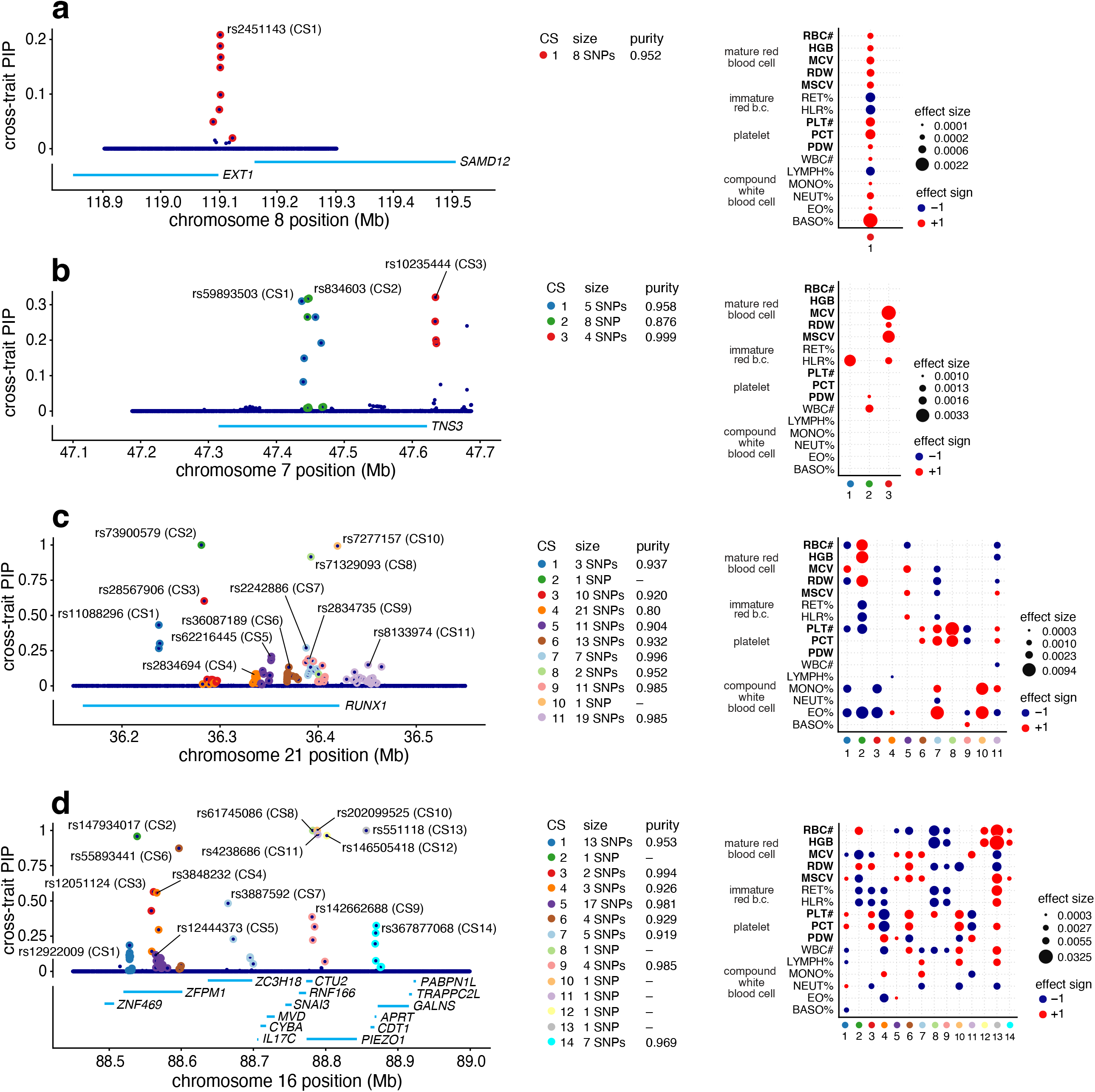
Examples of blood cell trait loci fine-mapped using mvSuSiE. The left-hand plot in each panel a–d is a “PIP plot” showing cross-trait posterior inclusion probabilities (PIPs) for each SNP. The cross-trait PIP is an estimate of the probability that the SNP is causal for at least one trait. The labeled SNPs are the “sentinel SNPs”, the SNPs with the highest cross-trait PIP in each CS. “Purity” is defined as the minimum absolute pairwise correlation (Pearson’s *r*) among SNPs in the CS. The right-hand plots a–d shows the posterior effect estimates of the sentinel SNPs (only for CSs that are significant for the given trait, with *average lfsr* < 0.01). All estimates and tests are from a data sample of size *n* = 248,980.

Fig. 4a shows the mvSuSiE results for the *EXT1-SAMD12* locus. Single-trait association analysis of this region shows only one trait, basophil percentage, with a genome-wide significant association (PLINK two-sided *t*-test *p*-value < 5 × 10^-8^). Similarly, single-trait fine-mapping with SuSiE identified a single CS for basophil percentage containing 10 candidate SNPs, and no CSs ifor other traits. From these results one might conclude that the causal SNP is specific to basophil percentage. However, the mvSuSiE fine-mapping results assess the CS as significant in most traits, suggesting that in fact the causal SNP has broad effects across many traits. (Indeed, all traits had marginal association *p-*values less than 0.003 with the lead SNP, which in some situations might be considered “significant”.) The mvSuSiE CS is smaller than the single-trait CS (8 vs. 10 SNPs), illustrating the improved fine-mapping resolution that can come from combining information across traits (see also Supplementary Fig. 19).

Fig. 4b shows mvSuSiE results for the Tensin 3 locus. Vuckovic et al [42] used single-trait fine-mapping to identify causal signals for several red and white blood cell traits at this locus. However, a single-trait analysis does not tell us whether these signals are due to one or a few causal SNPs affecting many blood cell traits, or due to many causal SNPs affecting individual traits. The multi-trait mvSuSiE analysis identified three causal signals (cross-trait CSs) with three distinct patterns of genetic effect: one mostly affects red blood cell traits (CS3); another has a detectable effect in HLR% only (CS1); and a third has smaller effects in both white blood cell and platelet traits (CS2). The three different patterns suggest that the biological effects of these SNPs are also different, and they suggest a multi-faceted role for *TNS3* in affecting blood-cell traits. This example illustrates the flexibility of mvSuSiE, including its ability to capture different patterns of effect-sharing even within a single locus, and its ability to extract relatively simple inferences in quite complex situations.

Fig. 4c shows a more complex example involving many signals in and around *RUNX1*. SNPs in the *RUNX1* locus have previously been associated with rheumatoid arthritis [44, 45] and other immune-related diseases (e.g., [46, 47]), and colocalization analyses have suggested that the causal SNPs are also associated with eosinophil proportions in blood [42]. Multi-trait fine-mapping results from mvSuSiE suggest a complex picture with 11 signals (cross-trait CSs), each with detectable effects in many different blood cell traits, and some with no detectable effect on eosinophil proportions. These results suggest that the mechanisms by which this gene affects immune-related diseases may be more complex than just through eosinophils, possibly involving many platelet, red blood cell and other white blood cell traits.

Finally, Fig. 4d shows an even more complex example where many causal signals are mapped to a region containing many genes, including *PIEZO1* and *ZFPM1*. This is a gene-dense region with well-studied connections to blood cell traits and blood-related diseases [48–52]. mvSuSiE identified 14 independent signals (cross-trait CSs) in the region. These 14 signals show a wide variety of effect patterns; for example, some are significant in only a few traits related to mature red blood cells (e.g., CS12, CS14), some are significant across a broader range of red blood cell traits (CS2), and some are significant across most traits (CS13). Regions of this level of complexity may take considerable additional investigation to fully understand. Although this is a complex example, we note that of the 14 CSs identified in this region, 7 contain a single SNP, demonstrating that even in complex regions mvSuSiE can identify high-confidence causal SNPs.

## Discussion

We have introduced mvSuSiE, a fast and flexible multi-trait fine-mapping method. mvSuSiE outperformed single-trait fine-mapping methods in both power and resolution. Unlike most available multi-trait fine-mapping methods, mvSuSiE can efficiently analyze dozens of correlated traits and can model complex patterns of effect size variation via a flexible data-driven prior distribution. The prior model also includes as special cases several simpler models that are commonly used in meta-analyses, such as the “fixed effects” model which assumes equal effects in all traits, and the “random effects” model which allows for different effect sizes among traits [53]. These models can be used in place of the data-driven prior to speed up computation if users desire, though at a potential loss of power. See the Supplementary Note for additional discussion, where we discuss some of mvSuSiE’s limitations, and give practical guidance on applying mvSuSiE to other types of traits (e.g., binary traits) and on dealing with other potential complications (e.g., missing data).

## Acknowledgements

We thank Karl Tayeb for helping with CAFEH, Ru Feng for helping with the enrichment analyses, and Fabio Morgante and Anqi Wang for their contributions to the mvsusieR software. We thank the staff at the Research Computing Center and the Center for Research Informatics at the University of Chicago for maintaining the high-performance computing resources used to implement the numerical experiments. This work was supported in part by the NHGRI at the National Institutes of Health under award number R01HG002585 (to MS) and by a grant from the Gordon and Betty Moore Foundation (to MS). GW was supported by the NIA at the National Institutes of Health under award number R01AG076901 and by a grant from the Urbut Family Foundation. This research has been conducted using the UK Biobank Resource under Application Number 27386.

## Author contributions

YZ, GW and MS conceived of the project and developed the statistical methods. YZ implemented the methods comparisons in simulations and performed the statistical analyses, with help from GW and PC. YZ and GW wrote the software, with contributions from MS and PC. YZ and GW prepared the online code and data resources. All authors prepared the results for the manuscript and wrote the manuscript.

## Methods

### Ethics statement

This work used publicly available data sets and so ethical approval was not required.

### Multivariate multiple regression

mvSuSiE is based on a basic multivariate multiple regression model for *R* quantitative traits observed in *N* individuals,

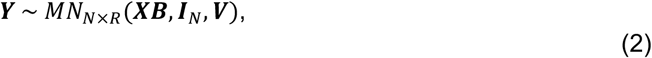

where ***Y*** ∈ ℝ^*N*×*R*^ is a matrix storing *N* observations of *R* traits, ***X*** ∈ ℝ^*N*×*J*^ is a matrix of *N* genotypes at *J* SNPs, ***B*** ∈ ℝ^*J*×*R*^ is a matrix of regression coefficients (“effects”) for the *J* SNPs and *R* traits, ***V*** is an *R* × *R* covariance matrix (we assume ***V*** is invertible), *I*_*N*_ is the *N* × *N* identity matrix, and *MN*_*N*×*R*_(***M*, Σ**^row^, **Σ**^col^< denotes the matrix normal distribution [54, 55] with mean ***M*** ∈ ℝ^*N*×*R*^ and covariance matrices **Σ**^row^, **Σ**^col^ (of dimension *N* × *N* and *R* × *R*, respectively).

### Intercept

We do not explicitly include an intercept in (2). Instead, we account for an intercept implicitly by “centering” the columns of ***X*** and the columns of ***Y*** so that the mean of each column is zero. From a Bayesian perspective, centering the columns of ***X*** and ***Y*** is equivalent to integrating with respect to an (improper) uniform prior on the intercept. (This is a multivariate generalization of the result for univariate regression in [56]. See the Supplementary Note for a more formal proof of this result.) In short, centering eliminates the need to explicitly include an intercept in (2), and we proceed with mvSuSiE assuming that ***X*** and ***Y*** have been centered.

### The mvSuSiE model

mvSuSiE generalizes the “Sum of Single Effects” (SuSiE) model [13] to the multivariate setting:

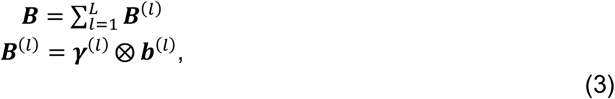

where ***γ***^(*l*)^ ∈ {0,1}^*J*^ is a vector of indicator variables in which exactly one of the *J* elements is one and the remaining are zero, ***b***^(*l*)^ ∈ ℝ^*R*^ is a vector of regression coefficients, and ***u*** ⨂ ***v*** = ***uv***^⊤^ denotes the outer product of (column) vectors ***u*** and ***v***. The coefficients ***B*** defined in this way are a sum of *L* “single effects” ***B***^(*l*)^. In particular, matrix ***B***^(*l*)^ ∈ ℝ^*J*×*R*^ has at most one row containing non-zero values, and these non-zero values are determined by ***b***^(*l*)^. We therefore refer to ***B***^(*l*)^ as a “single effect matrix” because it encodes the effects of a single SNP. The final set of coefficients ***B*** is a matrix with at most *L* rows containing non-zero values.

Similar to SuSiE, we introduce priors for the indicator variables ***γ***^(*l*)^ and regression coefficients ***b***^(*l*)^,

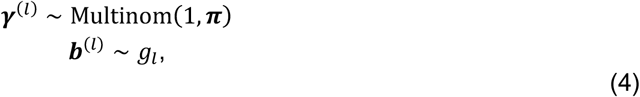

in which Multinom(*m*, ***π***) denotes the multinomial distribution for *m* random multinomial trials with category probabilities ***π*** = (*π*_1_, …, *π*_*J*_<, such that *π*_j_ ≥ 0, ∑^*J*^ *π*_j_ = 1. The *π*_j_’s are the prior inclusion probabilities. By default, we assume a uniform prior; that is, *π*_j_ = 1/*J*, for *j* = 1, …, *J*. (All the results in this paper use this default prior.) Our software implementation of mvSuSiE also support for other choices of ***π***; for example, ***π*** could be determined by external biological information about the SNPs (e.g., [57]).

The prior distribution *g*_*l*_ for each single effect ***b***^(*l*)^ should capture the variety of effect sharing patterns we expect from the multiple traits. To this end, we use a prior similar to the mixture of multivariate normals prior introduced in [31],

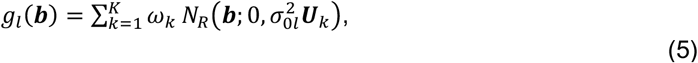

in which each ***U***_*k*_ is a (possibly singular) covariance matrix, 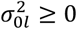 scales the prior for each single effect *l*, ***ω*** = (*ω*_1_, …, *ω*_*K*_) is a vector of mixture weights, such that 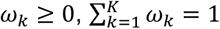, and *N*_*d*_(***x***; ***μ*, Σ**) denotes the multivariate normal distribution on ***x*** ∈ ℝ^*d*^ with mean ***μ*** ∈ ℝ^*d*^ and *d* × *d* covariance **Σ**. The covariance matrices 𝒰 = {***U***_1_, …, ***U***_***K***_} and the mixture weights ***ω*** must be chosen beforehand, whereas prior scaling parameters 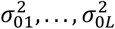 are treated as unknown, and are estimated from the data.

In summary, mvSuSiE is a multivariate regression model with a flexible mixture-of-normals prior on the “single effects” ***b***^(*l*)^. The unknowns of primary interest are the single-effect matrices ***B***^(*l*)^. As we explain in more detail below, we compute a posterior distribution of the single effects, which is then used to compute key fine-mapping statistics, including posterior inclusion probabilities (PIPs) and credible sets (CSs). The scaling factors 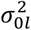 are not of primary interest to the fine-mapping (“nuisance parameters”), and are estimated from the data to aid in better posterior estimation of the single effects. Other model parameters, such as the residual covariance matrix ***V***, are assumed to be known, or should have been estimated previously. In the Supplementary Note, we give guidance on choosing these parameters or estimating them from data. See also the Supplementary Note for derivations, development of the model fitting algorithm, and additional technical details.

### UK Biobank data

The UK Biobank is a prospective cohort study with detailed phenotype and genotype data collected from approximately 500,000 participants recruited in the United Kingdom, with ages between 40 and 69 at time of recruitment [38, 58]. For fine-mapping, we focused on a subset of 16 blood cell traits from the UK Biobank haematology data collection [59]. These blood cell traits were also the focus of a recent association analysis [39, 40] and fine-mapping studies [41, 42]. Several of the UK Biobank blood cell traits are based on the same measured quantities and are therefore highly correlated so we did not include all the blood cell traits in our analyses. For example, relative volume of erythrocytes, also known as “hematocrit” (HCT), is calculated from mean corpuscular volume (MCV) and red blood cell count (RBC#), so to avoid including highly correlated traits we did not include HCT. The blood cell traits used in our fine-mapping analyses are summarized in Supplementary Table 3.

The UK Biobank imputed genotypes feature a high density of available SNPs so they are well suited for fine-mapping. We used a subset of the 502,492 available UK Biobank genotypes (version 3), removing samples that met one or more of the following criteria for exclusion: mismatch between self-reported and genetic sex; pregnant; one or more data entries needed for the analysis or data preparation steps are missing; and, following [39, 42], a blood-related disease was reported in the hospital in-patient data (blood-related diseases included were leukemia, lymphoma, bone marrow transplant, chemotherapy, myelodysplastic syndrome, anemia, HIV, end-stage kidney disease, dialysis, cirrhosis, multiple myeloma, lymphocytic leukemia, myeloid leukemia, polycythaemia vera, haemochromatosis). Additionally, we excluded outlying genotype samples based on heterozygosity and/or rate of missing genotypes as defined by UK Biobank (data field 22027), and we removed any individuals having at least one relative in the cohort based on UK Biobank kinship calculations (samples with a value other than zero in data field 22021). Finally, to limit confounding due to population structure, we included only genotype samples marked as “White British” (based on a principal components analysis of the genotypes [38] stored in data field 22009). After filtering genotype samples according to these criteria, 257,605 samples remained.

We applied quantile normalization to the 16 blood cell traits measured in the 257,605 samples, separately for each trait, to transform each trait to the standard normal distribution. Since ultimately we aimed to jointly model the 16 blood cell traits, we removed outlying phenotypes according to a simple multivariate normal distribution of the phenotypes. Specifically, after quantile normalization, we measured the Mahalanobis distance ***y***^⊤^**Σ**c^™1^***y***_*i*_ for each individual *i*, where ***y***_*i*_ is the vector of 16 blood cell traits measured in individual *i*, and **Σ**c is the sample covariance matrix estimated from the 257,605 UK Biobank samples. We discarded samples with Mahalanobis distance falling within the [0.99, 1] quantile of the chi-square distribution with 16 degrees of freedom. This step removed 8,625 samples, for a final total of 248,980 UK Biobank samples.

Base-pair positions of the SNPs are reported using Genome Reference Consortium human genome assembly 37 (hg19).

### Association analyses of UK Biobank blood cell traits

Using the UK Biobank genotype and phenotype data prepared as described above, we computed association statistics for each of the 16 blood cell traits and for all available biallelic SNPs on autosomal chromosomes meeting the following criteria: minor allele frequency of 0.1% or greater; information (“INFO”) score of 0.6 or greater (the INFO score quantifies imputation quality). The same criteria were used in [60] to filter the SNPs.

Association statistics were computed using the `--glm` function in PLINK (version 2.00a2LM, 64-bit Intel, Feb 21, 2009) [43] with `hide-covar no-x-sex omit-ref –vif 100`. Following [1, 42], we included the following covariates in the association analyses: sex (data field 31), age at recruitment (21022), age × age, assessment center (54), and genotype measurement batch (22000). To limit inflation of spurious associations due to population structure, we also included the top 10 genotype PCs as covariates following previous association analyses of UK Biobank data (e.g., [61]). (These PCs were previously computed by UK Biobank [38] and stored in data field 22009.) The covariates input file for PLINK was prepared by calling the `model.matrix` function in R and standardizing quantitative covariates (age, PCs) to have mean 0 and variance 1.

The summary data provided as input to SuSiE and mvSuSiE were the *z*-scores and *p*-values extracted from the `T_STAT` and `P` columns in the `plink2 --glm` outputs. The association statistics computed using PLINK have been made available in a Zenodo repository (see “Data availability”).

### Selection of regions for fine-mapping

To select regions for fine-mapping, we adapted the approach used in [42] to the multivariate setting. In brief, we began by identifying regions separately for each trait. For each significant association (PLINK two-sided *t*-test *p*-value less than 5 × 10^™8^), we defined the fine-mapping region as all SNPs within ±250 kb of the significant association. Next, any regions overlapping by one or more SNPs were combined into a larger region. We repeated combining regions until no regions overlapped. This resulted in a set of fine-mapping regions for each of the 16 blood cell traits, similar to [42]. To form a single set of fine-mapping regions for all 16 traits, we then merged two regions from different traits whenever they overlapped. The end result of this procedure was a set 975 of disjoint fine-mapping regions satisfying the following two properties: (i) all significant SNPs (with PLINK *p*-value for two-sided *t*-test less than 5 × 10^™8^) belong to exactly one region; and (ii) all SNPs within 250 kb of a significant SNP belong to exactly one region. This procedure generated fine-mapping regions that varied considerably in size: their lengths ranged from 411 kb to 8.73 Mb (average size: 961 kb; median size: 686 kb); and the number of SNPs ranged from 93 SNPs to 36,605 SNPs (average number of SNPs: 4,776; median number of SNPs: 3,514). A listing of all 975 regions is given in Supplementary Table 4. These same regions were used in both the single-trait and multi-trait fine-mapping.

Note that we did not finemap the extended MHC [36] (defined as base-pair positions 25–36 Mb on chromosome 6). The MHC is particularly challenging to analyze and interpret, and therefore is typically analyzed separately [37, 38, 39].

### Simulations using UK Biobank genotypes

We evaluated the fine-mapping methods on data sets generated using real genotypes ***X*** and simulated phenotypes ***Y***. For the genotypes, we used the UK Biobank imputed genotypes. We simulated ***Y*** from different mvSuSiE models (see below). The genotype data were curated following the data preparation steps described above, so *N* = 248,980 in all our simulations. (To clarify, these data preparation steps included removing outlying blood cell trait observations. Even though this particular filtering step was not needed since we did not use the UK Biobank phenotype data in the simulations, for convenience we used the data prepared with this filtering step.)

### Simulation scenarios

We implemented three fine-mapping scenarios in the simulations.

In the simplest simulations, which we used to compare all of the methods (SuSiE, mvSuSiE, CAFEH, PAINTOR and flashfm), we simulated 2 traits under a variety of conditions: (i) independent traits with independent effects; (ii) independent traits with correlated effects; and (iii) correlated traits with independent effects. This simpler scenario was intended mainly for comparisons with PAINTOR and flashfm so as to not unfairly disadvantage these methods; flashfm cannot handle a large number of traits, and PAINTOR cannot handle a large number of causal SNPs, and assumes independent traits and independent effects (Table 1). However, for completeness we also compared with SuSiE and CAFEH in this simulation scenario.

For comparing other fine-mapping methods (mvSuSiE, SuSiE, CAFEH), we simulated data sets under two more complex scenarios, which we refer to as “Scenario A” and “Scenario B”.

In Scenario A, we simulated 20 independent traits in which the SNP effects were either specific to one trait or shared among traits in simple ways (equal effects among 2 traits, equal effects among half of the traits, or correlated equally among all 20 traits). In the results, we call Scenario A the “Trait-specific + Shared Effects” scenario.

Scenario B was intended to capture a combination of factors that one might more realistically encounter in fine-mapping studies. It is also more challenging because the traits are correlated and the effects are shared among the traits in complex ways. Specifically, we simulated using a residual covariance matrix ***V*** and sharing patterns ***U***_*k*_ obtained from our analyses of the UK Biobank blood cell traits. In the results, we refer to Scenario B as the “Complex Shared Effects” scenario.

### Simulation procedure

Let ***X*** denote the *N* × *J* genotype matrix for a given fine-mapping region, where *J* is the number of SNPs in the region, and *N* = 248,980. The procedure we used to simulate an *N* × *R* matrix ***Y*** was the following.

1. Center and scale the columns of ***X*** so that each column has a mean of 0 and a variance of 1.
2. Choose *S*, the number of causal SNPs. For Scenarios A and B, set *S* to 1, 2, 3, 4 or 5 with probabilities 0.3, 0.3, 0.2, 0.1, 0.1, respectively. For the 2-trait simulations, set *S* = 2.
3. Sample the indices of the *S* causal SNPs uniformly at random from {1, …, *J*}. Denote the set of causal SNPs by 𝒞.
4. For each SNP *j* ∈ 𝒞, simulate the *R* effects, ***b***_j_ ∈ ℝ^*R*^, from the mixture of multivariate normals (5), in which 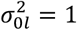. In the 2-trait simulations, we set 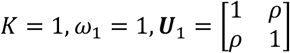,in which the correlation *ρ* between the two effects was 0, 0.5 or 1. We also simulated 2-trait data sets in which the effects were drawn from a mixture of *R* + 5 = 7 “canonical” covariance matrices (see the Supplementary Note). To draw each effect ***b***_j_ from this mixture, the mixture component probabilities were specified as follows: one of the 2 trait-specific covariances was chosen each with probability 0.2; or one of the remaining canonical covariances was chosen each with probability 0.12. In Scenario A, we simulated the effects of the causal SNPs ***b***_j_ using a mixture of 19 covariance matrices (Supplementary Fig. 4). For Scenario B, we simulated the effects using the mixture of 15 covariance matrices estimated from the UK Biobank data (Supplementary Fig. 5).
5. For each SNP *j* ∉ 𝒞, set ***b***_j_ = **0**.
6. Choose the residual variance *σ*^2^. To set *σ*^2^ to a realistic value, we set *σ*^2^ so that the greatest proportion of variance in a trait explained by the SNPs was 0.05%, which roughly corresponds to the proportion of variance explained in the mvSuSiE fine-mapping analyses of the UK Biobank blood cell traits. In particular, we solved for *σ*^2^ satisfying 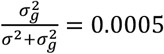, where 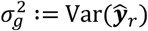 is the variance in the *r*th trait explained by the SNPs, in which Var(***θ***) denotes the sample variance, 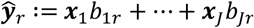, and 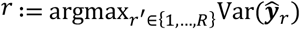.
7. Specify the *R* × *R* residual correlation matrix ***C***, then set ***V*** = *σ*^2^***C***. For Scenario A and the 2-trait scenario, ***C*** = ***I***_*R*_. For Scenario B, the 16 × 16 covariance matrix ***C*** was set to the correlation matrix estimated from the 16 blood cell traits after removing the linear effects of covariates (Supplementary Table 8). (Note that, although this correlation matrix was estimated from the UK Biobank data, for the simulations the ***V*** used in the mvSuSiE analyses of the simulated data was estimated using the simulated data and so the ***V*** used by mvSuSiE differed from the ***V*** used to simulate ***Y***.) In the 2-trait simulations, we set 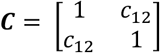, in which the correlation *c*_12_ between the two effects was 0, 0.4 or 0.8.
8. Simulate ***Y*** using (1).
9. Center and scale the columns of ***Y*** so that each column has a mean of zero and a variance of 1.
10. Compute the summary statistics—effect estimates 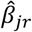, standard errors *ŝ*_*jr*_, *z*-scores *z*_*jr*_ and the in-sample LD matrix ***R***—using PLINK [43] and LDstore [66]. For these summary statistics, we extracted the `BETA`, `SE`, `T_STAT` and `P` columns from the `plink2 --glm` output (see above for more details on how PLINK was called). Note PLINK was applied to the raw genotypes without centering or scaling. We computed the *J* × *J* in-sample LD matrix 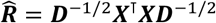, where ***D*** ≔ diag(***X***^⊤^***X***), using LDstore version 1.1.

We simulated data sets from the curated set of 975 regions for the UK Biobank blood cell traits (Supplementary Table 4). All selected regions had at least 1,000 SNPs and no more than 5,000 SNPs, and were at least 400 kb in size and at most 1.6 Mb.

These simulations produced an empirical distribution of *z*-scores roughly similar to the *z*-scores seen in association analyses of the blood cell traits; in our simulations, the largest *z*-score magnitude in each fine-mapping region had a median of 11.05, mean of 10.97 and a third quantile of 11.71, whereas the corresponding statistics for the UK Biobank blood cell traits were 8.01, 10.85 and 12.18.

### Details of the methods compared

In this section we describe how we ran the methods on the simulated data sets. SuSiE, mvSuSiE and PAINTOR were run using the *z*-scores and the in-sample LD. CAFEH and flashfm were run using the effect estimates, standard errors of these effect estimates, and in-sample LD. Some methods, including mvSuSiE, also accepted an additional input, the sample size (*N*), in which case we provided this as well. flashfm also required the reference allele frequencies, which in all our analyses were the minor allele frequencies.

### PAINTOR

We ran PAINTOR [19] in only the 2-trait simulations. PAINTOR was designed to work with functional genomic annotation data, so to run PAINTOR we created a single “dummy” annotation in which all SNPs were assigned to this annotation (that is, all entries of the annotation matrix were set to 1). For all data sets, we asked PAINTOR to enumerate all possible configurations up to 2 causal SNPs. (In the 2-trait simulations, the true number of causal SNPs was always 2.) We did not use the “mcmc” option (`-mcmc`) because the outputted PIPs when using this option were all zero in our tests. (The same issue was reported in https://github.com/gkichaev/PAINTOR_V3.0/issues/5.) All other PAINTOR options were kept at their default settings. Note that PAINTOR does not accept *N* (the sample size) as input. Also note that PAINTOR assumes that both traits and effects are independent across traits (Table 1).

### Flashfm

We ran flashfm [19] in only the 2-trait simulations. We ran flashfm by calling function `FLASHFMwithFINEMAP` from R package `flashfm` (version 0.0.0.9000). This function internally calls FINEMAP [10] (version 1.4.1) with settings `--sss --n-configs-top 1000 --n-causal-snps 10`, which allows configurations of up to 10 causal SNPs. We ran flashfm with 4 CPUs (`NCORES = 4`). All other flashfm settings were kept at their defaults. The inputs to `FLASHFMwithFINEMAP` were the effect estimates, the standard errors of these effect estimates, minor allele frequencies, vector of trait means, and sample size *N*. Since ***Y*** was centered and standardized in the simulations, the vector of trait means was simply a vector of zeros of length *R*.

### CAFEH

We used the `fit_cafeh_summary` interface in CAFEH 1.0 [43] installed with Python 3.7.4. The `fit_cafeh_summary` function accepts the following data inputs: effect estimates, standard errors of those estimates, LD matrix, and sample size *N*. When calling `fit_cafeh_summary`, all optional arguments were kept at the software defaults. CAFEH’s default setting for the upper limit on the number of single effects (“*K*” in the CAFEH model) is 10, which is the same default in SuSiE and mvSuSiE. Note that CAFEH assumes that traits and effects are independent across traits (Table 1). CAFEH outputs credible sets without any filter on the purity of the CSs. Therefore, to make the CAFEH credible sets comparable to SuSiE and mvSuSiE credible sets, we filtered out CSs with purity less than 0.5. For assessing performance of CAFEH PIPs and trait-wise PIPs (in CAFEH, these are called “study PIPs”), we called `get_pip` and `get_study_pip`.

Note that the two CAFEH summary data interfaces—`fit_cafeh_summary` and fit_cafeh_z—produce the same or very similar results when ***X*** is standardized. Both functions internally call function `CAFEHSummary` with the same LD matrix, but provide different effect estimates and standard errors of the effect estimates. Let 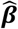 denote the vector of effect estimates (with one entry per SNP) and let ***ŝ*** denote the vector of standard errors (also with one entry per SNP). If `fit_cafeh_summary` calls `CAFEHSummary` with inputs 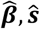, and assuming ***X*** is standardized, then it can be shown that `fit_cafeh_z` calls `CAFEHSummary` with inputs 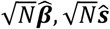. Since `CAFEHSummary` is invariant to rescaling of 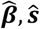—that is, `CAFEHSummary` generates the same result with inputs 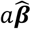 and 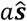 for any choice of scalar *a* > 0—it follows that `fit_cafeh_summary` and `fit_cafeh_z` also produce the same result when ***X*** is standardized. In practice, this invariance does not hold exactly since it requires that the prior on the effects also be appropriately rescaled, but empirically we have found that the CAFEH PIPs and posterior effect estimates are almost the same for different choices of *a* > 0. (See https://github.com/karltayeb/cafeh/blob/current_working_branch/notebooks/CAFEHS_scale_invariance.ipynb.)

### SuSiE

We ran SuSiE by calling function `susie_rss` from `susieR` [13] (version 0.12.12). In each data set, we ran `susie_rss` once per trait. The `susie_rss` interface accepts different types of summary data; we provided *z*-scores, in-sample LD, and sample size *N*. For all simulations, we set *L*, the maximum number of non-zero effects, to 10. (We also set *L* = 10 for the 2-trait simulations even though there were never more than 2 causal SNPs in these simulations.) We estimated the residual variance (`estimate_residual_variance = TRUE`), which is the recommended setting when the LD is estimated from the “in-sample” data. We set the maximum number of IBSS iterations to 1,000 (`max_iter = 1000`). The remaining optional arguments were kept at their defaults.

Since SuSiE analyzes each trait separately, it does not directly give evidence for a SNP being a cross-trait causal SNP. To quantify performance in this task and compare with mvSuSiE, we quantified the evidence for a cross-trait causal SNP using an *ad hoc* metric, the “maximum PIP”, defined as

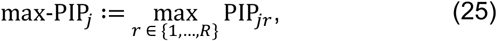

where PIP_*jr*_ is the PIP for SNP *j* obtained from the SuSiE analysis of trait *r*.

### mvSuSiE

We ran mvSuSiE using the `mvsusie_rss` interface from `mvsusieR` (version 0.0.3.0518, git commit id 9f28916). While `susie_rss` accepts a *vector* of *z*-scores, `mvsusie_rss` accepts a *matrix* of *z*-scores (specifically, a *J* × *R* matrix). In the simulations, we compared several mvSuSiE variants using different prior choices; for more details, see the Supplementary Note. (In the 2-trait simulations, we only used the canonical prior. This was a mixture of multivariate normals with *K* = 7 components.) We also compared mvSuSiE with different settings of the residual covariance ***V***; see the Supplementary Note. In all cases, we ran `mvsusie_rss` with the following settings: `L = 10`, `max_iter = 1000`, `estimate_prior_variance = TRUE`, `estimate_prior_method = “EM”, precompute_covariances = TRUE` and `n_thread = 4`. (We set *L* = 10 for the 2-trait simulations even though there were never more than 2 causal SNPs in these simulations.) All other options were kept at the default settings.

### Computing environment

All analyses of the simulated data sets were run on Linux machines (Scientific Linux 7.4) with 4 Intel Xeon E5-2680v4 (“Broadwell”) processors, and with R 4.1.0 linked to the OpenBLAS 0.3.13 optimized numerical libraries. At most 10 GB of memory was needed to perform a fine-mapping analysis of a single simulated data set using one of the methods. We used DSC version 0.4.3.5 to perform the simulations.

### Fine-mapping of UK Biobank blood cell traits using SuSiE and mvSuSiE

We fit a mvSuSiE model to each fine-mapping data set—specifically, the *z*-scores matrix ***Z*** and LD matrix ***R***—by calling `mvsusie_rss` from the mvsusieR package with the following settings: `L = 10, N = 248980, precompute_covariances = TRUE, estimate_prior_variance = TRUE, estimate_prior_method = “EM”, max_iter = 1000 and n_thread = 1. We ran SuSiE on` ***Z, R*** separately for each trait (*i*.*e*., column of ***Z***) by calling `susie_rss` with the following settings: `n = 248980, L = 10, max_iter = 1000, estimate_prior_variance` `= TRUE`, `refine = TRUE`. Any CSs returned by `susie_rss` or `mvsusie_rss` with purity less than 0.5 were removed.

### Enrichment analysis of regulatory annotations using GREGOR

We performed enrichment analyses of the SuSiE and mvSuSiE blood cell trait fine-mapping results using GREGOR [67] (version 1.4.0). In brief, GREGOR performs an enrichment analysis for a “positive set” of SNPs by calculating overlap with the given regulatory annotation, then estimates the probability of the observed overlap against its expectation using a set of “matched control SNPs”. We ran GREGOR with the following settings: `pop = ‘EUR’, r2_threshold = 0.7`, `ld_window_size = 10000`, `min_neighbor = 10`, `job_number = 10`.

Although GREGOR provides *p*-values, we found some issues with these *p*-values (e.g., some exceeded 1). Therefore, for each annotation, we extracted the intermediate GREGOR outputs to get a 2 × 2 table of the SNP counts of inside and outside the annotation intersected with positive set and matched control set. We then used this 2 × 2 table to perform Fisher’s exact test and this was the final *p*-value reported. Additional details about the GREGOR analysis can be found in the `20231106_GREGOR_functional_enrichment.ipynb` Jupyter notebook in of the Zenodo repositories.

We assessed enrichment for a total of 19 regulatory genomic annotations, including: enhancer promoter regions and transcription factor binding sites [68]; genomic structural elements [69]; eQTLs in multiple tissues (based on different false discovery rates) [70]; RNA polymerase II binding in EPC-treated HESCs [71]; and binding intervals for specific transcription factors. The specific transcription factors included were promyelocytic leukemia zinc finger protein (PLZF), *FOSL2, NR2F2 and FOXO1* [71]. The BED annotation files are included in one of the Zenodo repositories.

Using these regulatory genomic annotations, we performed two sets of GREGOR enrichment analyses, one using the SuSiE fine-mapping results, and another set using the mvSuSiE fine-mapping results. We performed these enrichment analyses separately for the fine-mapping results for each blood cell trait, as well as the “global” (cross-trait) results. For mvSuSiE, we included a SNP in the cross-trait positive set if the SNP was included in at least one 95% CS and/or the global PIP was greater than 0.7. We included a SNP in the positive set for a given blood cell trait if the SNP was included in at least one 95% CS and the *lfsr* for the given trait was less 0.01.

For SuSiE, we included a SNP in the positive set for a given trait if the SNP was included in at least one 95% CS and/or the PIP was greater than 0.7. The SuSiE cross-trait positive set was defined as the union of the 16 positive sets from the SuSiE analyses of each of the 16 traits.

### Enrichment analysis of hematopoietic cell-types using gchromVAR

We performed additional enrichment analyses of the SuSiE and mvSuSiE blood cell trait fine-mapping results using gchromVAR [41]. In brief, gchromVAR assesses overlap of the fine-mapped SNPs and regions of accessible chromatin, separately in different hematopoietic cell types. (This is actually a weighted overlap in which we have defined the weights as the SuSiE or mvSuSiE PIPs.) Each enrichment analysis was performed following the steps described in the gchromVAR R package vignette (we used version 0.3.2 of the gchromVAR R package). The *z*-scores returned by the `computeWeightedDeviations` function were then refined using adaptive shrinkage implemented in `ashr` [34] (version 2.2-57). The adaptive shrinkage posterior *z*-scores (posterior means divided by posterior standard deviations) and *lfsr* values were used to report the final enrichment results.

Similar to the GREGOR enrichment analyses, we performed two separate enrichment analyses with gchromVAR, one using the SuSiE results, and another using the mvSuSiE results. For SuSiE, we included a SNP for a given trait if the PIP > 0.01 for that trait. A total of 100,090 SNPs had PIP > 0.01 in at least one of the blood cell traits. For mvSuSiE, we included a SNP for a given trait if the global PIP > 0.01 and if the CS was significant for the given trait (*lfsr* < 0.01). A total of 39,884 SNPs had a global PIP > 0.01.

## Data availability

UK Biobank data, https://www.ukbiobank.ac.uk; PLINK association test statistics from the UK Biobank blood cell traits, https://doi.org/10.5281/zenodo.8088040.

## Code availability

The mvsusieR R package implementing our methods is available on GitHub at https://github.com/stephenslab/mvsusieR; mvsusieR 0.1.8 is also available via Zenodo at https://doi.org/10.5281/zenodo.17296669. Additional code resources accompanying this paper include: code for preparing the UK Biobank data, https://doi.org/10.5281/zenodo.8400278; code for performing the fine-mapping simulations, https://doi.org/10.5281/zenodo.8087907; additional code for the fine-mapping simulations and the fine-mapping analyses of the UK Biobank blood cell traits, https://doi.org/10.5281/zenodo.8094982. Other R packages and software used in this work include: susieR 0.12.12, https://github.com/stephenslab/susieR; mashr 0.2.59, https://github.com/stephenslab/mashr; flashr 0.6-8, https://github.com/stephenslab/flashr; PAINTOR 3.1, https://github.com/gkichaev/PAINTOR_V3.0; BayesSUR 2.0-1, https://cran.r-project.org/package=BayesSUR; flashfm 0.0.0.9000, https://github.com/jennasimit/flashfm; FINEMAP 1.4.1, http://www.christianbenner.com; msCAVIAR 0.1, https://github.com/nlapier2/MsCAVIAR; CAFEH 1.0, https://github.com/karltayeb/cafeh; hyprcoloc 1.0, https://github.com/cnfoley/hyprcoloc; moloc 0.1.0, https://bogdan.dgsom.ucla.edu/pages/MOLOC; MFM 0.2-1, https://github.com/jennasimit/MFM; gchromVAR 0.3.2, https://github.com/caleblareau/gchromVAR; GREGOR 1.4.0, http://csg.sph.umich.edu/GREGOR/; R 4.1.0, https://cran.r-project.org; Python 3.7.4, https://www.python.org; PLINK 2.00a2LM, 64-581 bit Intel, Feb 21, 2009, https://www.cog-genomics.org/plink2; LDstore 1.1, http://christianbenner.com; DSC 0.4.3.5, https://github.com/stephenslab/dsc.

## References

[1] O. Canela-Xandri, K. Rawlik and A. Tenesa. An atlas of genetic associations in UK Biobank. Nature Genetics 50, 1593–1599 (2018).

[2] P. M. Visscher, N. R. Wray, Q. Zhang, P. Sklar, M. I. McCarthy, M. A. Brown and J. Yang, 10 Years of GWAS Discovery: biology, function, and translation. American Journal of Human Genetics 101, 5–22 (2017).

[3] A. Buniello, J. A. L. MacArthur, M. Cerezo, L. W. Harris, J. Hayhurst and others. The NHGRI-EBI GWAS Catalog of published genome-wide association studies, targeted arrays and summary statistics 2019. Nucleic Acids Research 47, D1005–D1012 (2018).

[4] V. Tam, N. Patel, M. Turcotte, Y. Bossé, G. Paré and D. Meyre. Benefits and limitations of genome-wide association studies. Nature Reviews Genetics 20, 467–484 (2019).

[5] F. Hormozdiari, E. Kostem, E. Y. Kang, B. Pasaniuc and E. Eskin. Identifying causal variants at loci with multiple signals of association. Genetics 198, 497–508 (2014).

[6] G. Kichaev, W.-Y. Yang, S. Lindstrom, F. Hormozdiari, E. Eskin, A. L. Price, P. Kraft and B. Pasaniuc. Integrating functional data to prioritize causal variants in statistical fine-mapping studies. PLoS Genetics 10, e1004722 (2014).

[7] J. B. Maller, G. McVean, J. Byrnes, D. Vukcevic, K. Palin and others. Bayesian refinement of association signals for 14 loci in 3 common diseases. Nature Genetics 44, 1294–1301 (2012).

[8] J. Yang, T. Ferreira, A. P. Morris, S. E. Medland, P. A. F. Madden, A. C. Heath, N. G. Martin, G. W. Montgomery, M. N. Weedon, R. J. Loos, T. M. Frayling, M. I. McCarthy, J. N. Hirschhorn, M. E. Goddard, P. M. Visscher, GIANT Consortium and DIAGRAM Consortium. Conditional and joint multiple-SNP analysis of GWAS summary statistics identifies additional variants influencing complex traits. Nature Genetics 44, 369–375 (2012).

[9] W. Chen, B. R. Larrabee, I. G. Ovsyannikova, R. B. Kennedy, I. H. Haralambieva, G. A. Poland and D. J. Schaid. Fine mapping causal variants with an approximate Bayesian method using marginal test statistics. Genetics 200, 719–736 (2015).

[10] C. Benner, C. C. A. Spencer, A. S. Havulinna, V. Salomaa, S. Ripatti and M. Pirinen, FINEMAP: efficient variable selection using summary data from genome-wide association studies. Bioinformatics 32, 1493–1501 (2016).

[11] X. Wen, Y. Lee, F. Luca and R. Pique-Regi. Efficient integrative multi-SNP association analysis via deterministic approximation of posteriors. American Journal of Human Genetics 98, 1114–1129 (2016).

[12] Y. Lee, L. Francesca, R. Pique-Regi and X. Wen, Bayesian multi-SNP genetic association analysis: control of FDR and use of summary statistics. bioRxiv doi:10.1101/316471 (2018).

[13] G. Wang, A. Sarkar, P. Carbonetto and M. Stephens. A simple new approach to variable selection in regression, with application to genetic fine mapping. Journal of the Royal Statistical Society, Series B 82, 1273–1300 (2020).

[14] C. Wallace, A. J. Cutler, N. Pontikos, M. L. Pekalski, O. S. Burren, J. D. Cooper, A. R. García, R. C. Ferreira, H. Guo, N. M. Walker, D. J. Smyth, S. S. Rich, S. Onengut-Gumuscu, S. J. Sawcer, M. Ban, S. Richardson, J. A. Todd and L. S. Wicker. Dissection of a complex disease susceptibility region using a Bayesian stochastic search approach to fine mapping. PLoS Genetics 11, e1005272 (2015).

[15] D. J. Schaid, W. Chen and N. B. Larson, From genome-wide associations to candidate causal variants by statistical fine-mapping. Nature Reviews Genetics 19, 491–504 (2018).

[16] Y. Zou, P. Carbonetto, G. Wang and M. Stephens. Fine-mapping from summary data with the “Sum of Single Effects” model. PLoS Genetics 18, e1010299 (2022).

[17] M. Stephens. A unified framework for association analysis with multiple related phenotypes. PLoS ONE 8, e65245 (2013).

[18] A. Lewin, H. Saadi, J. E. Peters, A. Moreno-Moral, J. C. Lee, K. G. C. Smith, E. Petretto, L. Bottolo and S. Richardson. MT-HESS: an efficient Bayesian approach for simultaneous association detection in OMICS datasets, with application to eQTL mapping in multiple tissues. Bioinformatics 32, 523–532 (2016).

[19] G. Kichaev, M. Roytman, R. Johnson, E. Eskin, S. Lindstroem, P. Kraft and B. Pasaniuc. Improved methods for multi-trait fine mapping of pleiotropic risk loci. Bioinformatics 33, 248–255 (2017).

[20] N. Hernández, J. Soenksen, P. Newcombe, M. Sandhu, I. Barroso, C. Wallace and J. L. Asimit. The flashfm approach for fine-mapping multiple quantitative traits. Nature Communications 12, 6147 (2021).

[21] Z. Zhao, M. Banterle, L. Bottolo, S. Richardson, A. Lewin and M. Zucknick. BayesSUR: an R Package for high-dimensional multivariate Bayesian variable and covariance selection in linear regression. Journal of Statistical Software 100, 1–32 (2021).

[22] N. LaPierre, K. Taraszka, H. Huang, R. He, F. Hormozdiari and E. Eskin. Identifying causal variants by fine mapping across multiple studies. PLoS Genetics 17, e1009733 (2021).

[23] M. Arvanitis, K. Tayeb, B. J. Strober and A. Battle. Redefining tissue specificity of genetic regulation of gene expression in the presence of allelic heterogeneity. American Journal of Human Genetics 109, 223–239 (2022).

[24] J. L. Asimit, D. B. Rainbow, M. D. Fortune, N. F. Grinberg, L. S. Wicker and C. Wallace, Stochastic search and joint fine-mapping increases accuracy and identifies previously unreported associations in immune-mediated diseases. Nature Communications 10, 3216 (2019).

[25] C. N. Foley, J. R. Staley, P. G. Breen, B. B. Sun, P. D. W. Kirk, S. Burgess and J. M. M. Howson. A fast and efficient colocalization algorithm for identifying shared genetic risk factors across multiple traits. Nature Communications 12, 764 (2021).

[26] C. Giambartolomei, J. Zhenli Liu, W. Zhang, M. Hauberg, H. Shi, J. Boocock, J. Pickrell, A. E. Jaffe, T. C. Consortium, B. Pasaniuc and P. Roussos. A Bayesian framework for multiple trait colocalization from summary association statistics. Bioinformatics 34, 2538–2545 (2018).

[27] C. Wallace. A more accurate method for colocalisation analysis allowing for multiple causal variants. PLoS Genetics, 17, e1009440 (2021).

[28] C. Giambartolomei, D. Vukcevic, E. E. Schadt, L. Franke, A. D. Hingorani, C. Wallace and V. Plagnol. Bayesian test for colocalisation between pairs of genetic association studies using summary statistics. PLoS Genetics 10, e1004383 (2014).

[29] F. Hormozdiari, M. Van De Bunt, A. V. Segre, X. Li, J. W. J. Joo, M. Bilow, J. H. Sul, S. Sankararaman, B. Pasaniuc and E. Eskin. Colocalization of GWAS and eQTL signals detects target genes. American Journal of Human Genetics 99, 1245–1260 (2016).

[30] X. Wen, R. Pique-Regi and F. Luca. Integrating molecular QTL data into genome-wide genetic association analysis: probabilistic assessment of enrichment and colocalization. PLoS Genetics 13, e1006646 (2017).

[31] S. M. Urbut, G. Wang, P. Carbonetto and M. Stephens. Flexible statistical methods for estimating and testing effects in genomic studies with multiple conditions. Nature Genetics 51, 187–195, 2019.

[32] M. Kanai, R. Elzur, W. Zhou, M. Kanai, K.-H. H. Wu and others. Meta-analysis fine-mapping is often miscalibrated at single-variant resolution. Cell Genomics 2, 100210 (2022).

[33] B. Pasaniuc and A. L. Price. Dissecting the genetics of complex traits using summary association statistics. Nature Reviews Genetics 18, 117–127 (2017).

[34] M. Stephens. False discovery rates: a new deal. Biostatistics 18, 275–294 (2017).

[35] J. Bovy, D. W. Hogg and S. T. Roweis. Extreme Deconvolution: inferring complete distribution functions from noisy, heterogeneous and incomplete observations. Annals of Applied Statistics 5, 1657–1677 (2011).

[36] Z. Zhao, M. Banterle, A. Lewin and M. Zucknick. Multivariate Bayesian structured variable selection for pharmacogenomic studies. Journal of the Royal Statistical Society, Series C 73, 420–443 (2024).

[37] L. Bottolo, M. Banterle, S. Richardson, M. Ala-Korpela, M.-R. Järvelin and A. Lewin. A computationally efficient Bayesian seemingly unrelated regressions model for high-dimensional quantitative trait loci discovery. Journal of the Royal Statistical Society, Series C, 70, 886–908 (2021).

[38] C. Bycroft, C. Freeman, D. Petkova, G. Band, L. T. Elliott, K. Sharp, A. Motyer, D. Vukcevic, O. Delaneau, J. O’Connell, A. Cortes, S. Welsh, A. Young, M. Effingham, G. McVean, S. Leslie, N. Allen, P. Donnelly and J. Marchini. The UK Biobank resource with deep phenotyping and genomic data. Nature 562, 203–209 (2018).

[39] W. J. Astle, H. Elding, T. Jiang, D. Allen, D. Ruklisa and others. The allelic landscape of human blood cell trait variation and links to common complex disease. Cell 167, 1415–1429 (2016).

[40] L. Kachuri, S. Jeon, A. T. DeWan, C. Metayer, X. Ma, J. S. Witte, C. W. K. Chiang, J. L. Wiemels and A. J. de Smith. Genetic determinants of blood-cell traits influence susceptibility to childhood acute lymphoblastic leukemia. American Journal of Human Genetics 108, 1823–1835 (2021).

[41] J. C. Ulirsch, C. A. Lareau, E. L. Bao, L. S. Ludwig, M. H. Guo, C. Benner, A. T. Satpathy, V. K. Kartha, R. M. Salem, J. N. Hirschhorn, H. K. Finucane, M. J. Aryee, J. D. Buenrostro and V. G. Sankaran. Interrogation of human hematopoiesis at single-cell and single-variant resolution. Nature Genetics 51, 683–693 (2019).

[42] D. Vuckovic, E. L. Bao, P. Akbari, C. A. Lareau, A. Mousas and others. The polygenic and monogenic basis of blood traits and diseases. Cell 182, 1214-1231.e11 (2020).

[43] C. C. Chang, C. C. Chow, L. C. A. M. Tellier, S. Vattikuti, S. M. Purcell and J. J. Lee. Second-generation PLINK: rising to the challenge of larger and richer datasets. Gigascience 4, s13742.–015–0047–8 (2015).

[44] S. Tokuhiro, R. Yamada, X. Chang, A. Suzuki, Y. Kochi, T. Sawada, M. Suzuki, M. Nagasaki, M. Ohtsuki, M. Ono, H. Furukawa, M. Nagashima, S. Yoshino, A. Mabuchi, A. Sekine, S. Saito, A. Takahashi, T. Tsunoda, Y. Nakamura and K. Yamamoto. An intronic SNP in a RUNX1 binding site of SLC22A4, encoding an organic cation transporter, is associated with rheumatoid arthritis. Nature Genetics 35, 341–348 (2003).

[45] C. J. F. Scheitz and T. Tumbar. New insights into the role of Runx1 in epithelial stem cell biology and pathology. Journal of Cellular Biochemistry 114, 985–993 (2013).

[46] K. Asano, S. Ueki, M. Tamari, Y. Imoto, S. Fujieda and M. Taniguchi. Adult-onset eosinophilic airway diseases. Allergy 75, 3087–3099 (2020).

[47] C. Helms, L. Cao, J. G. Krueger, E. M. Wijsman, F. Chamian, D. Gordon, M. Heffernan, J. A. W. Daw, J. Robarge, J. Ott, P.-Y. Kwok, A. Menter and A. M. Bowcock, “A putative RUNX1 binding site variant between SLC9A3R1 and NAT9 is associated with susceptibility to psoriasis. Nature Genetics 35, 349–356 (2003).

[48] S. L. Alper. Genetic diseases of PIEZO1 and PIEZO2 dysfunction. In Piezo Channels 79, P. A. Gottlieb, Ed., Academic Press, p. 97–134 (2017).

[49] S. M. Cahalan, V. Lukacs, S. S. Ranade, S. Chien, M. Bandell and A. Patapoutian, “Piezo1 links mechanical forces to red blood cell volume. eLife 4, e07370 (2015).

[50] T. Ling and J. D. Crispino. GATA1 mutations in red cell disorders. IUBMB Life 72, 106–118 (2020).

[51] S. Ma, A. E. Dubin, Y. Zhang, S. A. R. Mousavi, Y. Wang, A. M. Coombs, M. Loud, I. Andolfo and A. Patapoutian. A role of PIEZO1 in iron metabolism in mice and humans. Cell 184, 969–982.e13 (2021).

[52] K. E. Nichols, J. D. Crispino, M. Poncz, J. G. White, S. H. Orkin, J. M. Maris and M. J. Weiss. Familial dyserythropoietic anaemia and thrombocytopenia due to an inherited mutation in GATA1. Nature Genetics 24, 266–270 (2000).

[53] B. Han and E. Eskin. Random-effects model aimed at discovering associations in metaanalysis of genome-wide association studies. American Journal of Human Genetics 88, 586–598 (2011).

## Methods-only references

[54] A. P. Dawid. Some matrix-variate distribution theory: notational considerations and a Bayesian application. Biometrika 68, 265–274 (1981).

[55] A. K. Gupta and D. K. Nagar. Matrix variate distributions. Boca, Raton: Chapman & Hall, 2000.

[56] E. I. George and R. E. McCulloch. Approaches for Bayesian variable selection. Statistica Sinica 7, 339–373 (1997).

[57] J. Schwartzentruber, S. Cooper, J. Z. Liu, I. Barrio-Hernandez, E. Bello, N. Kumasaka, A. M. H. Young, R. J. M. Franklin, T. Johnson, K. Estrada, D. J. Gaffney, P. Beltrao and A. Bassett. Genome-wide meta-analysis, fine-mapping and integrative prioritization implicate new Alzheimer’s disease risk genes. Nature Genetics 53, 392–402 (2021).

[58] C. Sudlow, J. Gallacher, N. Allen, V. Beral, P. Burton, J. Danesh, P. Downey, P. Elliott, J. Green, M. Landray, B. Liu, P. Matthews, G. Ong, J. Pell, A. Silman, A. Young, T. Sprosen, T. Peakman and R. Collins. UK Biobank: an open access resource for identifying the causes of a wide range of complex diseases of middle and old age. PLoS Medicine 12, e1001779 (2015).

[59] S. M. Sheard, R. Nicholls and J. Froggatt. UK Biobank haematology data companion document, 2017.

[60] O. Weissbrod, F. Hormozdiari, C. Benner, R. Cui, J. Ulirsch, S. Gazal, A. P. Schoech, B. van de Geijn, Y. Reshef, C. Márquez-Luna, L. O’Connor, M. Pirinen, H. K. Finucane and A. L. Price. Functionally informed fine-mapping and polygenic localization of complex trait heritability. Nature Genetics 52, 1355–1363 (2020).

[61] J. Mbatchou, L. Barnard, J. Backman, A. Marcketta, J. A. Kosmicki, A. Ziyatdinov, C. Benner, C. O’Dushlaine, M. Barber, B. Boutkov, L. Habegger, M. Ferreira, A. Baras, J. Reid, G. Abecasis, E. Maxwell and J. Marchini. Computationally efficient whole-genome regression for quantitative and binary traits. Nature Genetics 53, 1097–1103 (2021).

[62] R. Horton, L. Wilming, V. Rand, R. C. Lovering, E. A. Bruford, V. K. Khodiyar, M. J. Lush, S. Povey, C. C. Talbot, M. W. Wright, H. M. Wain, J. Trowsdale, A. Ziegler and S. Beck. Gene map of the extended human MHC. Nature Reviews Genetics 5, 889–899 (2004).

[63] M. Caliskan, C. D. Brown and J. C. Maranville. A catalog of GWAS fine-mapping efforts in autoimmune disease. American Journal of Human Genetics 108, p. 549–563 (2021).

[64] V. Matzaraki, V. Kumar, C. Wijmenga and A. Zhernakova, “The MHC locus and genetic susceptibility to autoimmune and infectious diseases. Genome Biology 18, 76 (2017).

[65] S. Raychaudhuri, C. Sandor, E. a. Stahl, J. Freudenberg, H.-S. Lee, X. Jia, L. Alfredsson, L. Padyukov, L. Klareskog, J. Worthington, K. a. Siminovitch, S.-C. Bae, R. M. Plenge, P. K. Gregersen and P. I. W. de Bakker. Five amino acids in three HLA proteins explain most of the association between MHC and seropositive rheumatoid arthritis. Nature Genetics 44, 291–296 (2012).

[66] C. Benner, A. S. Havulinna, M.-R. Järvelin, V. Salomaa, S. Ripatti and M. Pirinen. Prospects of fine-mapping trait-associated genomic regions by using summary statistics from genome-wide association studies. American Journal of Human Genetics 101, 539–551 (2017).

[67] E. M. Schmidt, J. Zhang, W. Zhou, J. Chen, K. L. Mohlke, Y. E. Chen and C. J. Willer. GREGOR: evaluating global enrichment of trait-associated variants in epigenomic features using a systematic, data-driven approach. Bioinformatics 31, 2601–2606 (2015).

[68] H. K. Finucane, B. Bulik-Sullivan, A. Gusev, G. Trynka, Y. Reshef, P. R. Loh, V. Anttila, H. Xu, C. Zang, K. Farh, S. Ripke, F. R. Day, S. Purcell, E. Stahl, S. Lindstrom, J. R. B. Perry, Y. Okada, S. Raychaudhuri, M. J. Daly, N. Patterson, B. M. Neale and A. L. Price. Partitioning heritability by functional annotation using genome-wide association summary statistics. Nature Genetics 47, 1228–1235 (2015).

[69] A. Gusev, S. H. Lee, G. Trynka, H. Finucane, B. J. Vilhjálmsson, H. Xu, C. Zang, S. Ripke, B. Bulik-Sullivan, E. Stahl, A. K. Kähler, C. M. Hultman, S. M. Purcell, S. A. McCarroll, M. Daly, B. Pasaniuc, P. F. Sullivan, B. M. Neale, N. R. Wray, S. Raychaudhuri and A. L. Price. Partitioning heritability of regulatory and cell-type-specific cariants across 11 common diseases. American Journal of Human Genetics 95, 535–552 (2014).

[70] F. Hormozdiari, S. Gazal, B. van de Geijn, H. K. Finucane, C. J. T. Ju, P.-R. Loh, A. Schoech, Y. Reshef, X. Liu, L. O’Connor, A. Gusev, E. Eskin and A. L. Price. Leveraging molecular quantitative trait loci to understand the genetic architecture of diseases and complex traits. Nature Genetics 50, 1041–1047 (2018).

[71] Y. M. Vasquez, E. C. Mazur, X. Li, R. Kommagani, L. Jiang, R. Chen, R. B. Lanz, E. Kovanci, W. E. Gibbons and F. J. DeMayo. FOXO1 is required for binding of PR on IRF4, novel transcriptional regulator of endometrial stromal decidualization. Molecular Endocrinology 29, 421–433 (2015).

