## Supplementary Note and Figures for "Fast and flexible joint fine-mapping of multiple traits via the Sum of Single Effects model"

**Supplementary figures** 23

### Supplementary note

#### *More details about Table 1*

Here are more details about the columns in Table 1 in which we compared different statistical methods for multivariate fine-mapping:

1. Upper limit on number of causal SNPs: "User-specified" means that the method requires the user to specify an upper limit on the number of causal SNPs. "No limit" means that the method does not constrain the number of causal SNPs.

2. Data accepted, summary and sufficient. Methods marked "yes" under "sufficient" should reproduce the same result as if the methods were provided with the full data. Summary data approximate the summary statistics, and therefore are not expected to exactly reproduce the results that would be obtained with the full data. (All methods implicitly allow full data because summary statistics can always be computed from full data.) See [1] or below for definitions of "summary data" and "sufficient data".

3. CSs. Does the method compute credible sets (CS)? Note that calculation of CSs is straightforward when limiting to at most 1 causal SNP.

4. Allows correlated traits. Methods marked as "no" assume measurement error is independent across traits. Assuming independent errors is appropriate when non-overlapping samples were used in the different traits.

5. Models effect sharing. Methods with a "no" in this column assume (implicitly or explicitly) that the SNP effects on different traits are independent (conditioned on the SNP having a nonzero effect on one or more traits). A "yes" means that the method can model correlations among the effects on different traits.

6. Sample runtimes. Sample runtimes were obtained by running the software on a simulated data set with $J$ = 5,000 SNPs, $N$ = 250,000 individuals, and $R = 2$ or $R = 20$ traits. When the method accepted either full data or summary data, the summary-data version of the method was used. Note that the sample size, $N$, should only affect the running time for methods that only accept full data; it should not affect runtime of summary-data methods. When the method limits the number of causal SNPs, the upper limit was always set to 10 or the the largest acceptable value if this was less than 10. For PAINTOR, the upper limit was set to 2 because, in our tests, PAINTOR ran for a very long time when allowing 3 or more causal SNPs. See Methods for details about the computing environment used to obtain these runtimes.

7. Software and version. The name of the software and the version number of the software that was used in our evaluations. For mvSuSiE, the git commit id was given instead of the version number.

#### *Additional discussion of mvSuSiE: assumptions, limitations, and practical guidance*

Here we discuss some of mvSuSiE's limitations, and give practical guidance for applying mvSuSiE to other types of traits (e.g., binary traits) and for dealing with other complications not directly addressed in the paper (e.g., missing data).

*Modeling trait-specific sparsity.* The mvSuSiE model is "sparse" in that it assumes a small number of causal SNPs. However, the data-driven prior for the effect sizes at these causal SNPs will not generally be sparse. That is, each causal SNP is assumed to affect all traits (with some exceptions, such as when the prior includes sharing patterns reflecting an effect in one trait and no effects in the others). Instead of inducing sparsity on the effects of causal SNPs, we focussed on *estimating* these effects and assessing their significance by computing the *lfsr*s, building on the approach for analyzing multi-trait effects introduced in [2]. (Indeed, [2] contains other ideas for analyzing multi-trait effects that were not explored in this work, such as sharing of effects by both *sign* and *magnitude*, and these ideas could potentially be used to understand how the causal SNPs affect different traits.) This approach simplifies computation and worked well in our examples. Indeed, in additional simulation studies we found that mixture models constructed using our data-driven approach could capture the predominant sparsity patterns reasonably well (Supplementary Figures 6–7), and so mvSuSiE did not suffer from a loss of power to detect such sparse association signals. That being said, it is possible that explicitly modeling trait-specific sparsity of causal SNPs could be helpful in settings with large numbers of traits that are less related; with a large number of less related traits, the SNP effects may be shared primarily among small subsets of more related traits. This could perhaps be achieved by combining the mvSuSiE prior with indicator variables for each trait, similar to the strategy used in CAFEH.

*Fine-mapping binary traits and other types of traits.* mvSuSiE assumes a standard (Gaussian) multivariate linear model and so is most applicable to quantitative traits. However, provided the effect sizes are small, there is good theoretical and empirical justification for applying standard linear models directly to binary traits [3, 4]. Thus, in genetic association studies where individual SNP effects are small, it would be reasonable to apply mvSuSiE directly to studies where some or all of the traits are binary (e.g., disease status). When fine-mapping with summary data, this would mean applying mvSuSiE-RSS to the summary statistics from a *linear regression analysis* of the binary traits. While it may seem more intuitive to generate the summary statistics from a *logistic regression analysis* of the binary traits, there seems to be less theoretical or empirical support for this approach, even in the univariate case. More generally, there has been little theoretical or empirical assessment of univariate fine-mapping methods using summary data from linear mixed models or from generalized linear models, or for that matter any model other than a simple linear regression model; more work in this area seems important.

*Missing data.* The multivariate linear model in mvSuSiE assumes that the traits are measured in the same samples, with no missing data. When the traits involved are not independent—or, more precisely, when the residual error terms are not independent—correctly dealing with missing data in this model is complicated. Indeed, even maximum-likelihood estimation under a simple "missing-at-random" assumption becomes quite involved [5]. In cases with small amounts of missing data, we suggest imputing the missing values before fine-mapping. (This suggestion applies to both a full-data analysis and a summary-data analysis. Analyses of summary data computed with large or unknown amounts missingness should proceed with caution, if at all.) This "pre-imputation" approach may lose power compared with more rigorous approaches, but is more straightforward. If there are larger amounts of missing data, then it may be necessary to focus on a subset of samples with more overlap in available measurements. This could result in a tradeoff between sample size and number of traits analyzed; for example, with a large amount of missing data, it may be more powerful to analyze a subset of the traits at a larger sample size.

*Other types of analyses that are related to multi-trait fine-mapping.* Some multivariate association analyses involve traits measured in non-overlapping sets of individuals. Examples include colocalization of expression QTLs and GWAS traits in different samples [6]; multi-ancestry [7, 8]; and meta-analysis fine-mapping of multiple non-overlapping diseases with a common control set. Another important case is fine-mapping in which the summary statistics are generated from multiple studies ("meta-analysis fine-mapping") [9], as well as fine-mapping of multiple related traits from multiple studies ("multi-trait meta-analysis fine-mapping"). Formally, these analyses could all be implemented as extensions of mvSuSiE that allow for missing trait data. However, in practice these special cases yield important simplifications, and therefore it may be preferable to treat these special cases separately, with dedicated software implementations. Indeed, other groups have already successfully developed versions of SuSiE for some of these settings [10–12].

### Conventions for mathematical expressions

Here we summarize the notational conventions used. Matrices are written using bold, uppercase letters (e.g., $\boldsymbol{A}$), column vectors are written as bold, lowercase letters (e.g., $\boldsymbol{a}$), and scalars are written in plain font (e.g., $a$, $A$). For indexing, we usually use a capital letter to denote the total number of elements, and we use the corresponding lowercase symbol to denote the index; e.g., $j = 1, \ldots, J$. We use $\mathbb{R}$ to denote the real numbers and $\mathbb{R}^d$ for the set of real vectors of dimension $d$. We use $\Delta^d$ to denote the simplex on $\mathbb{R}^d$; that is, all $x \in \mathbb{R}^d$ such that $x_1 + \cdots + x_d = 1$, $x_i \geqslant 0$, $i = 1, \ldots, d$. We use $\mathbb{R}^{m \times n}$ to denote the set of all $m \times n$ matrices with real entries, $\mathbb{S}^n_{++}$ for the set of all $n \times n$ real, symmetric positive definite matrices, and $\mathbb{S}^n_{+}$ for the set of all $n \times n$ real, symmetric positive semi-definite matrices (this set may include matrices that are singular, or not invertible). We write the matrix transpose of $\boldsymbol{A}$ as $\boldsymbol{A}^\intercal$. For a square matrix $\boldsymbol{A}$, its inverse is $\boldsymbol{A}^{-1}$, its determinant is $\det \boldsymbol{A}$, also written as $|\boldsymbol{A}|$, the trace is $\mathrm{tr}(\boldsymbol{A})$, and $\boldsymbol{A}^\dagger$ denotes the Moore–Penrose inverse ("pseudoinverse") of $\boldsymbol{A}$. We use $\boldsymbol{I}_n$ for the $n \times n$ identity matrix, and we use $\mathbf{1}_n$ as a shorthand for a column vector of ones of length $n$. We use $\boldsymbol{a}^\intercal$ to denote a row vector. We denote the outer product of (column) vectors $\boldsymbol{a}$ and $\boldsymbol{b}$ as $\boldsymbol{a} \otimes \boldsymbol{b} := \boldsymbol{a}\boldsymbol{b}^\intercal$. Finally, we typically denote ordered or unordered sets using calligraphic letters (e.g., $\mathcal{A}$).

#### Posterior computation approach

Here we outline our approach to estimating the posterior distribution for the unknowns of primary interest, the single effect matrices $\boldsymbol{B}^{(1)}, \ldots, \boldsymbol{B}^{(L)}$, building on the ideas introduced in [13]. A more formal mathematical development of the mvSuSiE algorithms is given below. In this section, we assume that the model parameters $(\boldsymbol{V}, \boldsymbol{\pi}, \boldsymbol{\omega}, \mathcal{U})$ and $L$ are known, or have been estimated in earlier analysis steps. We also assume in this section that the scaling factors $\sigma_1^2, \ldots, \sigma_L^2$ are known (below we describe how the scaling factors are estimated).

The posterior distribution of $\boldsymbol{B}^{(1)}, \ldots, \boldsymbol{B}^{(L)}$, as in other Bayesian variable selection models, is intractable, and therefore we must resort to numerical approximations. A key consideration is that we would like these computations to scale well to large genetic data sets, which makes intensive Monte Carlo techniques such as Markov chain Monte Carlo [14–21] less attractive. Another key consideration is that we would like accurate estimates of posterior quantities which can be difficult to achieve when many variables (the SNPs) are highly correlated, or correlated in complicated ways, which is typically the case in genetic fine-mapping. These considerations, as well as others, motivated us to develop an alternative posterior computation approach for SuSiE based on variational approximation ideas [13]. The algorithm for performing the approximate posterior computations in SuSiE was called "Iterative Bayesian Stepwise Selection" (IBSS). In this paper, we have extended this approach to the mvSuSiE model. Therefore, extending the ideas developed in [13] leads to an IBSS algorithm for fitting the mvSuSiE model (this is Algorithm 1, given below).

#### Choice of L

The number of effects, $L$, is typically not known in advance. However, mvSuSiE is generally robust to misspecification of $L$ so long as $L$ is chosen to be larger than the (true) number of effects. That's because mvSuSiE prunes single effects when they are not needed by estimating the scaling factors $\sigma_{0l}^2$ in the prior (5) as zero or close to zero. This approach to estimating $L$ by adapting the prior is closely related to "automatic relevance determination" [22, 23], and this same approach was used in SuSiE [13].

#### Extension of posterior computation approach to work with summary data

The strategy used in [1] to extend SuSiE to summary data is quite general, and we take this same approach here. First, in we describe an algorithm that uses *sufficient statistics*, and produces the same result as running the mvSuSiE IBSS algorithm on the individual-level data. Second, we consider summary data that approximate the sufficient statistics, and therefore yield results that do not exactly reproduce mvSuSiE with individual-level data. Third, since many genetic association studies provide $z$-scores or other summary statistics that can be used to compute $z$-scores, we also focus on mvSuSiE-RSS with $z$-scores (see "Special case when $\boldsymbol{X}, \boldsymbol{Y}$ are standardized: mvSuSiE-RSS with $z$-scores").

*mvSuSiE with sufficient statistics.* The data enter the mvSuSiE model only through the likelihood, which from (2) is

$$\ell(\boldsymbol{B}; \boldsymbol{X}, \boldsymbol{Y}) = |2\pi\boldsymbol{V}|^{-N/2} \exp\big\{ -\tfrac{1}{2}\mathrm{tr}[\boldsymbol{V}^{-1}(\boldsymbol{Y}^\intercal\boldsymbol{Y} - 2\boldsymbol{B}^\intercal\boldsymbol{X}^\intercal\boldsymbol{Y} + \boldsymbol{B}^\intercal\boldsymbol{X}^\intercal\boldsymbol{X}\boldsymbol{B})]\big\}. \tag{6}$$

Here, we treat $\boldsymbol{V}$ as a fixed quantity so we do not explicitly mention this dependency in the notation for the likelihood. It is clear from this expression that the data influence the likelihood only through the quantities $\boldsymbol{X}^\intercal\boldsymbol{Y}$ and $\boldsymbol{X}^\intercal\boldsymbol{X}$. Therefore, $\boldsymbol{X}^\intercal\boldsymbol{Y}$ and $\boldsymbol{X}^\intercal\boldsymbol{X}$ are *sufficient statistics* for $\boldsymbol{B}$. Thus, by rearranging the computations, we obtain a variant of the IBSS algorithm that fits the mvSuSiE model using only sufficient statistics. We call this algorithm "IBSS-ss", and it is outlined in Algorithm 2.

We use IBSS($\boldsymbol{X}, \boldsymbol{Y}$) to denote the result of applying the IBSS algorithm (Algorithm 1) to the individual-level data, and we use IBSS-ss($\boldsymbol{X}^\intercal\boldsymbol{X}, \boldsymbol{X}^\intercal\boldsymbol{Y}$) to denote the result of applying the IBSS-ss algorithm (Algorithm 2) to the sufficient statistics. These two algorithms will give the same result; that is, IBSS($\boldsymbol{X}, \boldsymbol{Y}$) = IBSS-ss($\boldsymbol{X}^\intercal\boldsymbol{X}, \boldsymbol{X}^\intercal\boldsymbol{Y}$). However, the computational complexity of the two approaches is different. The computational complexity of one iteration of the IBSS algorithm is $O(L \times (NJR + KJR^3))$, whereas the complexity of a single iteration of the IBSS-ss algorithm is $O(L \times (J^2R + KJR^3))$. Therefore, when $N \gg J$, which is often the case in fine-mapping studies, IBSS-ss will usually be faster. We note, however, that computing the $J \times J$ matrix $\boldsymbol{X}^\intercal\boldsymbol{X}$ can be expensive, and potentially more expensive than running mvSuSiE itself. So IBSS-ss will be more computationally attractive than IBSS if $N$ is much larger than $J$ and if $\boldsymbol{X}^\intercal\boldsymbol{X}$ can be computed efficiently using a software such as PLINK [24] or LDstore [25].

*mvSuSiE with summary data: mvSuSiE-RSS.* We define mvSuSiE-RSS as the application the IBSS-ss algorithm to the sufficient statistics or approximations to these statistics (e.g., an LD estimate obtained from different genotype data than the genotype data used to obtain the other statistics). Conceptually, this approach combines the mixture prior (5) with an approximation to the likelihood (6). To formalize this, we write the likelihood as a function of the sufficient statistics,

$$\ell_{\rm ss}(\boldsymbol{B}; \boldsymbol{S}_{xx}, \boldsymbol{S}_{xy}, N) := |2\pi\boldsymbol{V}|^{-N/2} \exp\big\{ -\tfrac{N}{2}\mathrm{tr}[\boldsymbol{V}^{-1}(\boldsymbol{Y}^\intercal\boldsymbol{Y}/N - 2\boldsymbol{B}^\intercal\boldsymbol{S}_{xy} + \boldsymbol{B}^\intercal\boldsymbol{S}_{xx}\boldsymbol{B})]\big\}, \tag{7}$$

so that

$$\ell_{\rm ss}\big(\boldsymbol{B}; \tfrac{1}{N}\boldsymbol{X}^\intercal\boldsymbol{X}, \tfrac{1}{N}\boldsymbol{X}^\intercal\boldsymbol{Y}, N\big) = \ell(\boldsymbol{B}; \boldsymbol{X}, \boldsymbol{Y}). \tag{8}$$

Replacing $\boldsymbol{S}_{xx} = \frac{1}{N}\boldsymbol{X}^\intercal\boldsymbol{X}$ with an estimate $\hat{\boldsymbol{S}}_{xx} \approx \boldsymbol{S}_{xx}$ is therefore the same as replacing the sufficient-statistics likelihood (7) with

$$\ell_{\rm RSS}(\boldsymbol{B}) := \ell_{\rm ss}\big(\boldsymbol{B}; \hat{\boldsymbol{S}}_{xx}, \tfrac{1}{N}\boldsymbol{X}^\intercal\boldsymbol{Y}, N\big). \tag{9}$$

Note that when $\hat{\boldsymbol{S}}_{xx} = \boldsymbol{S}_{xx}$, the approximation is exact; that is, $\ell_{\rm RSS}(\boldsymbol{B}) = \ell(\boldsymbol{B}; \boldsymbol{X}, \boldsymbol{Y})$.

In summary, applying mvSuSiE with $\boldsymbol{S}_{xx}$ is equivalent to using the individual-data likelihood (6), and applying mvSuSiE with $\hat{\boldsymbol{S}}_{xx}$ uses the approximate likelihood (9).

*Special case when $\boldsymbol{X}, \boldsymbol{Y}$ are standardized: mvSuSiE-RSS with $z$-scores.* Now we consider the special case when $\boldsymbol{X}$ and $\boldsymbol{Y}$ are standardized, which is common in genetic association studies. By "standardized", we mean that the columns of $\boldsymbol{X}$ and $\boldsymbol{Y}$ have been scaled to have unit variance; $\sum_{i=1}^N x_{ij}^2 = N$, $j = 1, \ldots, J$, and $\sum_{i=1}^N y_{ir}^2 = N$, $r = 1, \ldots, R$. (See [1] for exact definitions of the $z$-scores and the LD matrix $\boldsymbol{R}$.) This is in addition to the assumption, mentioned earlier, that the columns of $\boldsymbol{X}$ and $\boldsymbol{Y}$ are centered to have means of zero. See [26, 27] for a discussion on the choice to standardize.

With standardized $\boldsymbol{X}$ and $\boldsymbol{Y}$, the sufficient statistics $\boldsymbol{X}^\intercal\boldsymbol{Y}$ and $\boldsymbol{X}^\intercal\boldsymbol{X}$ can be recovered from the sample size, $N$, the (in-sample) LD matrix $\boldsymbol{R}$, and the marginal association $z$-scores $\hat{z}_{jr}$, which are obtained from simple linear regressions between the traits $r$ and the SNPs $j$. (The $z$-scores should ideally be computed using the same samples for each trait so that the correlations among SNPs are same for all traits.) In particular, the sufficient statistics are recovered by the following two equations,

$$\boldsymbol{X}^\intercal\boldsymbol{X} = N \times \boldsymbol{R} \tag{10}$$

$$\boldsymbol{X}^\intercal\boldsymbol{Y} = \sqrt{N} \times \tilde{\boldsymbol{Z}} \tag{11}$$

in which $\tilde{\boldsymbol{Z}}$ denotes the $J \times R$ matrix of "adjusted $z$-scores",

$$\tilde{z}_{jr} := \frac{N}{N + \hat{z}_{jr}} \times \hat{z}_{jr}. \tag{12}$$

(Note that, when the effects are small, $\tilde{z}_{jr} \approx \hat{z}_{jr}$.)

Substituting equations (10–11) into the sufficient-statistics likelihood (7) gives

$$\ell_{\rm ss}(\boldsymbol{B}; \boldsymbol{S}_{xx}, \boldsymbol{S}_{xy}, N) = \ell_{\rm ss}(\boldsymbol{B}; \boldsymbol{R}, \tilde{\boldsymbol{Z}}/\sqrt{N}, N). \tag{13}$$

When the in-sample LD matrix $\boldsymbol{R}$ is not available, and is replaced with $\hat{\boldsymbol{R}} \approx \boldsymbol{R}$, the mvSuSiE-RSS likelihood (9) becomes

$$\ell_{\mathrm{RSS}}(\boldsymbol{B}) = \ell_{\mathrm{ss}}(\boldsymbol{B}; \hat{\boldsymbol{R}}, \tilde{\boldsymbol{Z}}/\sqrt{N}, N). \tag{14}$$

In summary, when $\boldsymbol{X}$ and $\boldsymbol{Y}$ are standardized, applying mvSuSiE with $\boldsymbol{R}$ is equivalent to using the individual-data likelihood (6), and applying mvSuSiE with $\hat{\boldsymbol{R}}$ is equivalent to using the approximate likelihood (14).

#### *mvSuSiE posterior statistics*

Here we describe the posterior statistics used in an mvSuSiE analysis.

*Basic posterior quantities.* We start with two basic posterior statistics that are used to calculate other statistics. The first posterior quantity is the posterior probability that the $l$th single effect is nonzero for SNP $j$,

$$\alpha_j^{(l)} := \Pr(\gamma_j^{(l)} = 1 \mid \boldsymbol{X}, \boldsymbol{Y}). \tag{15}$$

The second posterior quantity, denoted $\mathit{clfsr}_{jr}^{((l)}$, is the *local false sign rate* [2, 28] for SNP $j$ in trait $r$ and single effect $l$ *conditioned on* SNP $j$ having a nonzero effect in single effect $l$,

$$\mathit{clfsr}_{jr}^{(l)} := 1 - \max\{\Pr(b_{jr}^{(l)} > 0 \mid \boldsymbol{X}, \boldsymbol{Y}, \gamma_j^{(l)} = 1), \Pr(b_{jr}^{(l)} < 0 \mid \boldsymbol{X}, \boldsymbol{Y}, \gamma_j^{(l)} = 1)\}. \tag{16}$$

Intuitively, the *clfsr* ("conditional *lfsr*") measures how confident we can be in the sign of the effect of SNP $j$ in trait $r$ and single effect $l$ given that SNP $j$ has a nonzero effect in single effect $l$. A small *clfsr* indicates a small posterior probability that the sign of the estimated effect is incorrect, and thus a high confidence in the sign of an effect. The *lfsr* is similar to the commonly used local false discovery rate (*lfdr*), but more robust to modeling assumptions [28], which is helpful for reducing sensitivity to the choice of prior.

*Cross-trait PIPs.* The posterior inclusion probability (PIP) is a standard quantity reported by most fine-mapping methods, so PIPs are convenient for comparing performance of different fine-mapping methods. PIPs are also useful for visualizing the fine-mapping signal within a candidate fine-mapping region. For mvSuSiE, we define the PIP for SNP $j$ as the posterior probability that at least one of the regression coefficients for the $j$th SNP is not zero,

$$\begin{aligned} \mathrm{PIP}_j &:= \Pr(\boldsymbol{b}_j \neq \boldsymbol{0} \mid \boldsymbol{X}, \boldsymbol{Y}) \\ &= 1 - \Pr(\boldsymbol{b}_j = \boldsymbol{0} \mid \boldsymbol{X}, \boldsymbol{Y}) \\ &= 1 - \textstyle\prod_{l=1}^{L}(1 - \alpha_j^{(l)}). \end{aligned} \tag{17}$$

in which $\alpha_j^{(l)}$ is defined in (15).

*minimum lfsr.* The PIP tells us whether or not a SNP has an effect on at least one trait, but it does not tell us *which traits* are affected by the SNP. To quantify this, we calculate a *minimum lfsr* (*min-lfsr*), which we define as the smallest *lfsr* among the $L$ single effects,

$$\mathit{min\text{-}lfsr}_{jr} := \min_{l \in \{1,\ldots,L\}} \mathit{lfsr}_{jr}^{(l)}, \tag{18}$$

in which $\mathit{lfsr}_{jr}^{(l)}$ is the (unconditional) *lfsr* for SNP $j$ in outcome $r$ and single effect $l$,

$$\begin{aligned} \mathit{lfsr}_{jr}^{(l)} &:= 1 - \max\{\Pr(b_{jr}^{(l)} > 0 \mid \boldsymbol{X}, \boldsymbol{Y}), \Pr(b_{jr}^{(l)} < 0 \mid \boldsymbol{X}, \boldsymbol{Y})\} && (19) \\ &= 1 - \alpha_j^{(l)}(1 - \mathit{clfsr}_{jr}^{(l)}), && (20) \end{aligned}$$

and we use the definition of $\mathit{clfsr}_{jr}^{(l)}$ from (16). Intuitively, SNP $j$ is considered "significant" in trait $r$ if and only if it is significant in at least one of the $L$ effects.

*Credible sets.* A cross-trait credible set $\mathrm{CS}(\boldsymbol{\alpha}^{(l)}; \rho)$ is defined as a set of SNPs that has probability at least $\rho$ of containing an effect SNP [29]. The calculation of cross-trait CSs is described in [13].

A CS does not indicate *which traits* are affected by the SNPs. To assess significance of a CS for a specific trait $r$, we compute the *average lfsr*, defined as an average of the conditional *lfsr*s for all SNPs $j$ weighted by their posterior probabilities $\alpha_j^{(l)}$:

$$\mathit{lfsr}_r^{(l)} := \sum_{j=1}^{J} \alpha_j^{(l)}\, \mathit{clfsr}_{jr}^{(l)}. \tag{21}$$

If the *average lfsr* for trait $r$ is small, this indicates high confidence in the sign of the effect—small posterior probability that the sign is incorrect—and so we say the effects of the SNPs in the CS are *significant for trait* $r$ or we say the CS is a *trait-wise significant CS for trait* $r$.

#### *Specifying* $\boldsymbol{V}$ *and* $g$

In order to run mvSuSiE or mvSuSiE-RSS, we must first specify the $R \times R$ residual variance-covariance matrix, $\boldsymbol{V}$, and the prior on the regression coefficients, $g$. In the next two sections, we describe the steps that were taken to specify these model parameters in the simulations and the UK Biobank blood traits case study. Since we always applied mvSuSiE to summary data ("mvSuSiE-RSS"), and specifically $z$-scores, we describe estimation of $\boldsymbol{V}$ and $g$ using the $z$-scores.

*Estimating the residual variance matrix.* When the traits are measured in the same samples, it is important to account for possible correlations among the measurements of the different traits; failure to do so can result in miscalibrated fine-mapping statistics (Supplementary Fig. 1). The residual covariance matrix $\boldsymbol{V}$ is used to account for correlations among the measurements. The special case of independent measurements can be modeled by setting $\boldsymbol{V}$ to a diagonal matrix.

To estimate $\boldsymbol{V}$, we adapted the approach from [2], in which $\boldsymbol{V}$ was estimated from $z$-scores (e.g., $z$-scores obtained from marginal association tests). Specifically, we took the following steps. First, we pooled the $z$-scores from all the fine-mapping regions considered. Then we filtered out $z$-scores that are large in magnitude; specifically, we only considered SNPs in which the largest $z$-score magnitude for any trait was less than 2. This improved the estimate of $\boldsymbol{V}$ by removing SNPs that might affect one or more of the traits. Denoting the number of SNPs used in this calculation by $J'$, and letting $\hat{\boldsymbol{z}}_j$ denote the vector of $z$-scores obtained from the $R$ association tests for SNP $j$, we estimated $\boldsymbol{V}$ as

$$\hat{\boldsymbol{V}} = \frac{1}{J'} \sum_{j=1}^{J'} \hat{\boldsymbol{z}}_j \hat{\boldsymbol{z}}_j^{\intercal}. \tag{22}$$

We estimated $\boldsymbol{V}$ for the UK Biobank blood cell trait data using $J' = 1{,}950$ SNPs. (Two SNPs with small $z$-scores were selected from each of the 975 fine-mapping regions.) This estimate is given in Supplementary Table 8 ($n = 1{,}950$). Since the blood cell traits were standardized, we rescaled the estimate so that the final $\boldsymbol{V}$ used in the fine-mapping analyses was a correlation matrix.

*Specifying the prior.* The prior (5) can accommodate many different effect sharing patterns. However, to maximize the benefit of using this prior, it should capture the effect sharing patterns that are actually present in the data. Following [2], we considered three approaches to obtaining $g$:

1. A "random effects prior" that assumes the effects are independent across traits. This is a special case of (5) in which the mixture consists of a single mixture component ($K = 1, \omega_1 = 1$) with covariance matrix $\boldsymbol{U}_1 = \boldsymbol{I}_R$. Although simple, this prior is actually used (implicitly or explicitly) by methods that assume the effects are independent across traits.

2. A prior with a mixture of "canonical" sharing patterns. (See below for details.) This prior is not as flexible as the "data-driven" prior described next, but has the advantage of being easy to implement because it does not involve any separate estimation steps.

3. A "data-driven" prior in which the covariances and weights are estimated from the data. The basic idea behind this prior is to adapt the sharing patterns $\boldsymbol{U}_k$ and corresponding mixture weights $\omega_k$ to the data. (See below for details.) This requires more work to design but has a potentially greater payoff.

In simulations, we assessed the advantages of each of these approaches (Supplementary Fig. 2).

*Canonical prior.* The generation of the canonical covariance matrices for the prior $g$ is implemented by the `create_cov_canonical` function in the `mvsusieR` package. Following [2], this function generates the following covariance matrices: the $R \times R$ identity matrix, $\boldsymbol{I}_R$, modeling the case when all effects are independent (the same as the "random effects" prior); the "equal effects" matrix, an $R \times R$ matrix of all ones, which models the case in which all effects are the same; rank-1 matrices modeling trait-specific effects of the form $\boldsymbol{e}_r\boldsymbol{e}_r^\intercal$, in which $\boldsymbol{e}_r$ is a vector of length $R$ containing all zeros except for a 1 in the $r$th position; and matrices modeling uniformly heterogeneous effects, with ones on the diagonal and $\sigma$ on the off-diagonal, where $\sigma$ is 0.25, 0.5 or 0.75. In total, this results in $R + 5$ covariance matrices, where $R$ is the number of traits. Note that the canonical covariance matrices are all at the same scale (each matrix has entries spanning the range 0 to 1), and none of the matrices allow for negatively correlated effects (all of the entries in these matrices are non-negative).

To complete the canonical prior, we assigned uniform weights $\omega_k = 1/K$ to the $K = R + 5$ mixture components.

*Data-driven prior.* We also took an approach similar to [2] to generate the covariance matrices $\boldsymbol{U}_k$ and mixture weights $\omega_k$ in the "data-driven" prior.

First, we prepared a data set to learn the prior. For each candidate fine-mapping region, we identified the top $z$-score which was defined as the vector of association $z$-scores for the $R$ traits containing the largest (in magnitude) $z$-score among all SNPs in the given fine-mapping region. Letting $M$ denote the number of fine-mapping regions, we formed an $M \times R$ matrix containing the top $z$-scores. Here we denote this matrix by $\boldsymbol{Z}$.

Next, we generated *initial estimates* of covariance matrices using a variety of approaches:

1. $R + 5$ canonical covariance matrices (see above).

2. The empirical covariance matrix $\boldsymbol{Z}^\intercal\boldsymbol{Z}/M$.

3. Three rank-1 matrices of the form $\boldsymbol{v}_r\boldsymbol{v}_r^\intercal$, $r = 1, 2, 3$, in which $\boldsymbol{v}_r$ is the $r$th right singular vector of the reduced singular value decomposition (SVD) of $\boldsymbol{Z}$, $\boldsymbol{Z} = \sum_{r=1}^{R} \sigma_r \boldsymbol{w}_r \boldsymbol{v}_r{}^\intercal$, in which $\sigma_r$ is the $r$th singular value and $\boldsymbol{w}_r$ is the $r$th left singular vector.

4. A rank-3 approximation of $\boldsymbol{Z}$ based its SVD, $\boldsymbol{U} \approx \sum_{r=1}^{3} \sigma_r^2 \boldsymbol{v}_r\boldsymbol{v}_r^\intercal/M$.

5. A sparse, low-rank approximation of $\boldsymbol{Z}$ obtained using the R package `flashr` [30] (version 0.6-8), $\boldsymbol{U} \approx \boldsymbol{F}\boldsymbol{L}^\intercal\boldsymbol{L}\boldsymbol{F}^\intercal/M$, where $\boldsymbol{L}$ is the $M \times R'$ loadings matrix and $\boldsymbol{F}$ is the $R \times R'$ matrix of estimated factors, and $R' \leqslant R$ is the rank of the approximation. The rank, $R'$, was automatically determined by `flashr`.

6. $R'$ rank-1 matrices of the form $\boldsymbol{f}_r\boldsymbol{f}_r^\intercal$, $r = 1, \ldots, R'$, in which $\boldsymbol{f}_r$ denotes the $r$th column of $\boldsymbol{F}$.

After completing these steps, we had initial estimates for $K = R + R' + 11$ covariance matrices.

Next, we ran Extreme Deconvolution (ED) [31] to refine these initial estimates and simultaneously estimate the mixture weights $\omega_k$. (We used the ED algorithm implemented in the `cov_ed` function from the `mashr` R package, version 0.2.59, which was adapted from [31].)

Finally, to avoid poor estimation of the *lfsr* that can happen when the prior covariances are singular (*i.e.*, not invertible), we added a small positive constant to the diagonals of all the covariance matrices $\boldsymbol{U}_k$ in the data-driven prior. This step ensured that these matrices were all invertible.

The data-driven prior obtained by running the above procedure on the UK Biobank data is shown in Supplementary Fig. 5. The data-driven priors obtained by running the above procedure separately in Scenarios A and B of the simulations are shown in Supplementary Figures 6 and 7.

#### *The multivariate simple regression model*

The mvSuSiE model is based on a simple multivariate regression model with one variable which we refer to as the "multivariate simple regression model." This model is

$$\boldsymbol{Y} \sim MN_{N \times R}(\boldsymbol{x}\boldsymbol{b}^{\intercal}, \boldsymbol{I}_N, \boldsymbol{V}). \tag{23}$$

where $\boldsymbol{Y} \in \mathbb{R}^{N \times R}$ is a matrix of $R$ observed responses in $N$ samples, $\boldsymbol{x}$ is a vector of $N$ observations for a single explanatory variable, $\boldsymbol{b} \in \mathbb{R}^R$ is the (unknown) vector of regression coefficients for the $R$ responses, $\boldsymbol{V} \in \mathbb{S}_{++}^R$ is an invertible covariance matrix, and $MN_{n \times m}(\boldsymbol{M}, \boldsymbol{U}, \boldsymbol{V})$ is the matrix normal distribution [32, 33] with mean $\boldsymbol{M} \in \mathbb{R}^{n \times m}$ and covariance matrices $\boldsymbol{U}, \boldsymbol{V} \in \mathbb{S}_+^{m \times m}$. For now, this model does not include an intercept.

Next, we give the likelihood for (23) and relate it to the more familiar multivariate normal distribution. The likelihood for the multivariate regression model (23) is

$$\begin{aligned} \ell(\boldsymbol{b}; \boldsymbol{x}, \boldsymbol{Y}, \boldsymbol{V}) &:= p(\boldsymbol{Y} \mid \boldsymbol{x}, \boldsymbol{b}, \boldsymbol{V}) \\ &= |2\pi \boldsymbol{V}|^{-N/2} \exp\big\{ -\tfrac{1}{2}\mathrm{tr}[\boldsymbol{V}^{-1}(\boldsymbol{Y} - \boldsymbol{x}\boldsymbol{b}^{\intercal})^{\intercal}(\boldsymbol{Y} - \boldsymbol{x}\boldsymbol{b}^{\intercal})]\big\}. \end{aligned} \tag{24}$$

Given $\boldsymbol{V}$, the least-squares estimate of $\boldsymbol{b}$, denoted by $\hat{\boldsymbol{b}}$, and its variance-covariance matrix, $\hat{\boldsymbol{S}}$, are

$$\hat{\boldsymbol{b}} = \tfrac{\boldsymbol{Y}^{\intercal}\boldsymbol{x}}{\boldsymbol{x}^{\intercal}\boldsymbol{x}} \tag{25}$$

$$\hat{\boldsymbol{S}} = \tfrac{\boldsymbol{V}}{\boldsymbol{x}^{\intercal}\boldsymbol{x}}. \tag{26}$$

Note that $\hat{\boldsymbol{b}}$ is also the value of $\boldsymbol{b}$ maximizing the likelihood (24). Using these quantities, the likelihood (24) can be rewritten as

$$\ell(\boldsymbol{b}; \boldsymbol{x}, \boldsymbol{Y}, \boldsymbol{V}) = |2\pi \boldsymbol{V}|^{-N/2} \exp\big\{ -\tfrac{1}{2}[\mathrm{tr}(\boldsymbol{V}^{-1}\boldsymbol{Y}^{\intercal}\boldsymbol{Y}) + (\boldsymbol{b} - \hat{\boldsymbol{b}})^{\intercal}\hat{\boldsymbol{S}}^{-1}(\boldsymbol{b} - \hat{\boldsymbol{b}}) - \hat{\boldsymbol{b}}^{\intercal}\hat{\boldsymbol{S}}^{-1}\hat{\boldsymbol{b}}]\big\}. \tag{27}$$

This expression is convenient because the terms involving $\boldsymbol{b}$ are in the form of a multivariate normal density up to a constant of proportionality, and in particular we have that

$$\ell(\boldsymbol{b}; \boldsymbol{x}, \boldsymbol{Y}, \boldsymbol{V}) \propto N_R(\boldsymbol{b}; \hat{\boldsymbol{b}}, \hat{\boldsymbol{S}}), \tag{28}$$

where $N_d(\boldsymbol{\theta}; \boldsymbol{\mu}, \boldsymbol{\Sigma})$ denotes the multivariate normal density at $\boldsymbol{\theta} \in \mathbb{R}^d$ with mean $\boldsymbol{\mu} \in \mathbb{R}^d$ and covariance $\boldsymbol{\Sigma} \in \mathbb{S}_+^d$.

Remark 1. Calculation of $\hat{\boldsymbol{b}}$ and $\hat{\boldsymbol{S}}$ only requires the summary statistics $\boldsymbol{x}^{\intercal}\boldsymbol{x}$, $\boldsymbol{Y}^{\intercal}\boldsymbol{x}$. Also, since the likelihood (28) only involves $\hat{\boldsymbol{b}}$ and $\hat{\boldsymbol{S}}$, the likelihood up to a constant of proportionality can also be computed with only these summary statistics.

#### *The multivariate simple regression model with a normal prior*

In the following proposition, we apply the above results for the multivariate simple regression model to a Bayesian multivariate simple regression model with a normal prior.

Proposition 1 (Bayesian multivariate simple regression with a normal prior). Consider the multivariate simple regression model (23) with a multivariate normal prior on the regression coefficients,

$$\boldsymbol{b} \mid \boldsymbol{S}_0 \sim N_R(\boldsymbol{0}, \boldsymbol{S}_0), \tag{29}$$

where $\boldsymbol{S}_0 \in \mathbb{S}_+^R$ is a (possibly singular) covariance matrix. The posterior of $\boldsymbol{b}$ is

$$\boldsymbol{b} \mid \boldsymbol{x}, \boldsymbol{Y}, \boldsymbol{V}, \boldsymbol{S}_0 \sim N_R(\boldsymbol{b}_1, \boldsymbol{S}_1), \tag{30}$$

where

$$\boldsymbol{b}_1 = \boldsymbol{S}_1\hat{\boldsymbol{S}}^{-1}\hat{\boldsymbol{b}} \tag{31}$$

$$\boldsymbol{S}_1 = (\boldsymbol{S}_0^{-1} + \hat{\boldsymbol{S}}^{-1})^{-1}. \tag{32}$$

The Bayes Factor (BF) comparing this model against the null model ($\boldsymbol{b} = \boldsymbol{0}$) is

$$\begin{aligned}\mathrm{BF}(\boldsymbol{x}, \boldsymbol{Y}, \boldsymbol{V}, \boldsymbol{S}_0) &= \frac{p(\boldsymbol{Y} \mid \boldsymbol{x}, \boldsymbol{V}, \boldsymbol{S}_0)}{p(\boldsymbol{Y} \mid \boldsymbol{x}, \boldsymbol{V}, \boldsymbol{b} = \boldsymbol{0})} \\ &= \frac{\int \ell(\boldsymbol{b}; \boldsymbol{x}, \boldsymbol{Y}, \boldsymbol{V})\, p(\boldsymbol{b} \mid \boldsymbol{S}_0)\, d\boldsymbol{b}}{\ell(\boldsymbol{b} = \boldsymbol{0}; \boldsymbol{x}, \boldsymbol{Y}, \boldsymbol{V})}, \end{aligned} \tag{33}$$

which simplifies to

$$\begin{aligned}\mathrm{BF}(\boldsymbol{x}, \boldsymbol{Y}, \boldsymbol{V}, \boldsymbol{S}_0) &= \frac{|\hat{\boldsymbol{S}}|^{1/2}}{|\boldsymbol{S}_0 + \hat{\boldsymbol{S}}|^{1/2}} \exp(\tfrac{1}{2}\hat{\boldsymbol{b}}^\intercal \hat{\boldsymbol{S}}^{-1} \boldsymbol{S}_1 \hat{\boldsymbol{S}}^{-1} \hat{\boldsymbol{b}}) & (34) \\ &= \frac{|\hat{\boldsymbol{S}}|^{1/2}}{|\boldsymbol{S}_0 + \hat{\boldsymbol{S}}|^{1/2}} \exp(\tfrac{1}{2}\boldsymbol{b}_1^\intercal \boldsymbol{S}_1^{-1} \boldsymbol{b}_1). & (35)\end{aligned}$$

The same BF can also be equivalently expressed as a ratio of two multivariate normal densities,

$$\mathrm{BF}(\boldsymbol{x}, \boldsymbol{Y}, \boldsymbol{V}, \boldsymbol{S}_0) = \frac{N_R(\hat{\boldsymbol{b}}; \boldsymbol{0}, \boldsymbol{S}_0 + \hat{\boldsymbol{S}})}{N_R(\hat{\boldsymbol{b}}; \boldsymbol{0}, \hat{\boldsymbol{S}})}. \tag{36}$$

Remark 2. Since the data $\boldsymbol{x}, \boldsymbol{Y}$ only enter the expressions for the posterior mean $\boldsymbol{b}_1$ and posterior covariance $\boldsymbol{S}_1$ through $\hat{\boldsymbol{b}}$ and $\hat{\boldsymbol{S}}$, it follows that calculation of the posterior mean and covariance only requires summary statistics $\boldsymbol{x}^\intercal\boldsymbol{x}, \boldsymbol{Y}^\intercal\boldsymbol{x}$. Similarly, the BF (36) can be computed with only $\boldsymbol{x}^\intercal\boldsymbol{x}, \boldsymbol{Y}^\intercal\boldsymbol{x}$.

*Updating a scaling factor in the prior.* Suppose the prior covariance is parameterized as $\boldsymbol{S}_0 = \sigma_0^2 \boldsymbol{U}$, in which $\boldsymbol{U} \in \mathbb{S}_+^R$, $\sigma_0 \geqslant 0$. Here we assume $\boldsymbol{U}$ is a fixed parameter and we would like to estimate the scaling factor $\sigma_0$ by maximizing the likelihood,

$$\hat{\sigma}_0^2 := \mathrm{argmax}_{\sigma_0^2}\, p(\boldsymbol{Y} \mid \boldsymbol{x}, \boldsymbol{V}, \sigma_0^2\boldsymbol{U}), \tag{37}$$

or, equivalently, by maximizing the Bayes Factor, which may be more convenient to compute,

$$\hat{\sigma}_0^2 = \mathrm{argmax}_{\sigma_0^2}\, \mathrm{BF}(\boldsymbol{x}, \boldsymbol{Y}, \boldsymbol{V}, \sigma_0^2\boldsymbol{U}). \tag{38}$$

When $\boldsymbol{U}$ is invertible, the maximum-likelihood estimate $\hat{\sigma}_0^2$ can be computed using a simple EM algorithm [34], in which the M-step update is

$$\sigma_0^2 = \mathrm{tr}(\boldsymbol{U}^{-1}\mathbb{E}[\boldsymbol{b}\boldsymbol{b}^\intercal])/R. \tag{39}$$

The E-step then consists of computing the posterior second moment,

$$\mathbb{E}[\boldsymbol{b}\boldsymbol{b}^\intercal] = \boldsymbol{b}_1\boldsymbol{b}_1^\intercal + \boldsymbol{S}_1. \tag{40}$$

The maximum-likelihood estimate $\hat{\sigma}_0^2$ is then recovered by iterating the E-step (40) and M-step (39) until convergence.

The update (39) requires that $\boldsymbol{U}$ be invertible. To allow for singular matrices, a more general M-step update is

$$\sigma_0^2 = \mathrm{tr}(\boldsymbol{U}^\dagger\mathbb{E}[\boldsymbol{b}\boldsymbol{b}^\intercal])/R'. \tag{41}$$

in which $R' \leqslant R$ is the rank of $\boldsymbol{U}$. Note (39) and (41) are equivalent when $\boldsymbol{U}$ is invertible; that is, when $R' = R$.

#### *The multivariate simple regression model with an intercept*

Now we extend the multivariate simple regression model (23) to include an intercept. We show that including an intercept is equivalent to "centering" $\boldsymbol{x}$ and the columns of $\boldsymbol{Y}$ so that they all have means of zero. More precisely, centering is equivalent to integrating out the intercept with respect to an (improper) uniform prior on the intercept. This is the multivariate generalization of the result for univariate regression given in [35]. This result is summarized in Proposition 2.

The multivariate simple regression model with an intercept is

$$\boldsymbol{Y} \sim MN_{N\times R}(\mathbf{1}_N\boldsymbol{\mu}^\intercal + \boldsymbol{x}\boldsymbol{b}^\intercal, \boldsymbol{I}_N, \boldsymbol{V}), \tag{42}$$

in which $\boldsymbol{\mu} \in \mathbb{R}^R$ is the (unknown) intercept. The likelihood of $\boldsymbol{\mu}, \boldsymbol{b}$ under this model is

$$\begin{aligned} \ell(\boldsymbol{\mu}, \boldsymbol{b}; \boldsymbol{x}, \boldsymbol{Y}, \boldsymbol{V}) &:= p(\boldsymbol{Y} \mid \boldsymbol{x}, \boldsymbol{\mu}, \boldsymbol{b}, \boldsymbol{V}) \\ &= |2\pi\boldsymbol{V}|^{-N/2} \exp\{-\tfrac{1}{2}\mathrm{tr}[\boldsymbol{V}^{-1}(\boldsymbol{Y} - \mathbf{1}\boldsymbol{\mu}^\intercal - \boldsymbol{x}\boldsymbol{b}^\intercal)^\intercal(\boldsymbol{Y} - \mathbf{1}\boldsymbol{\mu}^\intercal - \boldsymbol{x}\boldsymbol{b}^\intercal)]\}. \end{aligned} \tag{43}$$

Proposition 2 (Multivariate simple regression with an intercept). Consider the multivariate simple regression model with an intercept (42). The least-squares estimate of $\boldsymbol{\mu}$—which is also the value of $\boldsymbol{\mu}$ maximizing the likelihood (43)—and its covariance matrix are

$$\hat{\boldsymbol{\mu}} = \bar{\boldsymbol{y}} - \bar{x}\boldsymbol{b} \tag{44}$$

$$\hat{\boldsymbol{S}}_\mu = \tfrac{1}{N}\boldsymbol{V}, \tag{45}$$

in which $\bar{x} := \frac{1}{N}\boldsymbol{x}^\intercal\mathbf{1}_N = \frac{1}{N}\sum_{i=1}^N x_i$ is the sample mean of $\boldsymbol{x}$, and $\bar{\boldsymbol{y}} := \frac{1}{N}\boldsymbol{Y}^\intercal\mathbf{1}_N$ is the vector containing the column means of $\boldsymbol{Y}$.

The profile likelihood for $\boldsymbol{b}$ is

$$\begin{aligned} \ell^*(\boldsymbol{b}; \boldsymbol{x}, \boldsymbol{Y}, \boldsymbol{V}) &:= \max_{\boldsymbol{\mu}} \ell(\boldsymbol{\mu}, \boldsymbol{b}; \boldsymbol{x}, \boldsymbol{Y}, \boldsymbol{V}) \\ &= \ell(\boldsymbol{b}; \tilde{\boldsymbol{x}}, \tilde{\boldsymbol{Y}}, \boldsymbol{V}), \end{aligned} \tag{46}$$

in which $\tilde{\boldsymbol{x}} := \boldsymbol{x} - \bar{x}\mathbf{1}_N$ and $\tilde{\boldsymbol{Y}} := \boldsymbol{Y} - \mathbf{1}_N\bar{\boldsymbol{y}}^\intercal$ are the "centered" versions of $\boldsymbol{x}$ and $\boldsymbol{Y}$. In other words, the profile likelihood for the multivariate simple regression with an intercept is the same as the likelihood for the multivariate simple regression *without* an intercept if we first center $\boldsymbol{x}$ and $\boldsymbol{Y}$. Centering $\boldsymbol{x}$ and $\boldsymbol{Y}$ is therefore equivalent to including an intercept in the multivariate regression and estimating the intercept by maximum-likelihood.

Next, consider Bayesian calculations for $\boldsymbol{\mu}$ with a multivariate normal prior, $\boldsymbol{\mu} \mid \boldsymbol{S}_{0\mu} \sim N_R(0, \boldsymbol{S}_{0\mu})$, in which $\boldsymbol{S}_{0\mu} \in \mathbb{S}_+^R$ is a (possibly singular) covariance matrix. The posterior for $\boldsymbol{\mu}$ conditioned on $\boldsymbol{b}$ is

$$\boldsymbol{\mu} \mid \boldsymbol{x}, \boldsymbol{Y}, \boldsymbol{V}, \boldsymbol{S}_{0\mu}, \boldsymbol{b} \sim N_R(\boldsymbol{\mu}_1, \boldsymbol{S}_{1\mu}), \tag{47}$$

where

$$\boldsymbol{\mu}_1 = \boldsymbol{S}_{1\mu}\hat{\boldsymbol{S}}_\mu^{-1}\hat{\boldsymbol{\mu}} \tag{48}$$

$$\boldsymbol{S}_{1\mu} = (\boldsymbol{S}_{0\mu}^{-1} + \hat{\boldsymbol{S}}_\mu^{-1})^{-1}. \tag{49}$$

The marginal likelihood obtained by averaging over the intercept is

$$\begin{aligned} \ell^\star(\boldsymbol{b}; \boldsymbol{x}, \boldsymbol{Y}, \boldsymbol{V}, \boldsymbol{S}_{0\mu}) &:= \textstyle\int \ell(\boldsymbol{\mu}, \boldsymbol{b}; \boldsymbol{x}, \boldsymbol{Y}, \boldsymbol{V})\, p(\boldsymbol{\mu} \mid \boldsymbol{S}_{0\mu})\, d\boldsymbol{\mu} \\ &= |2\pi\boldsymbol{V}|^{-N/2}|\boldsymbol{S}_{0\mu}^{-1}\boldsymbol{S}_{1\mu}|^{1/2} \exp\{\tfrac{1}{2}\boldsymbol{\mu}_1^\intercal\boldsymbol{S}_{1\mu}^{-1}\boldsymbol{\mu}_1 - \tfrac{1}{2}\mathrm{tr}[\boldsymbol{V}^{-1}(\boldsymbol{Y} - \boldsymbol{x}\boldsymbol{b}^\intercal)^\intercal(\boldsymbol{Y} - \boldsymbol{x}\boldsymbol{b}^\intercal)]\}. \end{aligned} \tag{50}$$

In the special case of an (improper) uniform prior on $\boldsymbol{\mu}$, defined as $\boldsymbol{\mu} \sim N_R(0, \boldsymbol{S}_{0\mu})$ with $\boldsymbol{S}_{0\mu}^{-1} \to 0$, the posterior mean reduces to the least-squares estimate $\boldsymbol{\mu}_1 = \hat{\boldsymbol{\mu}}$, with covariance matrix $\boldsymbol{S}_{1\mu} = \hat{\boldsymbol{S}}_\mu$, and the marginal likelihood (50) simplifies to

$$\begin{aligned} \ell^\star(\boldsymbol{b}; \boldsymbol{x}, \boldsymbol{Y}, \boldsymbol{V}, \boldsymbol{S}_{0\mu}) &= |2\pi\boldsymbol{V}|^{-N/2}|\boldsymbol{S}_{0\mu}^{-1}\hat{\boldsymbol{S}}_\mu|^{1/2} \exp\left\{\tfrac{1}{2}\hat{\boldsymbol{\mu}}^\intercal\hat{\boldsymbol{S}}_\mu^{-1}\hat{\boldsymbol{\mu}} - \tfrac{1}{2}\mathrm{tr}\big[\boldsymbol{V}^{-1}(\boldsymbol{Y} - \boldsymbol{x}\boldsymbol{b}^\intercal)^\intercal(\boldsymbol{Y} - \boldsymbol{x}\boldsymbol{b}^\intercal)\big]\right\} \\ &= |\boldsymbol{S}_{0\mu}^{-1}\hat{\boldsymbol{S}}_\mu|^{1/2} \times \ell(\boldsymbol{b}; \tilde{\boldsymbol{x}}, \tilde{\boldsymbol{Y}}, \boldsymbol{V}). \end{aligned} \tag{51}$$

In other words, the marginal likelihood for multivariate simple regression with an intercept (42), when we use an improper uniform prior for the intercept, is the same (up to a constant of proportionality) as the likelihood for multivariate simple regression *without* an intercept (23) after first centering $\boldsymbol{x}$ and $\boldsymbol{Y}$.

Remark 3. To account for an intercept when computing posterior quantities and BFs for the multivariate simple regression model (23), $\boldsymbol{x}$ and $\boldsymbol{Y}$ should be centered before computing the summary statistics; that is, the summary statistics should be $\tilde{\boldsymbol{x}}^\intercal\tilde{\boldsymbol{x}}$ and $\tilde{\boldsymbol{Y}}^\intercal\tilde{\boldsymbol{x}}$. See [1] for how to center summary statistics if they are not already centered.

*The Bayesian multivariate simple regression model with a mixture prior*

Here we extend the Bayesian multivariate simple regression model with a normal prior to a model with a mixture-of-normals prior,

$$\boldsymbol{b} \mid \mathcal{S}_0, \boldsymbol{\omega} \sim \sum_{k=1}^{K} \omega_k N_R(\mathbf{0}, \boldsymbol{S}_{0k}), \tag{52}$$

in which $\mathcal{S}_0 := \{\boldsymbol{S}_{01}, \ldots, \boldsymbol{S}_{0K}\}$, each $\boldsymbol{S}_{0k} \in \mathbb{S}_+^R$ is a (possibly singular) covariance matrix, and $\boldsymbol{\omega} := (\omega_1, \ldots, \omega_K) \in \Delta^K$ are the mixture weights. Note that the normal prior (29) is as a special case of (52) when $K = 1$.

To facilitate derivation of the posterior computations, we introduce the following data augmentation that recovers (52) after integrating over a latent random variable $\xi \in \{1, \ldots, K\}$,

$$\begin{aligned} p(\xi = k \mid \boldsymbol{\omega}) &= \omega_k \\ \boldsymbol{b} \mid \mathcal{S}_0, \xi = k &\sim N_R(\mathbf{0}, \boldsymbol{S}_{0k}). \end{aligned} \tag{53}$$

This augmented model allows us to reuse the posterior computations from the simpler models; in particular, posterior computations conditioned on $\xi$ reduce to computations for the Bayesian multivariate regression model with a normal prior, which we state more formally in the following proposition.

PROPOSITION 3. Given $\mathcal{S}_0$ and $\boldsymbol{\omega}$, the Bayes factor comparing this model against the null model ($\boldsymbol{b} = \mathbf{0}$) is

$$\begin{aligned} \mathrm{BF}^{\mathsf{mix}}(\boldsymbol{x}, \boldsymbol{Y}, \boldsymbol{V}, \mathcal{S}_0, \boldsymbol{\omega}) &= \frac{p(\boldsymbol{Y} \mid \boldsymbol{x}, \boldsymbol{V}, \mathcal{S}_0, \boldsymbol{\omega})}{p(\boldsymbol{Y} \mid \boldsymbol{x}, \boldsymbol{V}, \boldsymbol{b} = \mathbf{0})} \\ &= \sum_{k=1}^{K} \omega_k \mathrm{BF}(\boldsymbol{x}, \boldsymbol{Y}, \boldsymbol{V}, \boldsymbol{S}_{0k}), \end{aligned} \tag{54}$$

where the expressions for the individual BFs in the sum are given in Proposition 1. The posterior distribution of $\boldsymbol{b}$ is a mixture of normals,

$$\boldsymbol{b} \mid \boldsymbol{x}, \boldsymbol{Y}, \boldsymbol{V}, \mathcal{S}_0, \boldsymbol{\omega} \sim \sum_{k=1}^{K} \omega_{1k} N_R(\boldsymbol{b}_{1k}, \boldsymbol{S}_{1k}), \tag{55}$$

in which $\boldsymbol{b}_{1k}$ and $\boldsymbol{S}_{1k}$ are the posterior mean and covariance of $\boldsymbol{b}$ conditioned on $\xi = k$, given by (31) and (32), respectively, after substituting $\boldsymbol{S}_0$ with $\boldsymbol{S}_{0k}$,

$$\boldsymbol{b}_{1k} := \boldsymbol{b}_{1k}(\boldsymbol{x}, \boldsymbol{Y}, \boldsymbol{V}, \mathcal{S}_0, \boldsymbol{\omega}) = \boldsymbol{S}_{1k} \hat{\boldsymbol{S}}^{-1} \hat{\boldsymbol{b}} \tag{56}$$

$$\boldsymbol{S}_{1k} := \boldsymbol{S}_{1k}(\boldsymbol{x}, \boldsymbol{Y}, \boldsymbol{V}, \mathcal{S}_0, \boldsymbol{\omega}) = (\boldsymbol{S}_{0k}^{-1} + \hat{\boldsymbol{S}}^{-1})^{-1}, \tag{57}$$

and the posterior mixture assignment probabilities are

$$\begin{aligned} \omega_{1k} &:= \omega_{1k}(\boldsymbol{x}, \boldsymbol{Y}, \boldsymbol{V}, \mathcal{S}_0, \boldsymbol{\omega}) \\ &= p(\xi = k \mid \boldsymbol{x}, \boldsymbol{Y}, \boldsymbol{V}, \mathcal{S}_0, \boldsymbol{\omega}) \\ &= \frac{\omega_k \mathrm{BF}(\boldsymbol{x}, \boldsymbol{Y}, \boldsymbol{V}, \boldsymbol{S}_{0k})}{\sum_{k'=1}^{K} \omega_{k'} \mathrm{BF}(\boldsymbol{x}, \boldsymbol{Y}, \boldsymbol{V}, \boldsymbol{S}_{0k'})}. \end{aligned} \tag{58}$$

The posterior mean and covariance of $\boldsymbol{b}$ are

$$\boldsymbol{b}_1^{\mathsf{mix}} := \boldsymbol{b}_1^{\mathsf{mix}}(\boldsymbol{x}, \boldsymbol{Y}, \boldsymbol{V}, \mathcal{S}_0, \boldsymbol{\omega}) = \textstyle\sum_{k=1}^{K} \omega_{1k} \boldsymbol{b}_{1k} \tag{59}$$

$$\boldsymbol{S}_1^{\mathsf{mix}} := \boldsymbol{S}_1^{\mathsf{mix}}(\boldsymbol{x}, \boldsymbol{Y}, \boldsymbol{V}, \mathcal{S}_0, \boldsymbol{\omega}) = \textstyle\sum_{k=1}^{K} \omega_{1k} (\boldsymbol{b}_{1k} \boldsymbol{b}_{1k}^{\mathsf{T}} + \boldsymbol{S}_{1k}) - \boldsymbol{b}_1^{\mathsf{mix}} (\boldsymbol{b}_1^{\mathsf{mix}})^{\mathsf{T}}. \tag{60}$$

From the above remarks, the posterior quantities and Bayes factors for this model can be computed using the summary statistics $\boldsymbol{x}^{\mathsf{T}}\boldsymbol{x}$, $\boldsymbol{Y}^{\mathsf{T}}\boldsymbol{x}$ instead of using the full data $\boldsymbol{x}, \boldsymbol{Y}$. To formalize these computations with summary statistics, we introduce notation for Bayes factors and posteriors in terms of summary statistics:

$$\boldsymbol{b}_{1k}^{\mathsf{ss}}(\boldsymbol{x}^\intercal\boldsymbol{x}, \boldsymbol{Y}^\intercal\boldsymbol{x}, \boldsymbol{V}, \mathcal{S}_0, \boldsymbol{\omega}) := \boldsymbol{b}_{1k}(\boldsymbol{x}, \boldsymbol{Y}, \boldsymbol{V}, \mathcal{S}_0, \boldsymbol{\omega}) \tag{61}$$

$$\boldsymbol{S}_{1k}^{\mathsf{ss}}(\boldsymbol{x}^\intercal\boldsymbol{x}, \boldsymbol{Y}^\intercal\boldsymbol{x}, \boldsymbol{V}, \mathcal{S}_0, \boldsymbol{\omega}) := \boldsymbol{S}_{1k}(\boldsymbol{x}, \boldsymbol{Y}, \boldsymbol{V}, \mathcal{S}_0, \boldsymbol{\omega}) \tag{62}$$

$$\omega_{1k}^{\mathsf{ss}}(\boldsymbol{x}^\intercal\boldsymbol{x}, \boldsymbol{Y}^\intercal\boldsymbol{x}, \boldsymbol{V}, \mathcal{S}_0, \boldsymbol{\omega}) := \omega_{1k}(\boldsymbol{x}, \boldsymbol{Y}, \boldsymbol{V}, \mathcal{S}_0, \boldsymbol{\omega}) \tag{63}$$

$$\mathrm{BF}^{\mathsf{mix\text{-}ss}}(\boldsymbol{x}^\intercal\boldsymbol{x}, \boldsymbol{Y}^\intercal\boldsymbol{x}, \boldsymbol{V}, \mathcal{S}_0, \boldsymbol{\omega}) := \mathrm{BF}^{\mathsf{mix}}(\boldsymbol{x}, \boldsymbol{Y}, \boldsymbol{V}, \mathcal{S}_0, \boldsymbol{\omega}). \tag{64}$$

*Updating a scaling factor in the prior.* Similar to above, here we consider a special case of the mixture-of-normals prior in which the prior covariances are parameterized as $\boldsymbol{S}_{0k} = \sigma_0^2\boldsymbol{U}_k$, $\boldsymbol{U}_k \in \mathbb{S}_+^R$, $k = 1, \ldots, K$, $\sigma_0 \geqslant 0$. We assume the $\boldsymbol{U}_1, \ldots, \boldsymbol{U}_K$ are fixed parameters and we would like to estimate the scaling factor $\sigma_0$ by maximizing the likelihood:

$$\hat{\sigma}_0^2 := \mathrm{argmax}_{\sigma_0^2}\, p(\boldsymbol{Y} \mid \boldsymbol{X}, \boldsymbol{V}, \mathcal{S}_0, \boldsymbol{\omega}), \tag{65}$$

This is the same as maximizing the BF since the denominator in the BF does not depend on $\sigma_0$, and the BF may be more convenient to compute:

$$\hat{\sigma}_0^2 = \mathrm{argmax}_{\sigma_0^2}\, \mathrm{BF}^{\mathsf{mix}}(\boldsymbol{x}, \boldsymbol{Y}, \boldsymbol{V}, \mathcal{S}_0, \boldsymbol{\omega}). \tag{66}$$

Again taking a simple EM approach to computing the maximum-likelihood estimate, the M-step update allowing for singular matrices is

$$\sigma_0^2 = \sum_{k=1}^{K} \frac{\omega_{1k}}{R_k} \times \mathrm{tr}(\boldsymbol{U}_k^{\dagger}\mathbb{E}[\boldsymbol{b}\boldsymbol{b}^\intercal \mid \xi = k]), \tag{67}$$

in which the E-step involves computing the posterior probabilities $\omega_{1k}$ and the posterior second moments,

$$\mathbb{E}[\boldsymbol{b}\boldsymbol{b}^\intercal \mid \xi = k] = \boldsymbol{b}_{1k}\boldsymbol{b}_{1k}^\intercal + \boldsymbol{S}_{1k}. \tag{68}$$

Here, $R_k \leqslant R$ denotes the rank of $\boldsymbol{U}_k$.

#### *The multivariate single effect regression model*

The single effect regression (SER) model is a multiple regression model in which exactly one of the regression coefficients is non-zero [13]. The multivariate single effect regression (MSER) model extends the SER model to the multivariate setting, and forms the basis for mvSuSiE. The MSER model is simply a special case of the mvSuSiE model (2–5) with $L = 1$. The posterior distribution of $\boldsymbol{B}, \boldsymbol{\gamma}$ under the MSER model is summarized by the following proposition.

Proposition 4. Under the MSER model, the posterior distribution of $\boldsymbol{B}$ given the model parameters $\Theta := \{\boldsymbol{V}, \mathcal{S}_0, \boldsymbol{\omega}, \boldsymbol{\pi}\}$ is

$$\begin{aligned} \boldsymbol{\gamma} \mid \boldsymbol{X}, \boldsymbol{Y}, \Theta &\sim \mathrm{Multinom}(1, \boldsymbol{\alpha}) \\ \boldsymbol{b} \mid \boldsymbol{X}, \boldsymbol{Y}, \Theta, \gamma_j = 1 &\sim \sum_{k=1}^{K} \omega_{1jk} N_R(\boldsymbol{b}_{1jk}, \boldsymbol{S}_{1jk}), \end{aligned} \tag{69}$$

where $\boldsymbol{\alpha} = (\alpha_1, \ldots, \alpha_J)$ is the vector of posterior inclusion probabilities (PIPs), which can be written using the Bayes factors (54) for the Bayesian multivariate simple regression model with a mixture of normals prior,

$$\begin{aligned} \alpha_j &:= \alpha_j(\boldsymbol{x}_j, \boldsymbol{Y}, \Theta) \\ &= \Pr(\gamma_j = 1 \mid \boldsymbol{X}, \boldsymbol{Y}, \Theta) \\ &= \frac{\pi_j \mathrm{BF}^{\mathsf{mix}}(\boldsymbol{x}_j, \boldsymbol{Y}, \boldsymbol{V}, \mathcal{S}_0, \boldsymbol{\omega})}{\sum_{j'=1}^{J} \pi_{j'} \mathrm{BF}^{\mathsf{mix}}(\boldsymbol{x}_{j'}, \boldsymbol{Y}, \boldsymbol{V}, \mathcal{S}_0, \boldsymbol{\omega})}, \end{aligned} \tag{70}$$

and where the means $\boldsymbol{b}_{1jk}$, variances $\boldsymbol{S}_{1jk}$ and posterior mixture weights $\omega_{1jk}$ are given by

$$\boldsymbol{b}_{1jk} = \boldsymbol{b}_{1k}(\boldsymbol{x}_j, \boldsymbol{Y}, \boldsymbol{V}, \mathcal{S}_0, \boldsymbol{\omega}) \tag{71}$$

$$\boldsymbol{S}_{1jk} = \boldsymbol{S}_{1k}(\boldsymbol{x}_j, \boldsymbol{Y}, \boldsymbol{V}, \mathcal{S}_0, \boldsymbol{\omega}) \tag{72}$$

$$\omega_{1jk} = \omega_{1k}(\boldsymbol{x}_j, \boldsymbol{Y}, \boldsymbol{V}, \mathcal{S}_0, \boldsymbol{\omega}), \tag{73}$$

using the definitions of $\omega_{1k}$, $\boldsymbol{b}_{1k}$ and $\boldsymbol{S}_{1k}$ in (56–58). The posterior mean and covariance of $\boldsymbol{b}$ conditioned on $\boldsymbol{\gamma}$ are

$$\mathbb{E}[\boldsymbol{b} \mid \gamma_j = 1] = \boldsymbol{b}_1^{\mathsf{mix}}(\boldsymbol{x}_j, \boldsymbol{Y}, \boldsymbol{V}, \mathcal{S}_0, \boldsymbol{\omega}) \tag{74}$$

$$\mathrm{Cov}[\boldsymbol{b} \mid \gamma_j = 1] = \boldsymbol{S}_1^{\mathsf{mix}}(\boldsymbol{x}_j, \boldsymbol{Y}, \boldsymbol{V}, \mathcal{S}_0, \boldsymbol{\omega}), \tag{75}$$

using definitions (59, 60). Therefore, the posterior mean of $\boldsymbol{B}$ is

$$\mathbb{E}[\boldsymbol{B}] = \begin{bmatrix} \alpha_1 \boldsymbol{b}_1^{\mathsf{mix}}(\boldsymbol{x}_1, \boldsymbol{Y}, \boldsymbol{V}, \mathcal{S}_0, \boldsymbol{\omega})^\intercal \\ \vdots \\ \alpha_J \boldsymbol{b}_1^{\mathsf{mix}}(\boldsymbol{x}_J, \boldsymbol{Y}, \boldsymbol{V}, \mathcal{S}_0, \boldsymbol{\omega})^\intercal \end{bmatrix}. \tag{76}$$

For describing the algorithms below, we define a function, MSER, that returns the posterior distribution of $\boldsymbol{B}, \boldsymbol{\gamma}$ under the MSER model given the data $\boldsymbol{X}, \boldsymbol{Y}$ and the model parameters $\Theta$:

$$\mathrm{MSER}(\boldsymbol{X}, \boldsymbol{Y}; \Theta) := (\boldsymbol{\alpha}, \boldsymbol{\Omega}_1, \mathcal{B}_1, \mathcal{S}_1), \tag{77}$$

in which $\boldsymbol{\alpha} = (\alpha_1, \ldots, \alpha_J)$ is the vector of PIPs (70), $\boldsymbol{\Omega}_1$ is the $J \times K$ matrix of posterior mixture weights $\omega_{1jk}$ (73), $\mathcal{B}_1$ is the set of posterior means $\boldsymbol{b}_{1jk}$ (71) for all $j, k$, and $\mathcal{S}_1$ is the set of posterior covariances $\boldsymbol{S}_{1jk}$ (72) for all $j, k$.

*The multivariate single effect regression model with summary statistics.* From the above remarks, $\boldsymbol{X}^\intercal \boldsymbol{X}$ and $\boldsymbol{X}^\intercal \boldsymbol{Y}$ are sufficient to compute the posterior distribution of $\boldsymbol{B}, \boldsymbol{\gamma}$. (Note that only the diagonal elements of $\boldsymbol{X}^\intercal \boldsymbol{X}$ are actually needed to compute the posterior distribution of $\boldsymbol{B}, \gamma$.) To formalize these computations with summary statistics, we also define Bayes factors and posterior quantities in terms of summary statistics:

$$\boldsymbol{b}_{1jk}^{\mathsf{ss}} := \boldsymbol{b}_{1k}^{\mathsf{ss}}(\boldsymbol{x}_j^\intercal \boldsymbol{x}_j, \boldsymbol{Y}^\intercal \boldsymbol{x}_j, \boldsymbol{V}, \mathcal{S}_0, \boldsymbol{\omega}) \tag{78}$$

$$\boldsymbol{S}_{1jk}^{\mathsf{ss}} := \boldsymbol{S}_{1k}^{\mathsf{ss}}(\boldsymbol{x}_j^\intercal \boldsymbol{x}_j, \boldsymbol{Y}^\intercal \boldsymbol{x}_j, \boldsymbol{V}, \mathcal{S}_0, \boldsymbol{\omega}) \tag{79}$$

$$\omega_{1jk}^{\mathsf{ss}} := \omega_{1k}^{\mathsf{ss}}(\boldsymbol{x}_j^\intercal \boldsymbol{x}_j, \boldsymbol{Y}^\intercal \boldsymbol{x}_j, \boldsymbol{V}, \mathcal{S}_0, \boldsymbol{\omega}), \tag{80}$$

and the PIPs,

$$\alpha_j^{\mathsf{ss}} = \frac{\mathrm{BF}^{\mathsf{mix\text{-}ss}}(\boldsymbol{x}_j^\intercal \boldsymbol{x}_j, \boldsymbol{Y}^\intercal \boldsymbol{x}_j, \boldsymbol{V}, \mathcal{S}_0, \boldsymbol{\omega})}{\sum_{j'=1}^{J} \pi_{j'} \mathrm{BF}^{\mathsf{mix\text{-}ss}}(\boldsymbol{x}_{j'}^\intercal \boldsymbol{x}_{j'}, \boldsymbol{Y}^\intercal \boldsymbol{x}_{j'}, \boldsymbol{V}, \mathcal{S}_0, \boldsymbol{\omega})}. \tag{81}$$

To describe the algorithms that work with summary statistics, we define a function, MSER-ss, that returns the posterior distribution of $\boldsymbol{B}, \boldsymbol{\gamma}$ under the MSER model given the summary statistics $\boldsymbol{X}^\intercal \boldsymbol{X}, \boldsymbol{X}^\intercal \boldsymbol{Y}$ and the model parameters $\Theta$:

$$\text{MSER-ss}(\boldsymbol{X}^\intercal \boldsymbol{X}, \boldsymbol{X}^\intercal \boldsymbol{Y}; \Theta) := (\boldsymbol{\alpha}, \boldsymbol{\Omega}_1, \mathcal{B}_1, \mathcal{S}_1), \tag{82}$$

in which $\boldsymbol{\alpha} = (\alpha_1, \ldots, \alpha_J)$ is the vector of PIPs $\alpha_j^{\mathsf{ss}}$ (81), $\boldsymbol{\Omega}_1$ is the $J \times K$ matrix of posterior mixture weights $\omega_{1jk}^{\mathsf{ss}}$ (80), $\mathcal{B}_1$ is the set of posterior means $\boldsymbol{b}_{1jk}^{\mathsf{ss}}$ (78) for all $j, k$, and $\mathcal{S}_1$ is the set of posterior covariances $\boldsymbol{S}_{1jk}^{\mathsf{ss}}$ (79) for all $j, k$.

*Updating a scaling factor in the prior.* Here we extend the MSER model to allow for estimating a scaling parameter, $\sigma_0^2 \geqslant 0$, in which $\boldsymbol{S}_{0k} = \sigma_0^2 \boldsymbol{U}_k, k = 1, \ldots, K$. Similar to above, we estimate $\sigma_0^2$ by maximizing the likelihood,

$$\hat{\sigma}_0^2 := \mathrm{argmax}_{\sigma_0^2}\, p(\boldsymbol{Y} \mid \boldsymbol{X}, \Theta) \tag{83}$$

which is equivalent to maximizing a weighted sum of the Bayes factors,

$$\hat{\sigma}_0^2 = \underset{\sigma_0^2}{\operatorname{argmax}} \sum_{j=1}^{J} \pi_j \mathrm{BF}^{\mathsf{mix}}(\boldsymbol{x}_j, \boldsymbol{Y}, \boldsymbol{V}, \mathcal{S}_0, \boldsymbol{\omega}). \tag{84}$$

As before, the maximum-likelihood estimate can be computed using a simple EM algorithm. The M-step update in the EM algorithm is given by

$$\sigma_0^2 = \sum_{j=1}^{J} \sum_{k=1}^{K} \frac{\alpha_j \omega_{1jk}}{R_k} \times \operatorname{tr}(\boldsymbol{U}_k^{\dagger} \mathbb{E}[\boldsymbol{b}\boldsymbol{b}^{\mathsf{T}} \mid \gamma_j = 1, \xi_j = k]), \tag{85}$$

in which $R_k \leqslant R$ denotes the rank of $\boldsymbol{U}_k$. The E-step in the EM algorithm consists of computing the posterior inclusion probabilities $\alpha_j$, the posterior mixture weights $\omega_{1jk}$, and the posterior second moments

$$\mathbb{E}[\boldsymbol{b}\boldsymbol{b}^{\mathsf{T}} \mid \gamma_j = 1, \xi_j = k] = \boldsymbol{b}_{1jk}\boldsymbol{b}_{1jk}^{\mathsf{T}} + \boldsymbol{S}_{1jk} \tag{86}$$

at the current setting of $\sigma_0^2$.

#### *The mvSuSiE IBSS algorithm*

The Iterative Bayesian Stepwise Selection (IBSS) algorithm for fitting an mvSuSiE model extends the ideas of [13] to the multivariate setting. Similar to IBSS for SuSiE, the IBSS algorithm for mvSuSiE is a coordinate ascent algorithm for optimizing a variational approximation [36, 37] to the posterior distribution of $\boldsymbol{B}^{(1)}, \ldots, \boldsymbol{B}^{(L)}$ under an mvSuSiE model. The basic mvSuSiE IBSS algorithm is given in Algorithm 1. Lines 6–11 of the algorithm compute the posterior mean regression coefficients $\boldsymbol{b}^{(l)}$ for the $l$th single effect conditioned on each of the variables $j$ being the "single effect variable." These posteriors are then stored in a $J \times R$ matrix (lines 10–11). Once these are computed and stored, the unconditional posterior means $\bar{\boldsymbol{B}}^{(l)}$ are simply the conditional posterior means, $\boldsymbol{\mu}_l$, weighted by the PIPs $\boldsymbol{\alpha}_l$.

The optional step of estimating the scaling factors $\sigma_{0l}^2$ (line 5) is mainly for pruning unneeded single effects. This parameter estimation step can be viewed as an EM update in which the E-step is approximate.

#### *The IBSS algorithm for mvSuSiE with sufficient statistics*

Algorithm 2 describes the basic IBSS algorithm for mvSuSiE with sufficient statistics in which the computations are rearranged so that they only require the sufficient statistics $\boldsymbol{X}^{\mathsf{T}}\boldsymbol{Y}$ and $\boldsymbol{X}^{\mathsf{T}}\boldsymbol{X}$.

#### *Derivation of the mvSuSiE IBSS algorithm*

The IBSS algorithm fits an approximate posterior distribution for $\boldsymbol{B}^{(1)}, \ldots, \boldsymbol{B}^{(L)}$ by minimizing a Kullback-Leibler (K-L) divergence from the approximate posterior to the exact posterior subject to constraints on the approximate posterior. More precisely, denoting the exact posterior by $p_{\mathsf{post}}$ and the approximate posterior by $q$, we seek a $q$ that minimizes the K-L divergence $D_{\mathsf{KL}}(q \,\|\, p_{\mathsf{post}})$ [38] subject to constraints on $q$. Since the K-L divergence itself is hard to compute, we instead maximize the "evidence lower bound" (ELBO), which is equivalent to minimizing the K-L divergence, but more convenient to work with. The ELBO for the mvSuSiE model is

$$F(q; \boldsymbol{X}, \boldsymbol{Y}, \boldsymbol{V}) = \mathbb{E}_q[\log p(\boldsymbol{Y} \mid \boldsymbol{X}, \boldsymbol{B}^{(1)}, \ldots, \boldsymbol{B}^{(L)}, \boldsymbol{V})] + \mathbb{E}_q\left[\log\left\{\frac{p(\boldsymbol{B}^{(1)}, \ldots, \boldsymbol{B}^{(L)})}{q(\boldsymbol{B}^{(1)}, \ldots, \boldsymbol{B}^{(L)})}\right\}\right]. \tag{87}$$

Like [13], we constrain the approximate posterior so that it factorizes over the single effects:

$$q(\boldsymbol{B}^{(1)}, \ldots, \boldsymbol{B}^{(L)}) = \prod_{l=1}^{L} q_l(\boldsymbol{B}^{(l)}). \tag{88}$$

**Algorithm 1** Iterative Bayesian Stepwise Selection (IBSS) for mvSuSiE

**Require:** Data $\boldsymbol{X} \in \mathbb{R}^{N \times J}$, $\boldsymbol{Y} \in \mathbb{R}^{N \times R}$; upper limit on number of non-zero effects, $L \in \{1, \ldots, J\}$; initial estimates of the posterior mean single effects, $\bar{\boldsymbol{B}}^{(l)} \in \mathbb{R}^{J \times R}$, $l = 1, \ldots, L$; initial estimates of the prior scaling factors, $\sigma_{01}^2, \ldots, \sigma_{0L}^2 \geqslant 0$; a residual covariance matrix $\boldsymbol{V}$ (must be invertible); prior inclusion probabilities $\boldsymbol{\pi} = (\pi_1, \ldots, \pi_J)$; prior mixture weights $\boldsymbol{\omega} = (\omega_1, \ldots, \omega_K)$; prior covariance matrices $\mathcal{U} = (\boldsymbol{U}_1, \ldots, \boldsymbol{U}_K)$ (these do not need to be invertible); a function $\mathrm{MSER}(\boldsymbol{X}, \boldsymbol{Y}; \Theta) \rightarrow (\boldsymbol{\alpha}, \boldsymbol{\Omega}_1, \mathcal{B}_1, \mathcal{S}_1)$ that returns the posterior distribution of $\boldsymbol{B}, \boldsymbol{\gamma}$ under the MSER model with data $\boldsymbol{X}, \boldsymbol{Y}$ and parameters $\Theta$ (see eq. 77).

**repeat**
  $\bar{\boldsymbol{R}} \leftarrow \boldsymbol{Y} - \boldsymbol{X} \sum_{l=1}^{L} \bar{\boldsymbol{B}}^{(l)}$ ▻ Compute expected residuals.
  **for** $l$ in $1, \ldots, L$ **do**
    $\bar{\boldsymbol{R}}_l \leftarrow \bar{\boldsymbol{R}} + \boldsymbol{X} \bar{\boldsymbol{B}}^{(l)}$ ▻ Disregard $l$th single effect in residuals.
    Update $\sigma_{0l}^2$ ▻ Optional; see (85).
    $\Theta \leftarrow \{\boldsymbol{V}, \sigma_{0l}^2 \mathcal{U}, \boldsymbol{\omega}, \boldsymbol{\pi}\}$ ▻ Set MSER parameters.
    $(\boldsymbol{\alpha}, \boldsymbol{\Omega}_1, \mathcal{B}_1, \mathcal{S}_1) \leftarrow \mathrm{MSER}(\boldsymbol{X}, \bar{\boldsymbol{R}}_l; \Theta)$ ▻ Fit MSER to residuals.
    $\boldsymbol{\alpha}_l \leftarrow \boldsymbol{\alpha}$ ▻ Store PIPs for $l$th single effect.
    Initialize storage for $J \times R$ matrix $\boldsymbol{\mu}_l$ ▻ Compute conditional posterior means (74).
    **for** $j$ in $1, \ldots, J$ **do**
      Store $\sum_{k=1}^{K} \omega_{1jk} \boldsymbol{b}_{1jk}$ in row $j$ of $\boldsymbol{\mu}_l$
    $\bar{\boldsymbol{B}}^{(l)} \leftarrow (\boldsymbol{\alpha}_l \boldsymbol{1}_R^{\intercal}) \circ \boldsymbol{\mu}_l$ ▻ "$\circ$" denotes elementwise multiplication.
    $\bar{\boldsymbol{R}} \leftarrow \bar{\boldsymbol{R}}_l - \boldsymbol{X} \bar{\boldsymbol{B}}^{(l)}$ ▻ Update expected residuals.
**until** convergence criterion is met
**return** $\boldsymbol{\alpha}_1, \ldots, \boldsymbol{\alpha}_L, \boldsymbol{\mu}_1, \ldots, \boldsymbol{\mu}_L$

Under this constraint, $\boldsymbol{B}^{(1)}, \ldots, \boldsymbol{B}^{(L)}$ are independent *a posteriori*, and each factor $q_l(\boldsymbol{B}^{(l)})$ is a marginal posterior. The right-hand part of the ELBO now immediately simplifies:

$$F(q; \boldsymbol{X}, \boldsymbol{Y}, \boldsymbol{V}) = \mathbb{E}_q[\log p(\boldsymbol{Y} \mid \boldsymbol{X}, \boldsymbol{B}^{(1)}, \ldots, \boldsymbol{B}^{(L)}, \boldsymbol{V})] + \sum_{l=1}^{L} \mathbb{E}_q\left[\log\left\{\frac{p(\boldsymbol{B}^{(l)})}{q_l(\boldsymbol{B}^{(l)})}\right\}\right]. \tag{89}$$

And next we expanding the expected log-likelihood term on the left-hand side of this expression:

$$F(q; \boldsymbol{X}, \boldsymbol{Y}, \boldsymbol{V}) = -\tfrac{N}{2} \log|2\pi \boldsymbol{V}| - \tfrac{1}{2} \mathrm{tr}[\boldsymbol{V}^{-1} \mathrm{ERSS}(\boldsymbol{X}, \boldsymbol{Y}, \boldsymbol{B})] + \sum_{l=1}^{L} \mathbb{E}_q\left[\log\left\{\frac{p(\boldsymbol{B}^{(l)})}{q_l(\boldsymbol{B}^{(l)})}\right\}\right]. \tag{90}$$

Here, "ERSS" denotes the expected residual sum of squares,

$$\mathrm{ERSS}(\boldsymbol{X}, \boldsymbol{Y}, \boldsymbol{B}) := \mathbb{E}_q[(\boldsymbol{Y} - \boldsymbol{X}\boldsymbol{B})^{\intercal}(\boldsymbol{Y} - \boldsymbol{X}\boldsymbol{B})]. \tag{91}$$

In the following sections, we show that this simple conditional independence assumption (88) results in tractable computations that make up Algorithm 1. No other approximations or constraints are needed beyond this.

*Fitting the MSER to the residuals in Algorithm 1 maximizes the ELBO for a single effect.* The main result of this section is summarized by the following proposition.

Proposition 5. Let $F(q; \boldsymbol{X}, \boldsymbol{Y}, \boldsymbol{V})$ denote the ELBO (87) under the mvSuSiE model, and suppose that $q$ is constrained to factorize as in (88). The setting of $q_l(\boldsymbol{B}^{(l)})$ that maximizes the ELBO (87) while the other factors $q_{l'}(\boldsymbol{B}^{(l')})$ are fixed for all $l' \neq l$ has the following closed-form solution:

$$\hat{q}_l := \mathrm{argmax}_{q_l}\, F(q; \boldsymbol{X}, \boldsymbol{Y}, \boldsymbol{V}) = \mathrm{MSER}(\boldsymbol{X}, \bar{\boldsymbol{R}}_l; \Theta), \tag{92}$$

**Algorithm 2** IBSS for mvSuSiE with sufficient statistics (IBSS-ss)

**Require:** Data $\boldsymbol{X}^\intercal\boldsymbol{Y} \in \mathbb{R}^{J\times R}, \boldsymbol{X}^\intercal\boldsymbol{X} \in \mathbb{R}^{J\times J}$; maximum number of non-zero effects, $L \in \{1,\dots,J\}$; initial estimates of the posterior mean single effects, $\bar{\boldsymbol{B}}^{(l)} \in \mathbb{R}^{J\times R}$, $l = 1,\dots,L$; initial estimates of the prior scaling factors, $\sigma_{01}^2,\dots,\sigma_{0L}^2 \geqslant 0$; a residual covariance matrix $\boldsymbol{V}$ (must be invertible); prior inclusion probabilities $\boldsymbol{\pi} = (\pi_1,\dots,\pi_J)$; prior mixture weights $\boldsymbol{\omega} = (\omega_1,\dots,\omega_K)$; prior covariance matrices $\mathcal{U} = (\boldsymbol{U}_1,\dots,\boldsymbol{U}_K)$ (these do not need to be invertible); a function MSER-ss$(\boldsymbol{X}^\intercal\boldsymbol{X}, \boldsymbol{X}^\intercal\boldsymbol{Y};\Theta) \to (\boldsymbol{\alpha}, \boldsymbol{\Omega}_1, \mathcal{B}_1, \mathcal{S}_1)$ that returns the posterior distribution of $\boldsymbol{b}, \boldsymbol{\gamma}$ under the MSER model with data $\boldsymbol{X}^\intercal\boldsymbol{X}, \boldsymbol{X}^\intercal\boldsymbol{Y}$ and parameters $\Theta$ (see eq. 82).

**repeat**
  $\bar{\boldsymbol{P}} \leftarrow \boldsymbol{X}^\intercal\boldsymbol{Y} - \boldsymbol{X}^\intercal\boldsymbol{X}\sum_{l=1}^L \bar{\boldsymbol{B}}^{(l)}$ ▷ Compute expected residuals.
  **for** $l$ in $1,\dots,L$ **do**
    $\bar{\boldsymbol{P}}_l \leftarrow \bar{\boldsymbol{P}} + \boldsymbol{X}^\intercal\boldsymbol{X}\bar{\boldsymbol{B}}^{(l)}$ ▷ Disregard $l$th single effect in residuals.
    Update $\sigma_{0l}^2$ ▷ Optional; see (85).
    $\Theta \leftarrow \{\boldsymbol{V}, \sigma_{0l}^2\mathcal{U}, \boldsymbol{\omega}, \boldsymbol{\pi}\}$ ▷ Set MSER parameters.
    $(\boldsymbol{\alpha}, \boldsymbol{\Omega}_1, \mathcal{B}_1, \mathcal{S}_1) \leftarrow$ MSER-ss$(\boldsymbol{X}^\intercal\boldsymbol{X}, \bar{\boldsymbol{P}}_l;\Theta)$ ▷ Fit MSER to residuals.
    $\boldsymbol{\alpha}_l \leftarrow \boldsymbol{\alpha}$ ▷ Store PIPs for $l$th single effect.
    Initialize $\boldsymbol{\mu}_l$ to a $J \times R$ matrix of zeros ▷ Compute conditional posterior means (74).
    **for** $j$ in $1,\dots,J$ **do**
      Store $\sum_{k=1}^K \omega_{1jk}\boldsymbol{b}_{1jk}$ in row $j$ of $\boldsymbol{\mu}_l$
    $\bar{\boldsymbol{B}}^{(l)} \leftarrow (\boldsymbol{\alpha}_l\boldsymbol{1}_R^\intercal) \circ \boldsymbol{\mu}_l$ ▷ "$\circ$" denotes elementwise multiplication.
    $\bar{\boldsymbol{P}} \leftarrow \bar{\boldsymbol{P}}_l - \boldsymbol{X}^\intercal\boldsymbol{X}\bar{\boldsymbol{B}}^{(l)}$ ▷ Update expected residuals.
**until** convergence criterion is met
**return** $\boldsymbol{\alpha}_1,\dots,\boldsymbol{\alpha}_L, \boldsymbol{\mu}_1,\dots,\boldsymbol{\mu}_L$

in which MSER$(\boldsymbol{X},\boldsymbol{Y};\Theta)$ is defined in (77), and we further define $\bar{\boldsymbol{R}}_l$ as the $N \times R$ matrix of expected residuals that ignore the $l$th single effect, and $\bar{\boldsymbol{B}}^{(l)} := \mathbb{E}_q[\boldsymbol{B}^{(l)}]$:

$$\begin{aligned}\bar{\boldsymbol{R}}_l &:= \mathbb{E}_q[\boldsymbol{Y} - \boldsymbol{X}\textstyle\sum_{l'\neq l}\boldsymbol{B}^{(l')}] \\ &= \boldsymbol{Y} - \boldsymbol{X}\textstyle\sum_{l'\neq l}\bar{\boldsymbol{B}}^{(l')}.\end{aligned} \tag{93}$$

Corollary 1. A corollary of Proposition 5 is that the optimal $q(\boldsymbol{B})$ factorizing as (88) is a product of factors $q_l(\boldsymbol{B}^{(l)})$ in which each factor is the posterior distribution under an MSER model.

This proposition and the corollary are multivariate generalizations of results in [13] (Propositions 1 and A1 in that paper). The proof of this proposition is sketched out in the next sections.

*Special case when L = 1.* Recall, the MSER model is the mvSuSiE model with $L = 1$. For the MSER model, the ELBO (87) is

$$F_{\mathsf{MSER}}(q;\boldsymbol{X},\boldsymbol{Y},\boldsymbol{V}) = -\tfrac{N}{2}\log|2\pi\boldsymbol{V}| - \tfrac{1}{2}\mathrm{tr}[\boldsymbol{V}^{-1}\mathrm{ERSS}(\boldsymbol{X},\boldsymbol{Y},\boldsymbol{B}^{(1)})] - D_{\mathsf{KL}}(q\,\|\,p). \tag{94}$$

(In these expressions, we have dropped the "$l$" subscripts whenever they appear since there is only one single effect. However, we have kept the "$(l)$" superscript for $\boldsymbol{B}$ because $\boldsymbol{B}^{(1)}$ is not the same as $\boldsymbol{B}$.) The variational distribution maximizing this ELBO, that is, $\hat{q} := \mathrm{argmax}_q\, F_{\mathsf{MSER}}(q;\boldsymbol{X},\boldsymbol{Y},\boldsymbol{V})$, is the true posterior, $\hat{q}(\boldsymbol{B}^{(1)}) = p_{\mathsf{post}}(\boldsymbol{B}^{(1)}) := p(\boldsymbol{B}^{(1)} \mid \boldsymbol{X},\boldsymbol{Y},\boldsymbol{V})$. In other words, $\hat{q}$ is the exact posterior under the MSER model. Further, at $\hat{q}$ the ELBO (94) is equal to the marginal log-likelihood; that is, $F_{\mathsf{MSER}}(\hat{q};\boldsymbol{X},\boldsymbol{Y},\boldsymbol{V}) = \log p(\boldsymbol{Y} \mid \boldsymbol{X},\boldsymbol{V})$.

*Coordinate ascent update for $l$th single effect.* The fully-factorized contraint (88) allows for a divide-and-conquer approach; and therefore we now consider the problem of finding a $q_l(\boldsymbol{B}^{(l)})$ that maximizes the ELBO (87) while the remaining factors are fixed. Expanding terms involving $q_l$ only, the ELBO is

$$F(q;\boldsymbol{X},\boldsymbol{Y},\boldsymbol{V}) = -\tfrac{N}{2}\log|2\pi\boldsymbol{V}| - \tfrac{1}{2}\mathrm{tr}[\boldsymbol{V}^{-1}\mathrm{ERSS}(\boldsymbol{X},\boldsymbol{Y},\boldsymbol{B})] - D_{\mathsf{KL}}(q_l(\boldsymbol{B}^{(l)})\,\|\,p(\boldsymbol{B}^{(l)})) + \text{const}, \tag{95}$$

where the "const" is a placeholder for terms in the ELBO not involving $q_l$. As a reminder, $\boldsymbol{B} := \sum_{l=1}^{L} \boldsymbol{B}^{(l)}$. Expanding the ERSS further, and making use of the property from (88) that the covariances are zero between all $\boldsymbol{B}^{(l)}$ and $\boldsymbol{B}^{(l')}$ whenever $l \neq l'$, the ELBO can be rewritten as

$$F(q; \boldsymbol{X}, \boldsymbol{Y}, \boldsymbol{V}) = -\tfrac{N}{2} \log |2\pi \boldsymbol{V}| - \tfrac{1}{2}\mathrm{tr}[\boldsymbol{V}^{-1}\mathrm{ERSS}(\boldsymbol{X}, \bar{\boldsymbol{R}}_l, \boldsymbol{B}^{(l)})] - D_{\mathsf{KL}}(q_l(\boldsymbol{B}^{(l)}) \,\|\, p(\boldsymbol{B}^{(l)})) + \mathrm{const}, \quad (96)$$

where $\bar{\boldsymbol{R}}_l$ is defined in (93). For $l = 1$ without loss of generality, this expression is of the same form as (94) if we replace $\boldsymbol{Y}$ with $\bar{\boldsymbol{R}}_l$, and ignoring terms that do not involve $q_l$. In summary, the ELBO for mvSuSiE with $L > 1$ can be rearranged to exactly match the expression for the mvSuSiE ELBO with $L = 1$ if we ignore terms not involving $q_l$.

#### *Computing the ELBO*

While computing the mvSuSiE ELBO (87) is not strictly needed to implement the IBSS algorithm, in practice it is useful for monitoring progress of the IBSS algorithm and for comparing different mvSuSiE model fits.

Expanding the ERSS part of the ELBO, we obtain

$$\mathrm{ERSS}(\boldsymbol{X}, \boldsymbol{Y}, \boldsymbol{B}) = (\boldsymbol{Y} - \boldsymbol{X}\bar{\boldsymbol{B}})^\intercal(\boldsymbol{Y} - \boldsymbol{X}\bar{\boldsymbol{B}}) - \textstyle\sum_{l=1}^{L}(\bar{\boldsymbol{B}}^{(l)})^\intercal \boldsymbol{X}^\intercal \boldsymbol{X} \bar{\boldsymbol{B}}^{(l)} + \mathbb{E}_q[(\boldsymbol{B}^{(l)})^\intercal \boldsymbol{X}^\intercal \boldsymbol{X} \boldsymbol{B}^{(l)}]. \quad (97)$$

In the mvSuSiE model, for a given $l$ only one row of $\boldsymbol{B}^{(l)}$ contains nonzero values, so $\mathbb{E}_q[b_{jr} b_{kr'}] = b_{jr} b_{kr'} = 0$ for any $j \neq k$ and $r, r' \in \{1, \ldots, R\}$, where $b_{jr}$ is an entry of $\boldsymbol{B}^{(l)}$. With this property, the right-most term in the ERSS becomes

$$\mathbb{E}_q[(\boldsymbol{B}^{(l)})^\intercal \boldsymbol{X}^\intercal \boldsymbol{X} \boldsymbol{B}^{(l)}] = \sum_{l=1}^{L} (\bar{\boldsymbol{B}}^{(l)})^\intercal \boldsymbol{D} \bar{\boldsymbol{B}}^{(l)} + \sum_{l=1}^{L} \sum_{j=1}^{J} d_j \boldsymbol{C}_j^{(l)}, \quad (98)$$

where $\boldsymbol{D}$ is a $J \times J$ diagonal matrix with diagonal entries $d_j = (\boldsymbol{X}^\intercal \boldsymbol{X})_{jj}$, and $\boldsymbol{C}_j^{(l)}$ is the $R \times R$ covariance matrix for the $j$th row of $\boldsymbol{B}^{(l)}$ with respect to the approximate posterior $q_l$. This covariance matrix is easily computed from Proposition 4. The remaining terms in the ELBO are K-L divergences. It is convenient to compute these terms whenever $q_l(\boldsymbol{B}^{(l)})$ is updated, as we show next.

*Computing the K-L divergence when L = 1.* The K-L divergence term in the MSER ELBO (94) satisfies the following identity:

$$\begin{aligned} D_{\mathsf{KL}}(q \,\|\, p) &= \mathbb{E}_q\left[\log\left\{\frac{q(\boldsymbol{B}^{(1)})}{p(\boldsymbol{B}^{(1)})}\right\}\right] \\ &= -\tfrac{N}{2}\log|2\pi\boldsymbol{V}| - \tfrac{1}{2}\mathrm{tr}[\boldsymbol{V}^{-1}\mathrm{ERSS}(\boldsymbol{X}, \boldsymbol{Y}, \boldsymbol{B}^{(1)})] - F_{\mathsf{MSER}}(q; \boldsymbol{X}, \boldsymbol{Y}, \boldsymbol{V}). \end{aligned} \quad (99)$$

Recall, the optimal variational distribution is the true posterior, $\hat{q}(\boldsymbol{B}^{(1)}) = p_{\mathsf{post}}(\boldsymbol{B}^{(1)})$. And when the optimal variational distribution is attained, the ELBO is equal to the marginal log-likelihood. Therefore, at $\hat{q}$ the MSER ELBO is

$$\begin{aligned} D_{\mathsf{KL}}(\hat{q} \,\|\, p) &= -\tfrac{N}{2}\log|2\pi\boldsymbol{V}| - \tfrac{1}{2}\mathrm{tr}[\boldsymbol{V}^{-1}\mathrm{ERSS}(\boldsymbol{X}, \boldsymbol{Y}, \boldsymbol{B}^{(1)})] - \log p(\boldsymbol{Y} \mid \boldsymbol{X}, \boldsymbol{V}) \\ &= -\tfrac{N}{2}\log|2\pi\boldsymbol{V}| - \tfrac{1}{2}\mathrm{tr}[\boldsymbol{V}^{-1}\mathrm{ERSS}(\boldsymbol{X}, \boldsymbol{Y}, \boldsymbol{B}^{(1)})] - \log \textstyle\sum_{j=1}^{J} \mathrm{BF}^{\mathsf{mix}}(\boldsymbol{x}_j, \boldsymbol{Y}, \boldsymbol{V}, \sigma_0^2\mathcal{U}, \boldsymbol{\omega}) \\ &\quad - \log p(\boldsymbol{Y} \mid \boldsymbol{X}, \boldsymbol{V}, \boldsymbol{B} = \boldsymbol{0}) \\ &= \tfrac{1}{2}\mathrm{tr}(\boldsymbol{V}^{-1}\boldsymbol{Y}^\intercal\boldsymbol{Y}) - \tfrac{1}{2}\mathrm{tr}[\boldsymbol{V}^{-1}\mathrm{ERSS}(\boldsymbol{X}, \boldsymbol{Y}, \boldsymbol{B}^{(1)})] - \log \textstyle\sum_{j=1}^{J} \mathrm{BF}^{\mathsf{mix}}(\boldsymbol{x}_j, \boldsymbol{Y}, \boldsymbol{V}, \sigma_0^2\mathcal{U}, \boldsymbol{\omega}), \end{aligned} \quad (100)$$

in which we have defined $\sigma_0^2\mathcal{U}$ as the set of prior covariance matrices scaled by $\sigma_0^2$, $\sigma_0^2\mathcal{U} := \{\sigma_0^2\boldsymbol{U}_1, \ldots, \sigma_0^2\boldsymbol{U}_K\}$. Finally, to arrive at the desired K-L divergence $D_{\mathsf{KL}}(\hat{q}_l \,\|\, p_l)$ for the mvSuSiE ELBO, we substitute $\bar{\boldsymbol{R}}_l$ for $\boldsymbol{Y}$ in (100). A similar approach to computing the ELBO was taken in [13].

### References

[1] Zou, Y., P. Carbonetto, G. Wang, and M. Stephens (2022). Fine-mapping from summary data with the "Sum of Single Effects" model. *PLoS Genetics 18*(7), e1010299.

[2] Urbut, S. M., G. Wang, P. Carbonetto, and M. Stephens (2019). Flexible statistical methods for estimating and testing effects in genomic studies with multiple conditions. *Nature Genetics 51*, 187–195.

[3] Pirinen, M., P. Donnelly, and C. C. A. Spencer (2012). Including known covariates can reduce power to detect genetic effects in case-control studies. *Nature Genetics 44*(8), 848–851.

[4] Pirinen, M., P. Donnelly, and C. C. A. Spencer (2013). Efficient computation with a linear mixed model on large-scale data sets with applications to genetic studies. *Annals of Applied Statistics 7*(1), 369–390.

[5] Little, R. J. A. and D. B. Rubin (2020). *Statistical analysis with missing data* (third ed.). Hoboken, NJ: John Wiley & Sons, Inc.

[6] Giambartolomei, C., D. Vukcevic, E. E. Schadt, L. Franke, A. D. Hingorani, C. Wallace, and V. Plagnol (2014). Bayesian test for colocalisation between pairs of genetic association studies using summary statistics. *PLoS Genetics 10*(5), e1004383.

[7] LaPierre, N., K. Taraszka, H. Huang, R. He, F. Hormozdiari, and E. Eskin (2021). Identifying causal variants by fine mapping across multiple studies. *PLoS Genetics 17*(9), e1009733.

[8] Lu, Z., X. Wang, M. Carr, A. Kim, S. Gazal, P. Mohammadi, L. Wu, J. Pirruccello, L. Kachuri, A. Gusev, and N. Mancuso (2025). Improved multiancestry fine-mapping identifies cis-regulatory variants underlying molecular traits and disease risk. *Nature Genetics 57*(8), 1881–1889.

[9] Kanai, M., R. Elzur, W. Zhou, M. Kanai, K.-H. H. Wu, et al. (2022). Meta-analysis fine-mapping is often miscalibrated at single-variant resolution. *Cell Genomics 2*(12), 100210.

[10] Sarsani, V., S. M. Brotman, Y. Xianyong, L. Fernandes Silva, M. Laakso, and C. N. Spracklen (2024). A cross-ancestry genome-wide meta-analysis, fine-mapping, and gene prioritization approach to characterize the genetic architecture of adiponectin. *Human Genetics and Genomics Advances 5*(1), 100252.

[11] Yuan, K., R. J. Longchamps, A. F. Pardiñas, M. Yu, T.-T. Chen, S.-C. Lin, Y. Chen, M. Lam, R. Liu, Y. Xia, Z. Guo, W. Shi, C. Shen, M. J. Daly, B. M. Neale, Y.-C. A. Feng, Y.-F. Lin, C.-Y. Chen, M. C. O'Donovan, T. Ge, and H. Huang (2024). Fine-mapping across diverse ancestries drives the discovery of putative causal variants underlying human complex traits and diseases. *Nature Genetics 56*(9), 1841–1850.

[12] Zhang, X., W. Jiang, and H. Zhao (2024). Integration of expression QTLs with fine mapping via SuSiE. *PLoS Genetics 20*(1), e1010929.

[13] Wang, G., A. Sarkar, P. Carbonetto, and M. Stephens (2020). A simple new approach to variable selection in regression, with application to genetic fine mapping. *Journal of the Royal Statistical Society, Series B 82*(5), 1273–1300.

[14] Clyde, M. A., J. Ghosh, and M. L. Littman (2011). Bayesian adaptive sampling for variable selection and model averaging. *Journal of Computational and Graphical Statistics 20*(1), 80–101.

[15] Dellaportas, P., J. J. Forster, and I. Ntzoufras (2002). On Bayesian model and variable selection using MCMC. *Statistics and Computing 12*, 27–36.

[16] George, E. I. and R. E. McCulloch (1993). Variable selection via Gibbs sampling. *Journal of the American Statistical Association 88*(423), 881–889.

[17] Hoggart, C. J., J. C. Whittaker, M. De Iorio, and D. J. Balding (2008). Simultaneous analysis of all SNPs in genome-wide and re-sequencing association studies. *PLoS Genetics 4*(7), e1000130.

[18] Guan, Y. and M. Stephens (2011). Bayesian variable selection regression for genome-wide association studies and other large-scale problems. *Annals of Applied Statistics 5*(3), 1780–1815.

[19] Moser, G., S. H. Lee, B. J. Hayes, M. E. Goddard, N. R. Wray, and P. M. Visscher (2015). Simultaneous discovery, estimation and prediction analysis of complex traits using a Bayesian mixture model. *PLoS Genetics 11*(4), e1004969.

[20] Zanella, G. and G. Roberts (2019). Scalable importance tempering and Bayesian variable selection. *Journal of the Royal Statistical Society, Series B 81*(3), 489–517.

[21] Zhou, X., P. Carbonetto, and M. Stephens (2013). Polygenic modeling with Bayesian sparse linear mixed models. *PLoS Genetics 9*(2), e1003264.

[22] Neal, R. M. (1996). *Bayesian learning for neural networks*, Volume 118 of *Lecture Notes in Statistics*. New York, NY: Springer.

[23] Tipping, M. E. (2001). Sparse Bayesian learning and the Relevance Vector Machine. *Journal of Machine Learning Research 1*, 211–244.

[24] Chang, C. C., C. C. Chow, L. C. Tellier, S. Vattikuti, S. M. Purcell, and J. J. Lee (2015). Second-generation PLINK: rising to the challenge of larger and richer datasets. *Gigascience 4*(1), s13742–015–0047–8.

[25] Benner, C., A. S. Havulinna, M.-R. Järvelin, V. Salomaa, S. Ripatti, and M. Pirinen (2017). Prospects of fine-mapping trait-associated genomic regions by using summary statistics from genome-wide association studies. *American Journal of Human Genetics 101*(4), 539–551.

[26] Stephens, M. and D. J. Balding (2009). Bayesian statistical methods for genetic association studies. *Nature Reviews Genetics 10*(10), 681–690.

[27] Wakefield, J. (2009). Bayes factors for genome-wide association studies: comparison with P-values. *Genetic Epidemiology 33*(1), 79–86.

[28] Stephens, M. (2017). False discovery rates: a new deal. *Biostatistics 18*(2), 275–294.

[29] Maller, J. B., G. McVean, J. Byrnes, D. Vukcevic, K. Palin, et al. (2012). Bayesian refinement of association signals for 14 loci in 3 common diseases. *Nature Genetics 44*(12), 1294–1301.

[30] Wang, W. and M. Stephens (2021). Empirical Bayes matrix factorization. *Journal of Machine Learning Research 22*(120), 1–40.

[31] Bovy, J., D. W. Hogg, and S. T. Roweis (2011). Extreme Deconvolution: inferring complete distribution functions from noisy, heterogeneous and incomplete observations. *Annals of Applied Statistics 5*(2B), 1657–1677.

[32] Dawid, A. P. (1981). Some matrix-variate distribution theory: notational considerations and a Bayesian application. *Biometrika 68*(1), 265–274.

[33] Gupta, A. and D. Nagar (2000). *Matrix variate distributions*. Boca Raton, FL: Chapman & Hall.

[34] Dempster, A. P., N. M. Laird, and D. B. Rubin (1977). Maximum likelihood from incomplete data via the EM algorithm. *Journal of the Royal Statistical Society, Series B 39*(1), 1–22.

[35] Chipman, H., E. I. George, and R. E. McCulloch (2001). The practical implementation of Bayesian model selection. In P. Lahiri (Ed.), *Model Selection*, Volume 38 of *IMS Lecture Notes*, pp. 65–116. Beachwood, OH: Institute of Mathematical Statistics.

[36] Blei, D. M., A. Kucukelbir, and J. D. McAuliffe (2017). Variational inference: a review for statisticians. *Journal of the American Statistical Association 112*(518), 859–877.

[37] Jordan, M. I., Z. Ghahramani, T. S. Jaakkola, and L. K. Saul (1999). An introduction to variational methods for graphical models. *Machine Learning 37*(2), 183–233.

[38] Cover, T. M. and J. A. Thomas (2006). *Elements of Information Theory* (2nd ed.). Wiley-Interscience.

[39] Schmidt, E. M., J. Zhang, W. Zhou, J. Chen, K. L. Mohlke, Y. E. Chen, and C. J. Willer (2015). GREGOR: evaluating global enrichment of trait-associated variants in epigenomic features using a systematic, data-driven approach. *Bioinformatics 31*(16), 2601–2606.

[40] Finucane, H. K., B. Bulik-Sullivan, A. Gusev, G. Trynka, Y. Reshef, P. R. Loh, V. Anttila, H. Xu, C. Zang, K. Farh, S. Ripke, F. R. Day, S. Purcell, E. Stahl, S. Lindstrom, J. R. Perry, Y. Okada, S. Raychaudhuri, M. J. Daly, N. Patterson, B. M. Neale, and A. L. Price (2015). Partitioning heritability by functional annotation using genome-wide association summary statistics. *Nature Genetics 47*(11), 1228–1235.

[41] Gusev, A., S. H. Lee, G. Trynka, H. Finucane, B. J. Vilhjálmsson, H. Xu, C. Zang, S. Ripke, B. Bulik-Sullivan, E. Stahl, A. K. Kähler, C. M. Hultman, S. M. Purcell, S. A. McCarroll, M. Daly, B. Pasaniuc, P. F. Sullivan, B. M. Neale, N. R. Wray, S. Raychaudhuri, and A. L. Price (2014).

Partitioning heritability of regulatory and cell-type-specific cariants across 11 common diseases. *American Journal of Human Genetics 95*(5), 535–552.

[42] Vasquez, Y. M., E. C. Mazur, X. Li, R. Kommagani, L. Jiang, R. Chen, R. B. Lanz, E. Kovanci, W. E. Gibbons, and F. J. DeMayo (2015). FOXO1 is required for binding of PR on IRF4, novel transcriptional regulator of endometrial stromal decidualization. *Molecular Endocrinology 29*(3), 421–433.

[43] Hormozdiari, F., S. Gazal, B. van de Geijn, H. K. Finucane, C. J. Ju, P.-R. Loh, A. Schoech, Y. Reshef, X. Liu, L. O'Connor, A. Gusev, E. Eskin, and A. L. Price (2018). Leveraging molecular quantitative trait loci to understand the genetic architecture of diseases and complex traits. *Nature Genetics 50*(7), 1041–1047.

[44] Ulirsch, J. C., C. A. Lareau, E. L. Bao, L. S. Ludwig, M. H. Guo, C. Benner, A. T. Satpathy, V. K. Kartha, R. M. Salem, J. N. Hirschhorn, H. K. Finucane, M. J. Aryee, J. D. Buenrostro, and V. G. Sankaran (2019). Interrogation of human hematopoiesis at single-cell and single-variant resolution. *Nature Genetics 51*(4), 683–693.

[45] Vuckovic, D., E. L. Bao, P. Akbari, C. A. Lareau, A. Mousas, et al. (2020). The polygenic and monogenic basis of blood traits and diseases. *Cell 182*(5), 1214–1231.e11.

### Supplementary figures

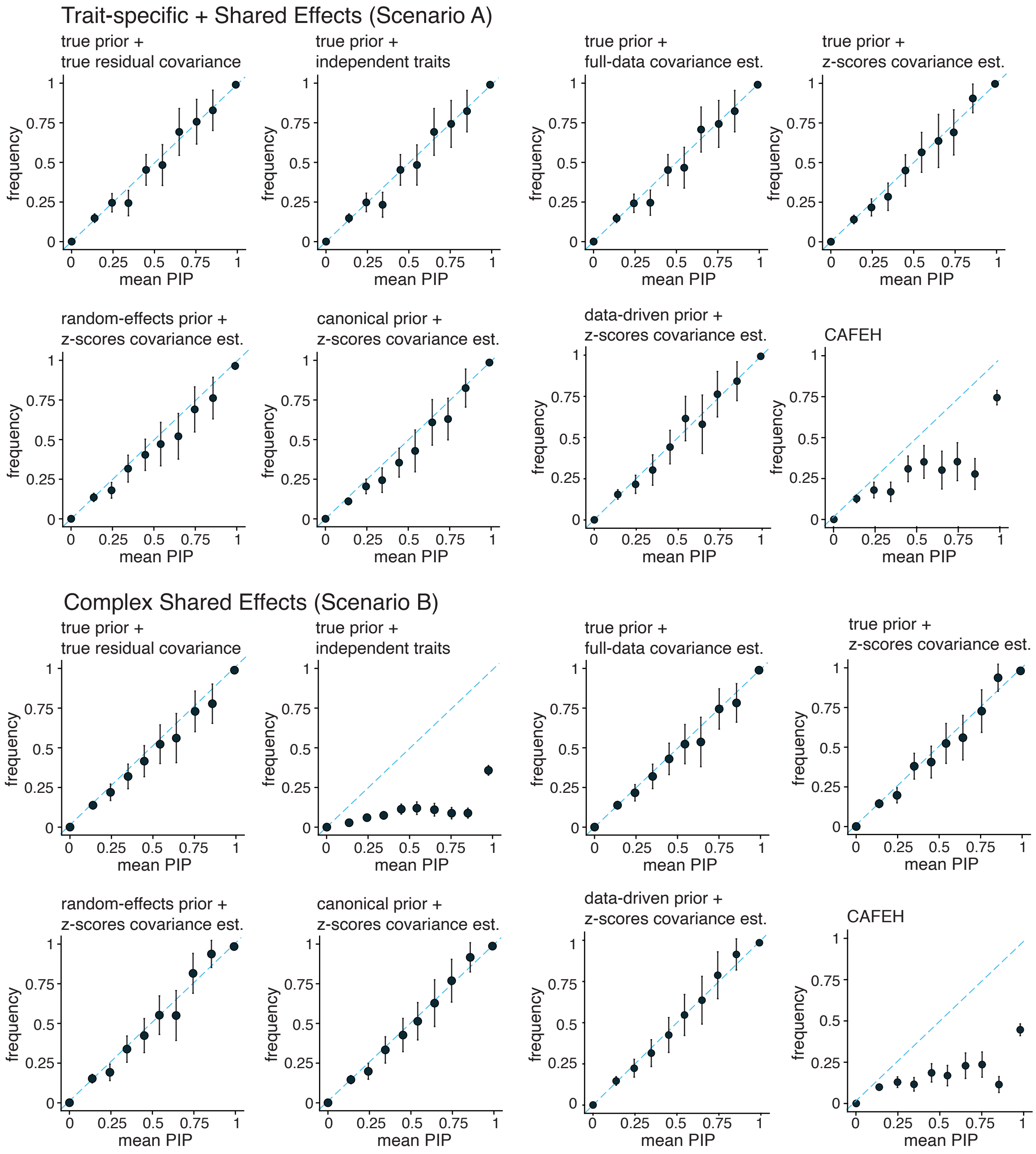

**Supplementary Figure 1. Assessment of mvSuSiE and CAFEH PIP calibration.** In each scenario, SNPs from all simulations ($n$ = 600) were grouped into bins according to their reported PIP (10 equally spaced bins from 0 to 1).The plots show the average PIP from each bin (X axis) against the proportion of SNPs in that bin that are causal (Y axis). For a given bin, the error bar depicts 2 times the empirical s.e. from all $n$ = 600 simulations. A well-calibrated method should produce points near the diagonal. See also Supplementary Figures 2 and 3.

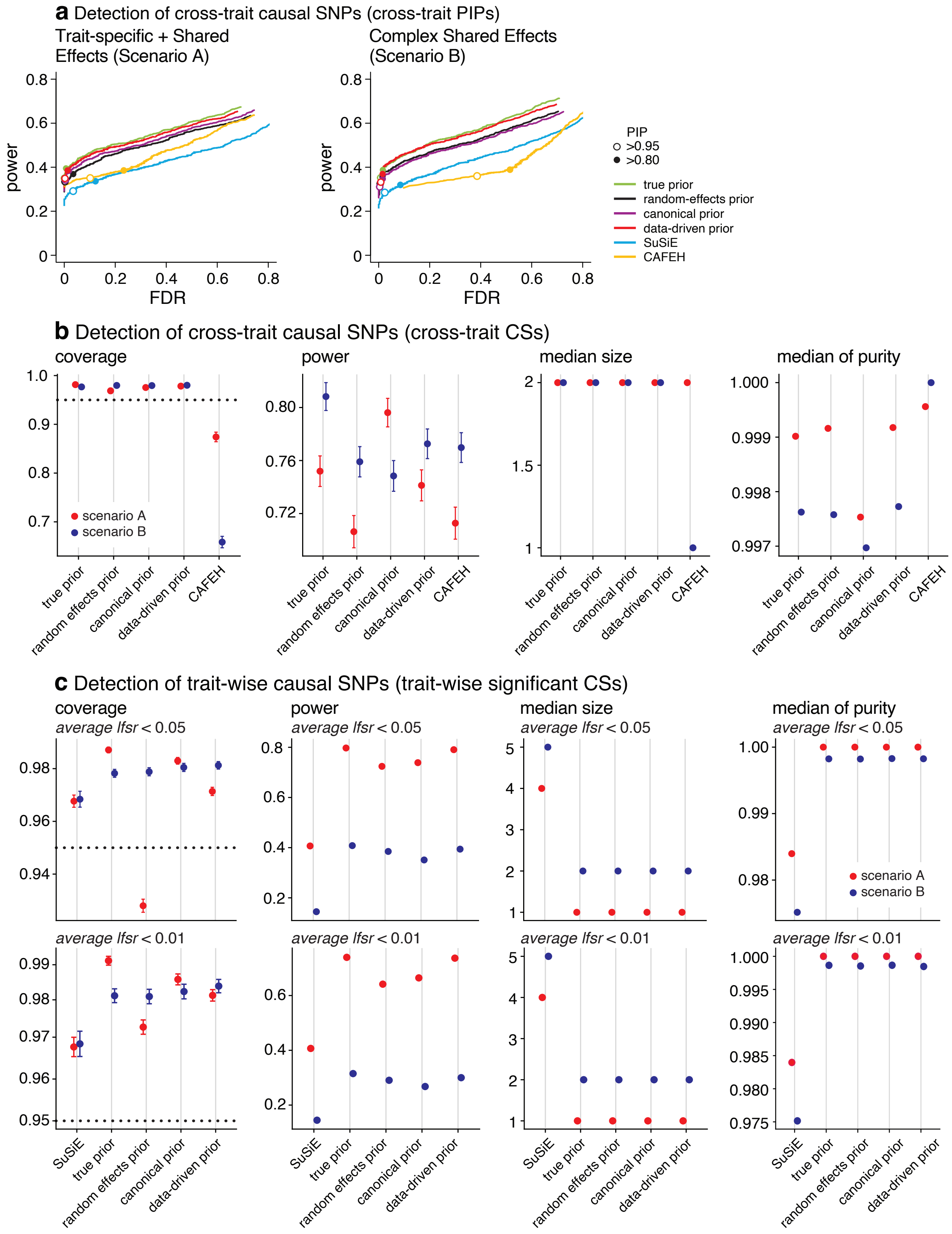

**Supplementary Figure 2. Comparison of mvSuSiE with different priors.** See legend on next page.

**Supplementary Figure 2 (previous page). Comparison of mvSuSiE with different priors.** This figure is a companion to Fig. 2 giving more detail about the performance of mvSuSiE with different priors. In addition to SuSiE and CAFEH, four variants of mvSuSiE were compared: mvSuSiE with a "random effects" prior; mvSuSiE with a "canonical" prior; mvSuSiE with a "data-driven" prior; and mvSuSiE with the true prior, i.e., the prior used to simulate the data. See the Fig. 2 caption for additional explanations of the plots. Note that for the power/FDR results at specific thresholds, some of the results are not visible in the plots because some circles lie directly on top of each other. Detailed power/FDR statistics at the selected thresholds are given in Supplementary Table 1.

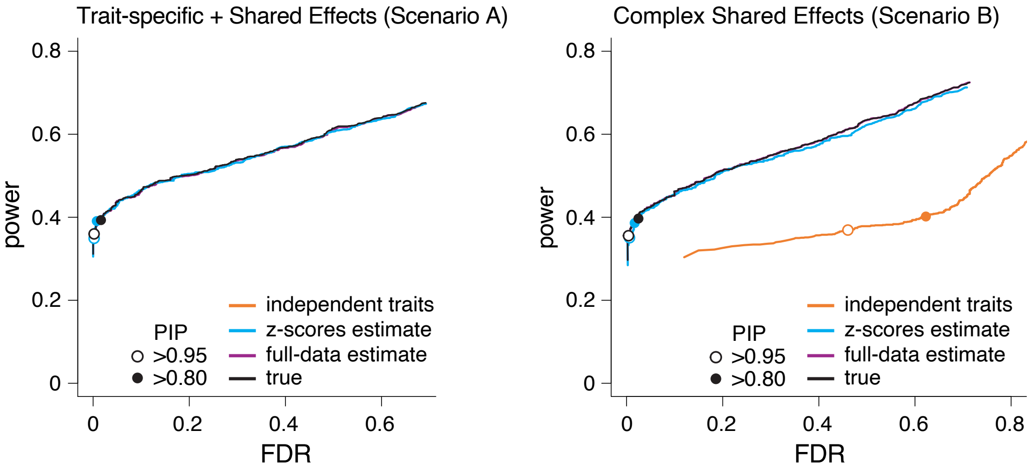

**Supplementary Figure 3. Comparison of mvSuSiE with different residual covariances.** mvSuSiE with following choices of $\boldsymbol{V}$ were compared: identity matrix ("independent traits"); $\boldsymbol{V}$ estimated from the individual-level data ("full-data estimate"); $\boldsymbol{V}$ estimated from the *z*-scores ("*z*-scores estimate"); and the $\boldsymbol{V}$ used to simulate the data ("true"). Note that in Scenario A, the true residual covariance matrix was the identity matrix, so the results are exactly the same for both "true" and "independent traits." FDR and power were calculated as the cross-trait PIP was varied from 0 to 1 ($n$ = 600 simulations). Power and FDR at specific thresholds are indicated by the circles. Note that the thresholds shown for the different variants of mvSuSiE are not necessarily equivalent or comparable; results are shown at these thresholds for illustration only. Also note that some of the power/FDR results are not visible in the plots because the circles lie directly underneath others. Detailed power/FDR statistics at the selected thresholds are given in Supplementary Table 1. For these comparisons only, the prior was set to the mixture distribution used to simulate the effects. In Scenario A, in which the traits were simulated with $\boldsymbol{V} = \boldsymbol{I}_R$, there was almost no difference in assuming or not assuming $\boldsymbol{V} = \boldsymbol{I}_R$ in the mvSuSiE analysis; all the power-FDR curves closely overlap in the plot. In the right-hand plot (Scenario B), $\boldsymbol{Y}$ was simulated with $\boldsymbol{V} \neq \boldsymbol{I}_R$, and in that case there was a substantial reduction in performance when $\boldsymbol{V} = \boldsymbol{I}_R$ ("independent traits").

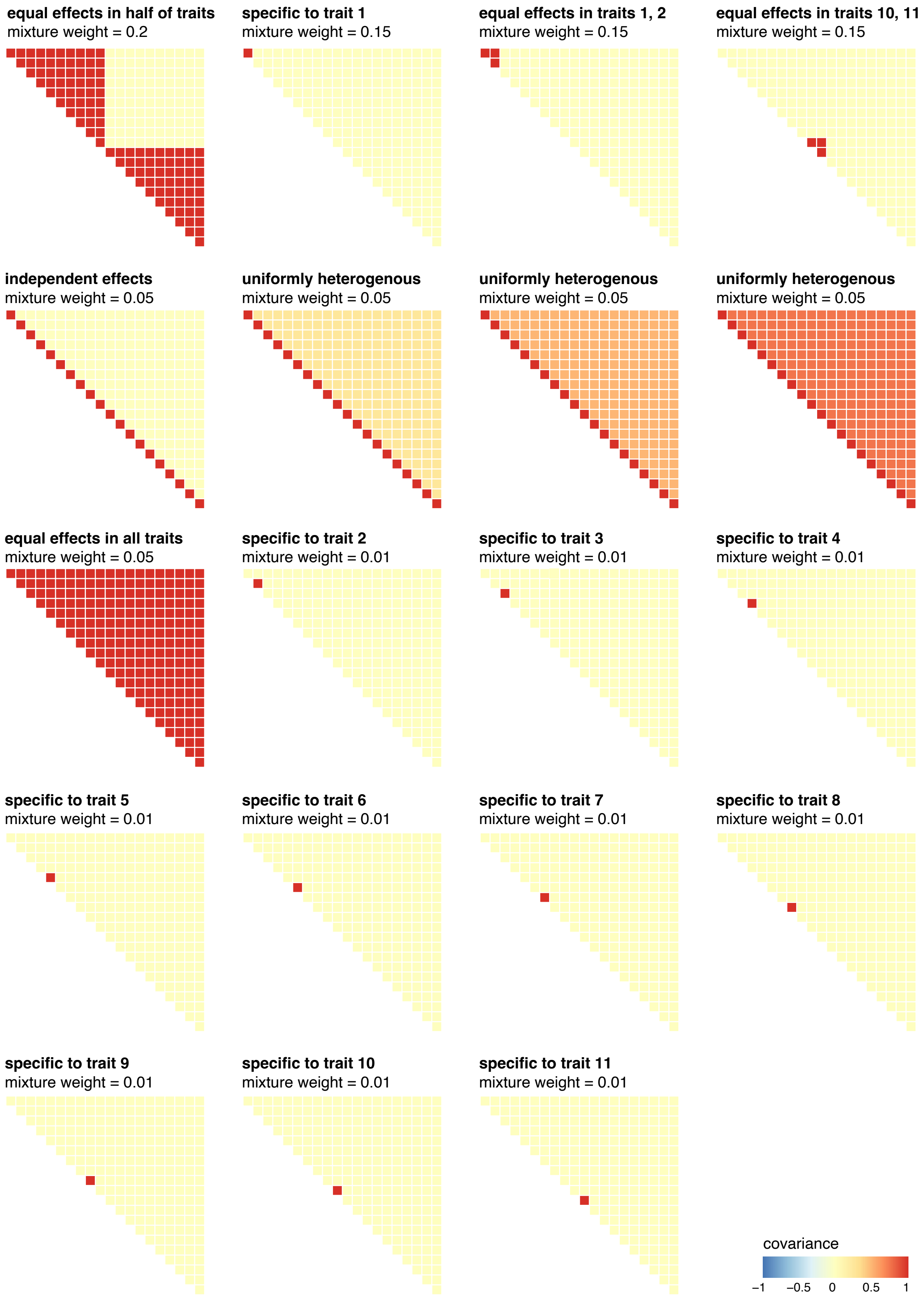

**Supplementary Figure 4. Covariance matrices used to simulate the effects of the causal SNPs in Scenario A.** Each plot shows a 20 × 20 covariance matrix, $U_k$, and its corresponding mixture weight, $\omega_k$, in the mixture-of-multivariate normals distribution used to simulate the effects of the causal SNPs. Note that all of these covariance matrices contain elements spanning the range 0 to 1.

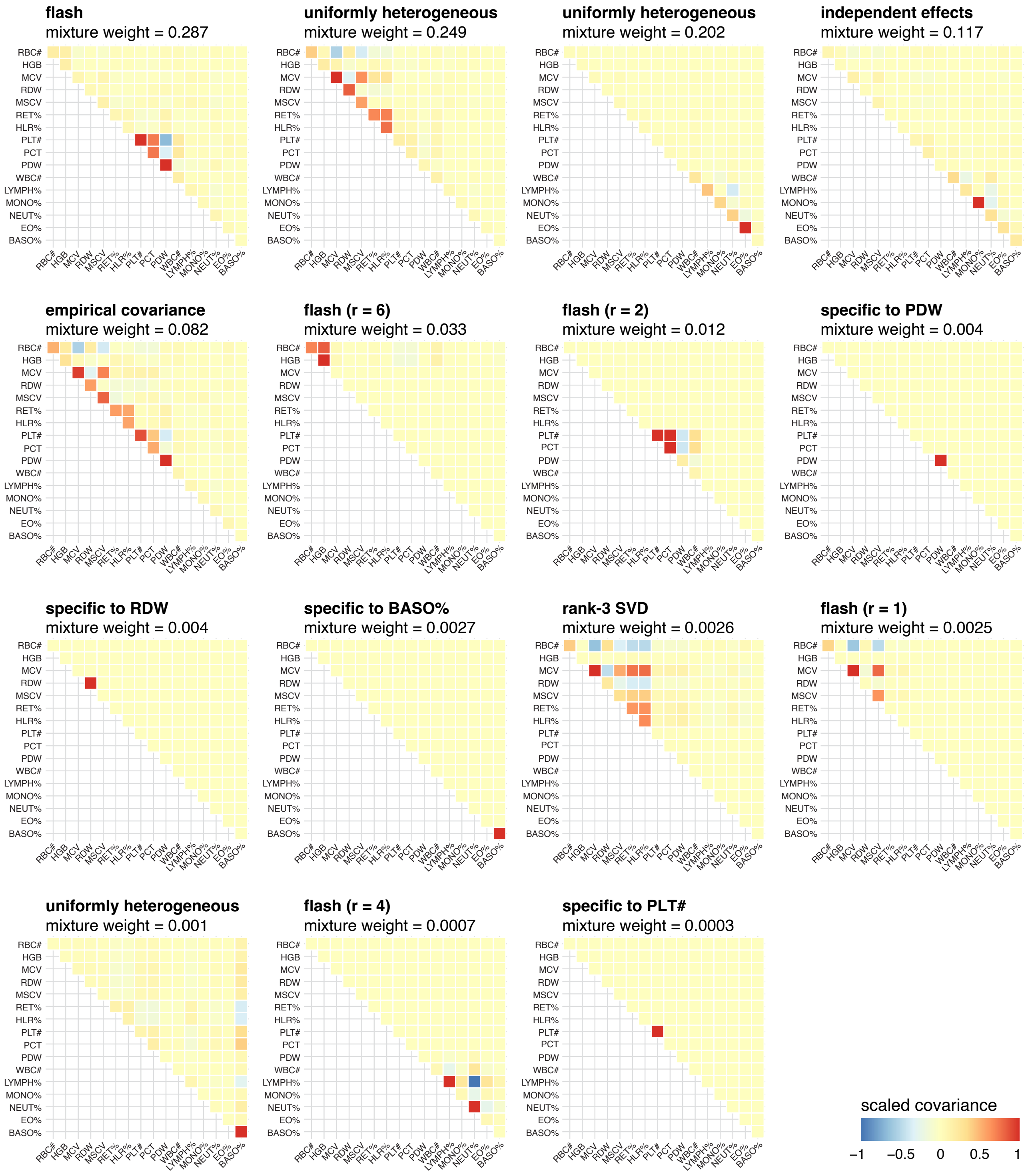

**Supplementary Figure 5. Prior on multivariate SNP effects estimated from the UK Biobank blood cell traits.** Each plot shows a 16 × 16 scaled covariance matrix $\boldsymbol{U}_k$ and its corresponding estimated mixture weight $\omega_k$. These covariance matrices and mixture weights specify the prior used in the mvSuSiE analyses of the UK Biobank blood cell traits. For visualization purposes only, each plot shows the scaled covariance matrix $\boldsymbol{U}_k/s_k^2$, where $s_k^2$ is the absolute value of the largest (in magnitude) entry of $\boldsymbol{U}_k$, so that all of the plotted values lie between -1 and 1. Each covariance matrix is labeled by the method used to initialize the estimate (see Supplementary Note).

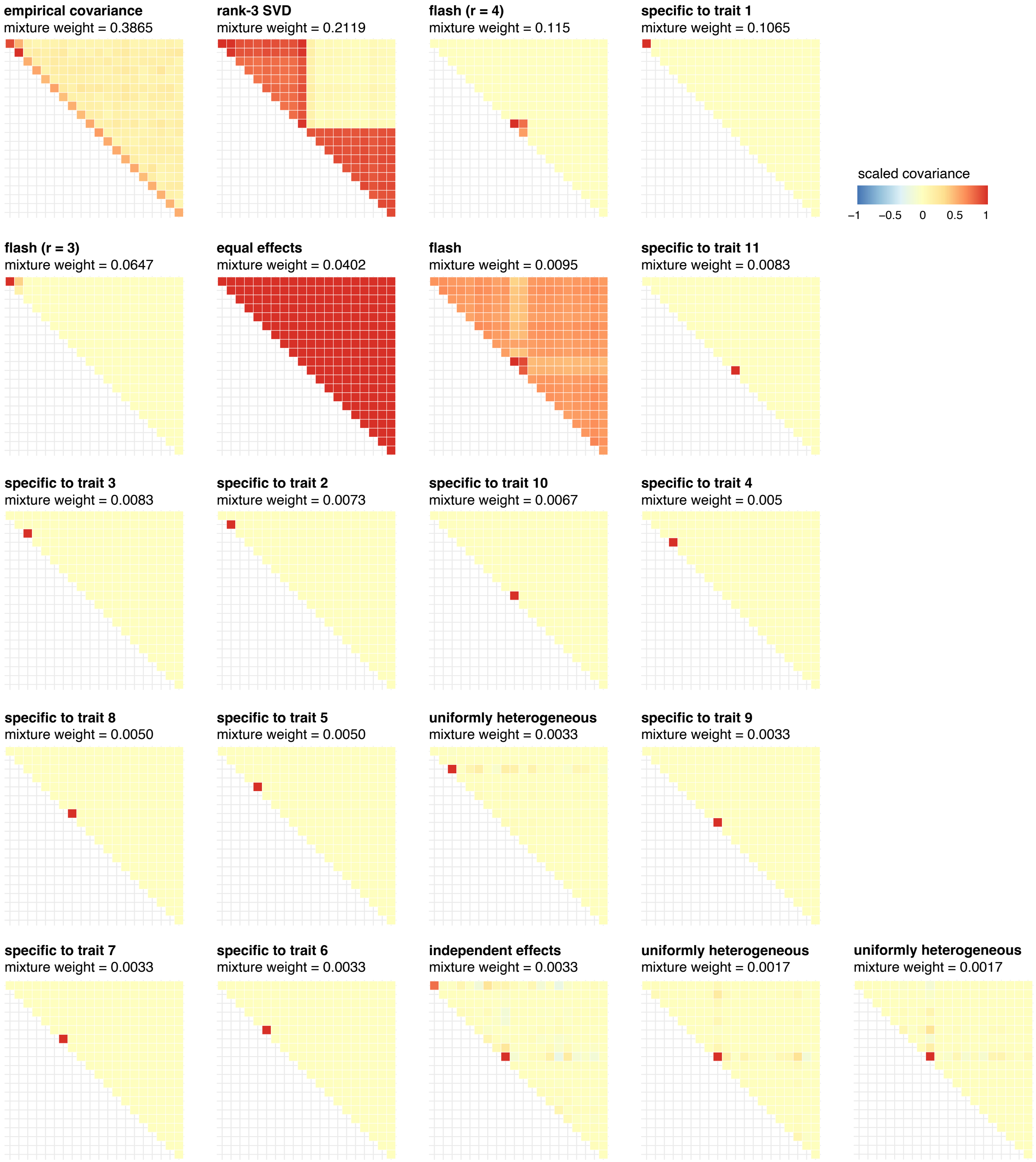

**Supplementary Figure 6. Data-driven prior estimated in Scenario A simulations.** Each plot shows a 20 $\times$ 20 scaled covariance matrix $\boldsymbol{U}_k$ and its corresponding estimated mixture weight $\omega_k$. These covariance matrices and mixture weights describe the "data-driven" prior used in the mvSuSiE analyses of the Scenario A simulated data sets. These covariance matrices capture many of the main effect sharing patterns used to simulate the data (compare to Supplementary Fig. 4) including tissue-specific effects (e.g., mixture components 8–18), independent effects (component 1), effects shared in subgroups (e.g., component 2), and effects shared equally across all tissues (component 6). For visualization purposes only, each plot shows the scaled covariance matrix $\boldsymbol{U}_k/s_k^2$, where $s_k^2$ is the absolute value of the largest (in magnitude) entry of $\boldsymbol{U}_k$, so that all of the plotted values lie between -1 and 1. Each covariance matrix is labeled by the method used to initialize the estimate (see Supplementary Note).

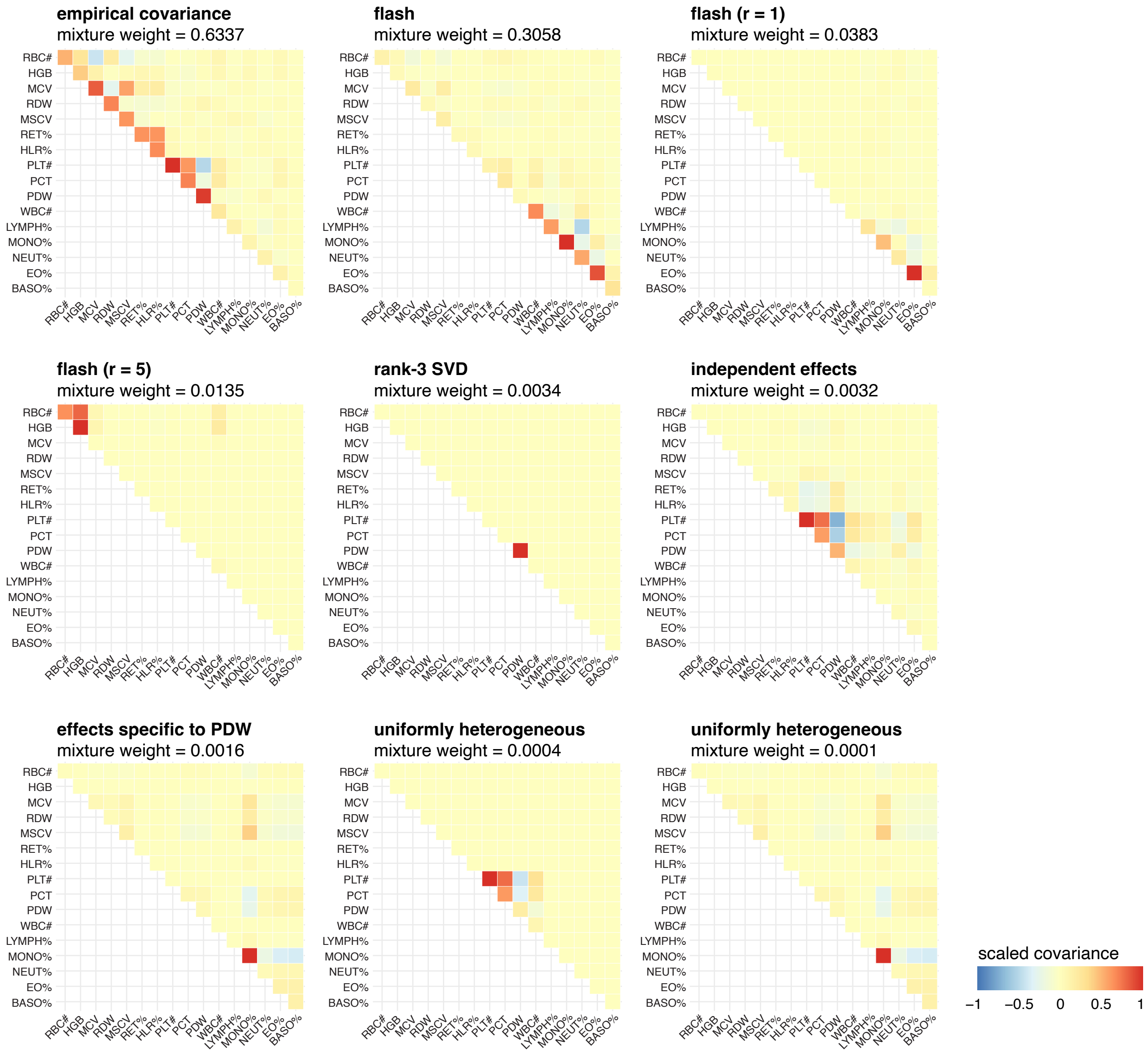

**Supplementary Figure 7. Data-driven prior used in Scenario B simulations.** Each plot shows a 16 × 16 scaled covariance $\boldsymbol{U}_k$ and its corresponding estimated mixture weight $\omega_k$. These covariance matrices and mixture weights describe the “data-driven” prior used in the mvSuSiE analyses of the Scenario B simulated data sets. Compare the covariances and mixture weights shown here to those in Supplementary Fig. 5, which were the covariances and weights used to simulate the data. For visualization purposes only, each plot shows the scaled covariance matrix $\boldsymbol{U}_k/s_k^2$, where $s_k^2$ is the absolute value of the largest (in magnitude) entry of $\boldsymbol{U}_k$, so that all of the plotted values lie between -1 and 1. Each covariance matrix is labeled by the method used to initialize the estimate (see Supplementary Note).

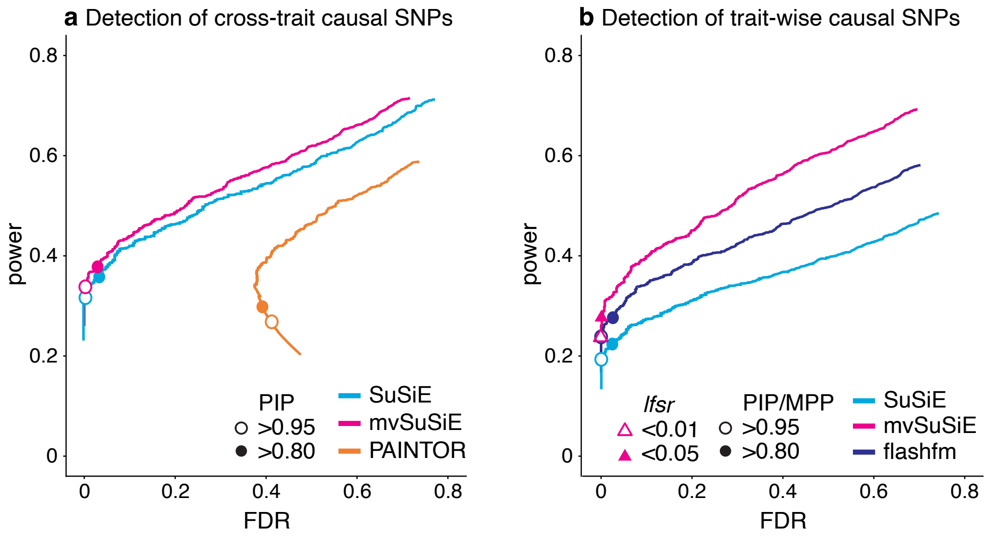

**Supplementary Figure 8. Comparison of fine-mapping methods in simulations with two independent traits and independent effects.** Panel a shows power vs. FDR in identifying cross-trait causal SNPs using PIPs (or max-PIP for SuSiE). In a, FDR and power were calculated as the threshold was varied from 0 to 1 ($n$ = 600 simulations). Note that flashfm does not provide a cross-trait measure so it was not included in a. Panel b shows power vs. FDR in identifying trait-wise causal SNPs. In b, FDR and power were calculated from the 600 simulations as the *marginal posterior probability (MPP)* threshold (flashfm), *PIP* (SuSiE), or *minimum lfsr* (mvSuSiE) was varied from 0 to 1. In a and b, power and FDR at specific thresholds are indicated by the circles and triangles. Note that the thresholds for the different methods are not equivalent or comparable; results are shown at these thresholds for illustration only. Also note that some of the power/FDR results at specific thresholds are not visible because the circles are directly underneath others. Detailed power/FDR statistics at the selected thresholds are given in Supplementary Table 1. Note that PAINTOR does not provide a trait-wise measure so was not included in B.

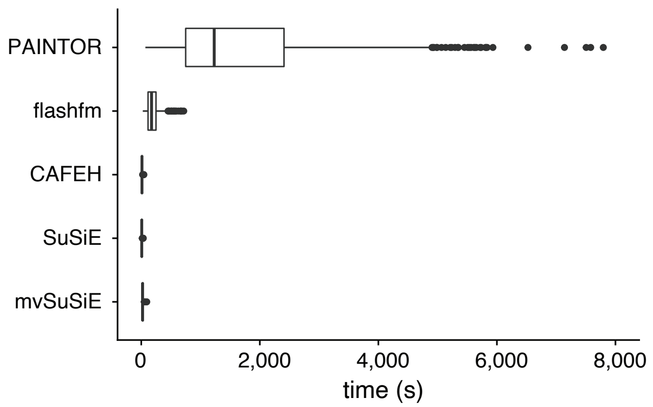

**Supplementary Figure 9. Analysis runtimes in simulations with two independent traits and independent effects.** The box plots summarize the analysis runtimes in the 2-trait simulations ($n$ = 600 simulations). The box plot whiskers depict 1.5$\times$ the interquartile range, the box bounds represent the upper and lower quartiles (25th and 75th percentiles), the center line represents the median (50th percentile), and points represent outliers. The flashfm runtimes include the FINEMAP computations. Note that a single run of PAINTOR that took over 8 h is not shown.

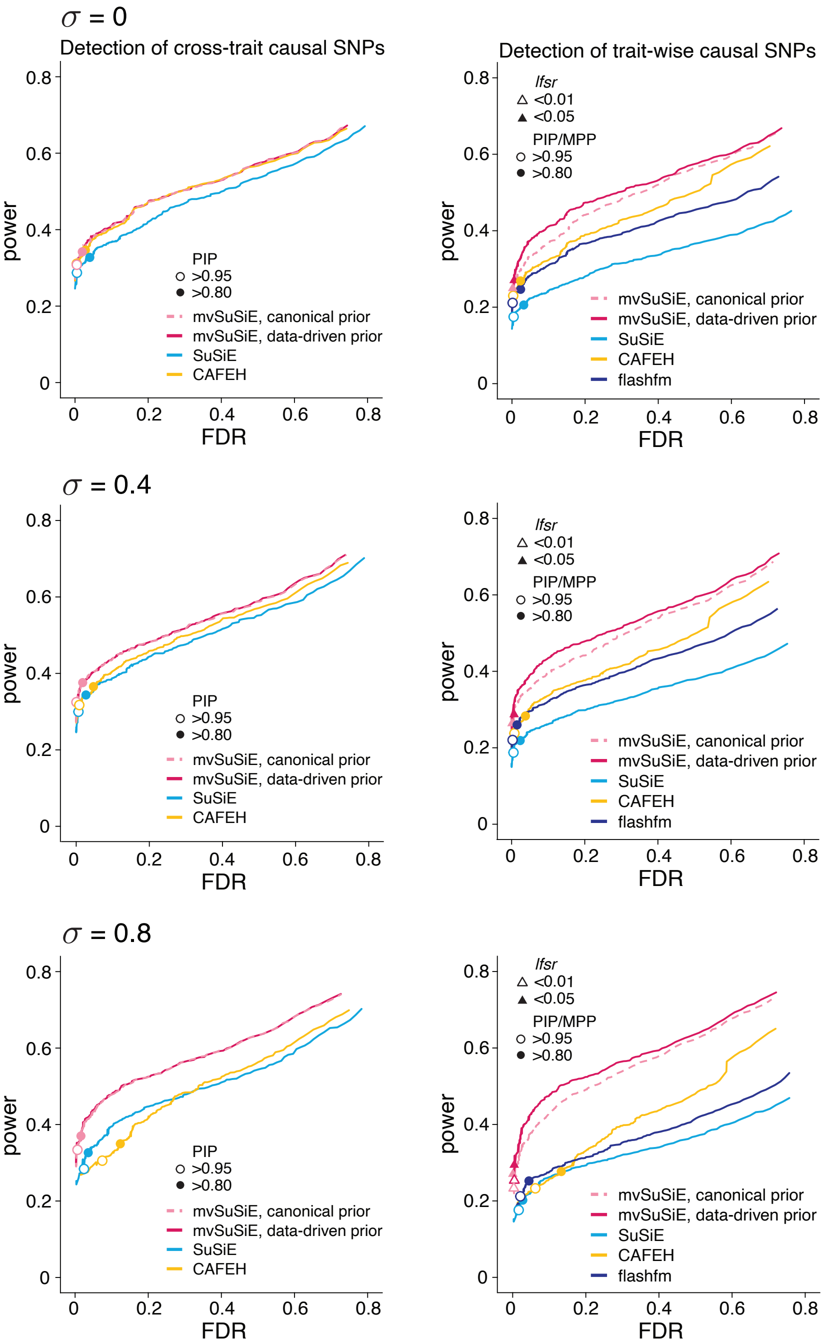

**Supplementary Figure 10. Comparison of fine-mapping methods in simulations with two correlated traits and independent effects, Part A: detection of cross-trait and trait-wise causal SNPs using SNP-wise measures.** In these simulations, two traits were simulated with correlated residuals, with correlation $\sigma$ = 0, 0.4, 0.8. For each setting of $\sigma$, 600 data sets were simulated. The plots on the left-hand side show power vs. FDR in identifying cross-trait causal SNPs using PIPs (or max-PIP for SuSiE). FDR and power were calculated as the threshold was varied from 0 to 1 ($n$ = 600 simulations). Open circles are drawn at a threshold of 0.95. Note that flashfm does not provide a cross-trait measure so it was not included in the left-hand plots. The plots on the right-hand side show power vs. FDR in identifying trait-wise causal SNPs. FDR and power were calculated from the 600 simulations as the *marginal posterior probability (MPP)* threshold (flashfm), *PIP* (SuSiE), or *min-lfsr* (mvSuSiE) was varied from 0 to 1. Power and FDR at specific thresholds are indicated by the circles and triangles. Note that the thresholds shown for the different methods are not necessarily equivalent or comparable; results are shown at these thresholds for illustration only. Also note that some of the power/FDR results are not visible in the plots because the circles lie directly underneath others. Detailed power/FDR statistics at the selected thresholds are given in Supplementary Table 1. The results shown here for $\sigma$ = 0 are the same as the top row of Supplementary Fig. 13 and (for some methods) in Supplementary Fig. 8.

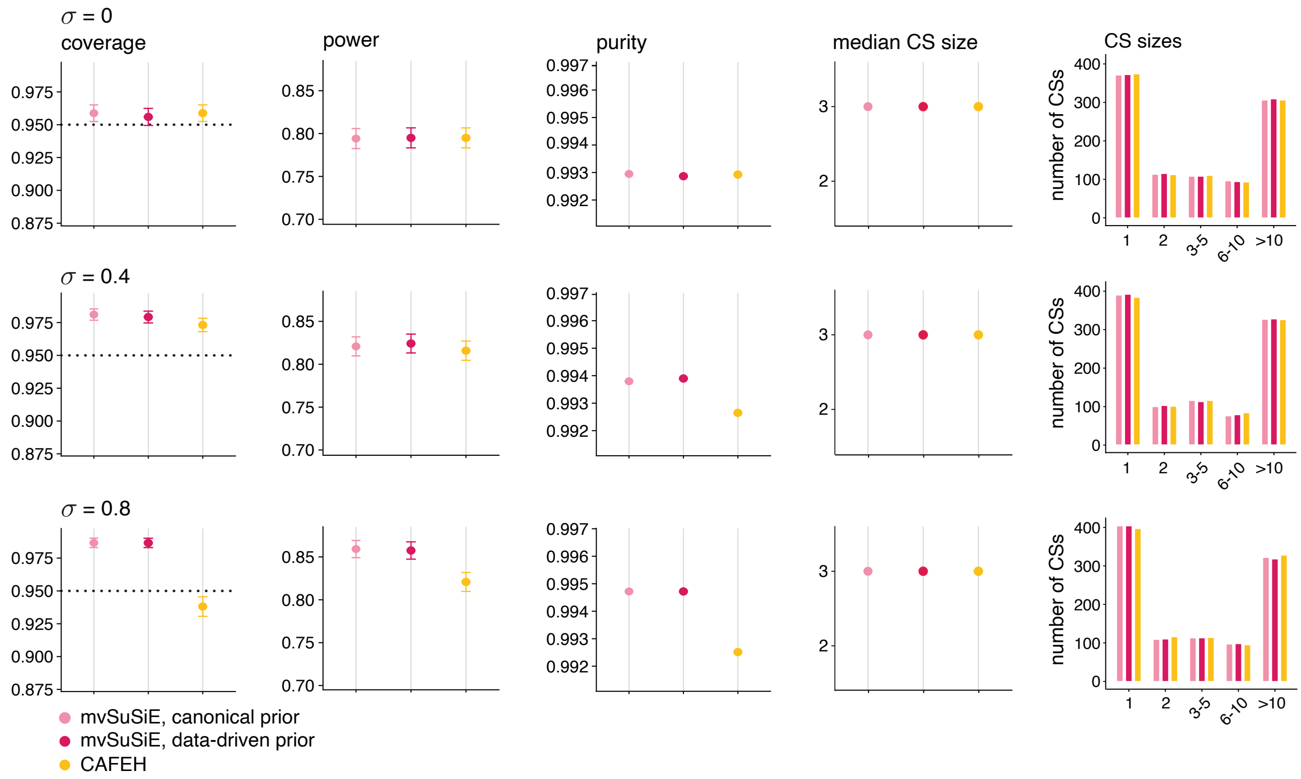

**Supplementary Figure 11. Comparison of fine-mapping methods in simulations with two correlated traits and independent effects, Part B: detection of cross-trait causal SNPs (cross-trait CSs).** The dotted horizontal lines show the target coverage (95%). Error bars show 2 times the empirical s.e. from the results across the $n = 600$ simulations. Note that flashfm does not provide cross-trait CSs and therefore was not included in these plots. See Supplementary Fig. 10 for Part A of these results, and for more details. Also note that the plots shown here for $\sigma = 0$ are the same as the plots in the top row of Supplementary Fig. 14.

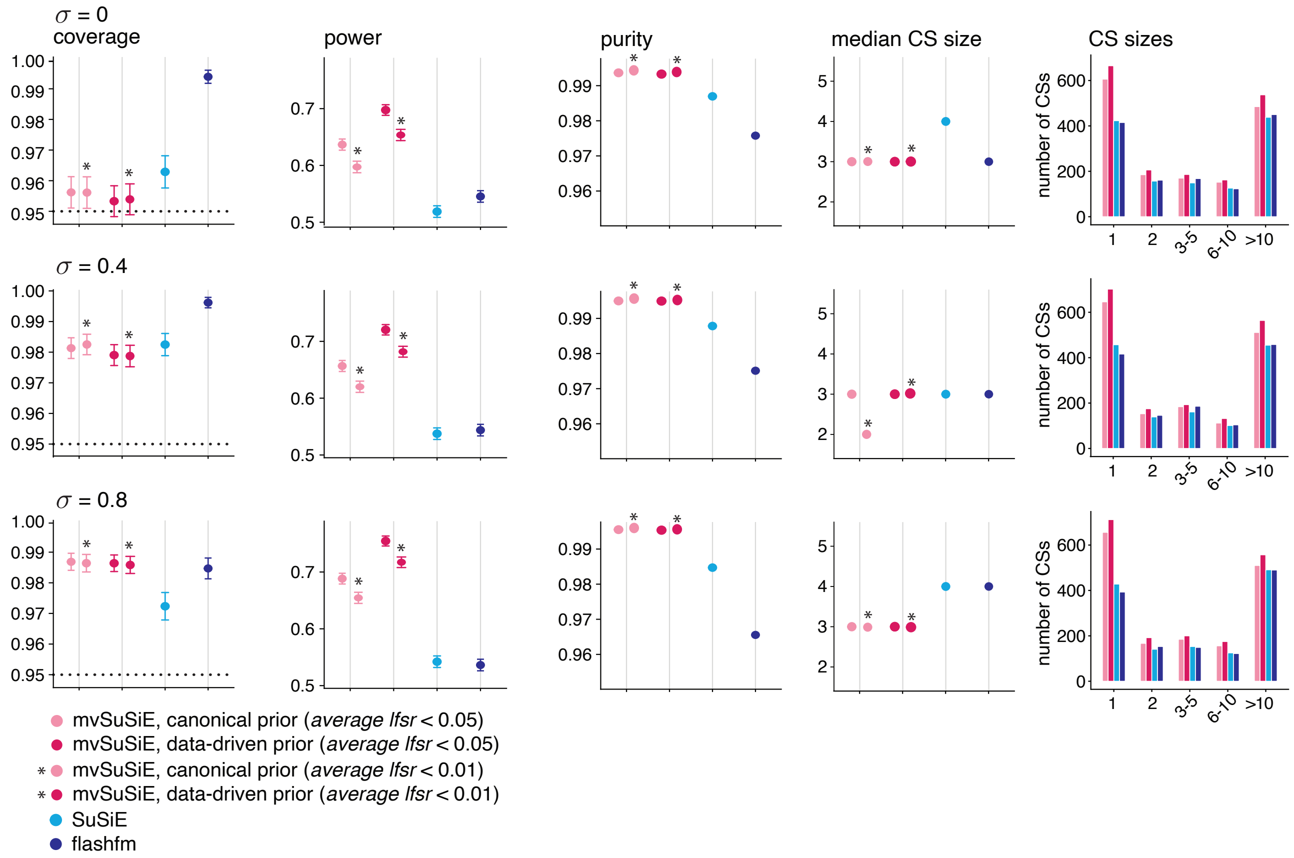

**Supplementary Figure 12. Comparison of fine-mapping methods in simulations with two correlated traits and independent effects, Part C: detection of trait-wise causal SNPs (trait-wise significant CSs).** The dotted horizontal lines show the target coverage (95%). See Supplementary Fig. 10 for Part A of these results, and for more details. CAFEH does not provide trait-wise significant CSs so was not included in these plots. Also note that the plots shown here for $\sigma = 0$ are the same as the plots in the top row of Supplementary Fig. 15.

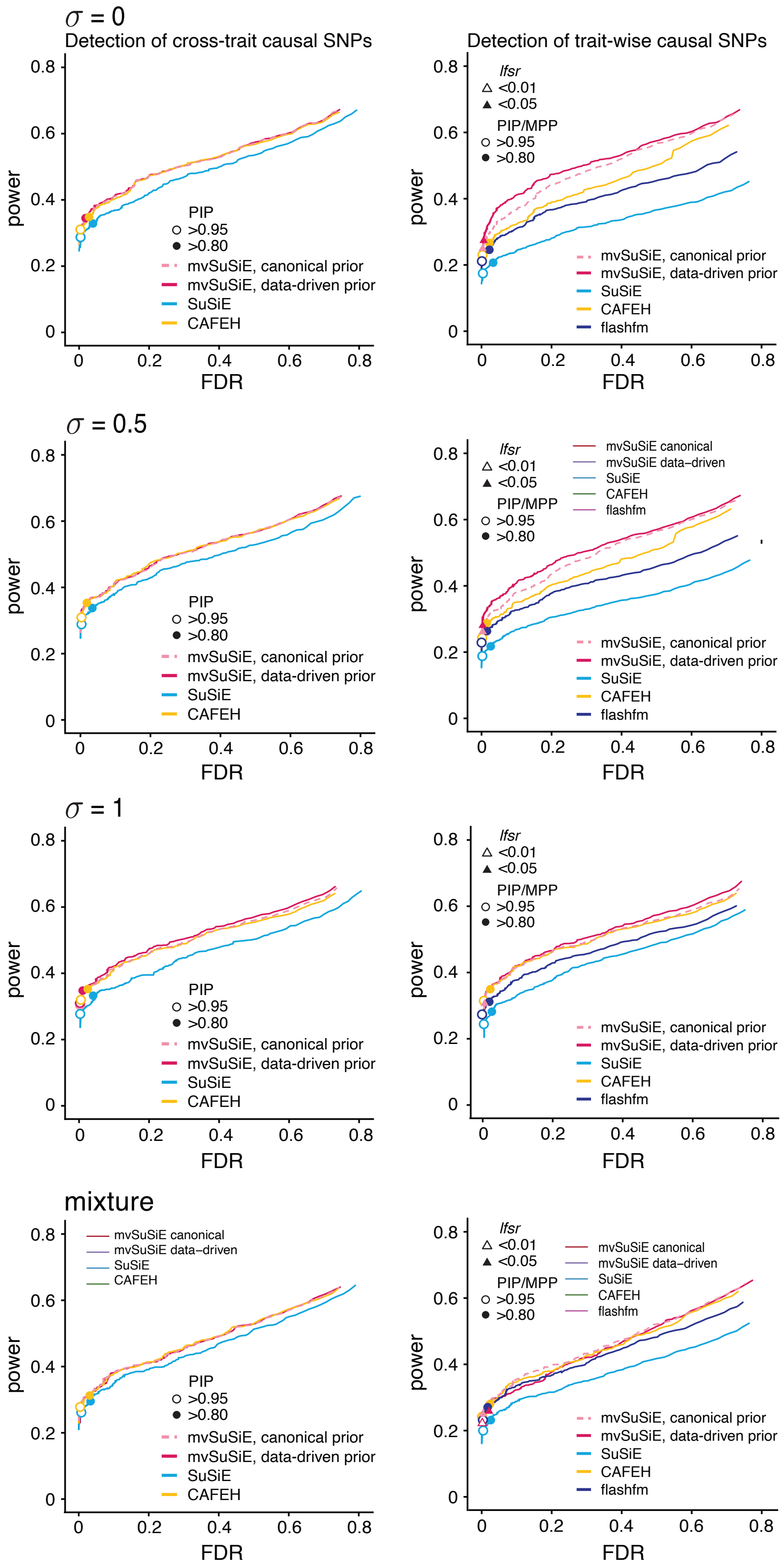

**Supplementary Figure 13. Comparison of fine-mapping methods in simulations with two independent traits and correlated effects, Part A: detection of cross-trait and trait-wise causal SNPs using SNP-wise measures.** See legend on next page.

**Supplementary Figure 13 (previous page). Comparison of fine-mapping methods in simulations with two independent traits and correlated effects, Part A: detection of cross-trait and trait-wise causal SNPs using SNP-wise measures.** In these simulations, two independent traits were simulated with correlated effects, with correlation $\sigma$ = 0, 0.5, 1, and 600 data sets were simulated for each choice of $\sigma$. In a fourth set of simulations (bottom row), the effects were simulated from a mixture of multivariate normals with different covariances (see Methods). The plots on the left-hand side show power vs. FDR in identifying cross-trait causal SNPs using PIPs (or max-PIP for SuSiE). FDR and power were calculated as the threshold was varied from 0 to 1 ($n$ = 600 simulations). Open circles are drawn at a threshold of 0.95. Note that flashfm does not provide a cross-trait measure so it is not included in the left-hand plots. The plots on the right-hand side show power vs. FDR in identifying trait-wise causal SNPs. FDR and power were calculated from the 600 simulations as the *marginal posterior probability (MPP)* threshold (flashfm), *PIP* (SuSiE), or *min-lfsr* (mvSuSiE) was varied from 0 to 1. Power and FDR at specific thresholds are indicated by the circles and triangles. Note that the thresholds shown for the different methods are not necessarily equivalent or comparable; results are shown at these thresholds for illustration only. Also note that some of the power/FDR results are not visible in the plots because the circles lie directly underneath others. Detailed power/FDR statistics at the selected thresholds are given in Supplementary Table 1. The results shown here for $\sigma$ = 0 are the same as the top row of Supplementary Fig. 10 and (for some methods) in Supplementary Fig. 8.

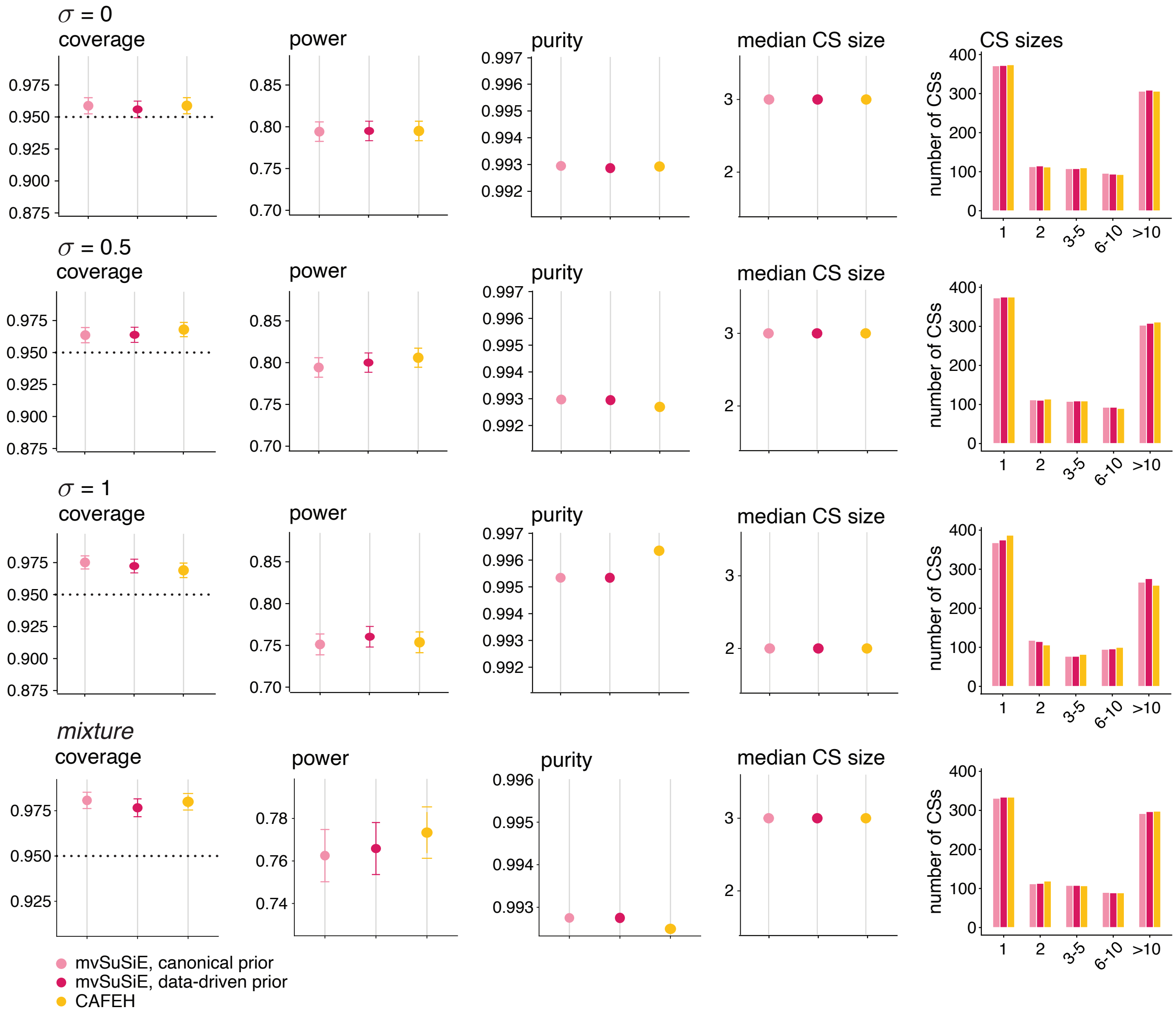

**Supplementary Figure 14. Comparison of fine-mapping methods in simulations with two independent traits and correlated effects, Part B: detection of cross-trait causal SNPs (cross-trait CSs).** See Supplementary Fig. 13 for Part A of these results, and for further explanations. The dotted horizontal lines show the target coverage (95%) and error bars show 2 times the empirical s.e. from the results in the $n = 600$ simulations. Note that flashfm does not provide cross-trait CSs and therefore was not included in these plots. Also note that the plots in the top row are the same as the $\sigma = 0$ plots in Supplementary Fig. 11.

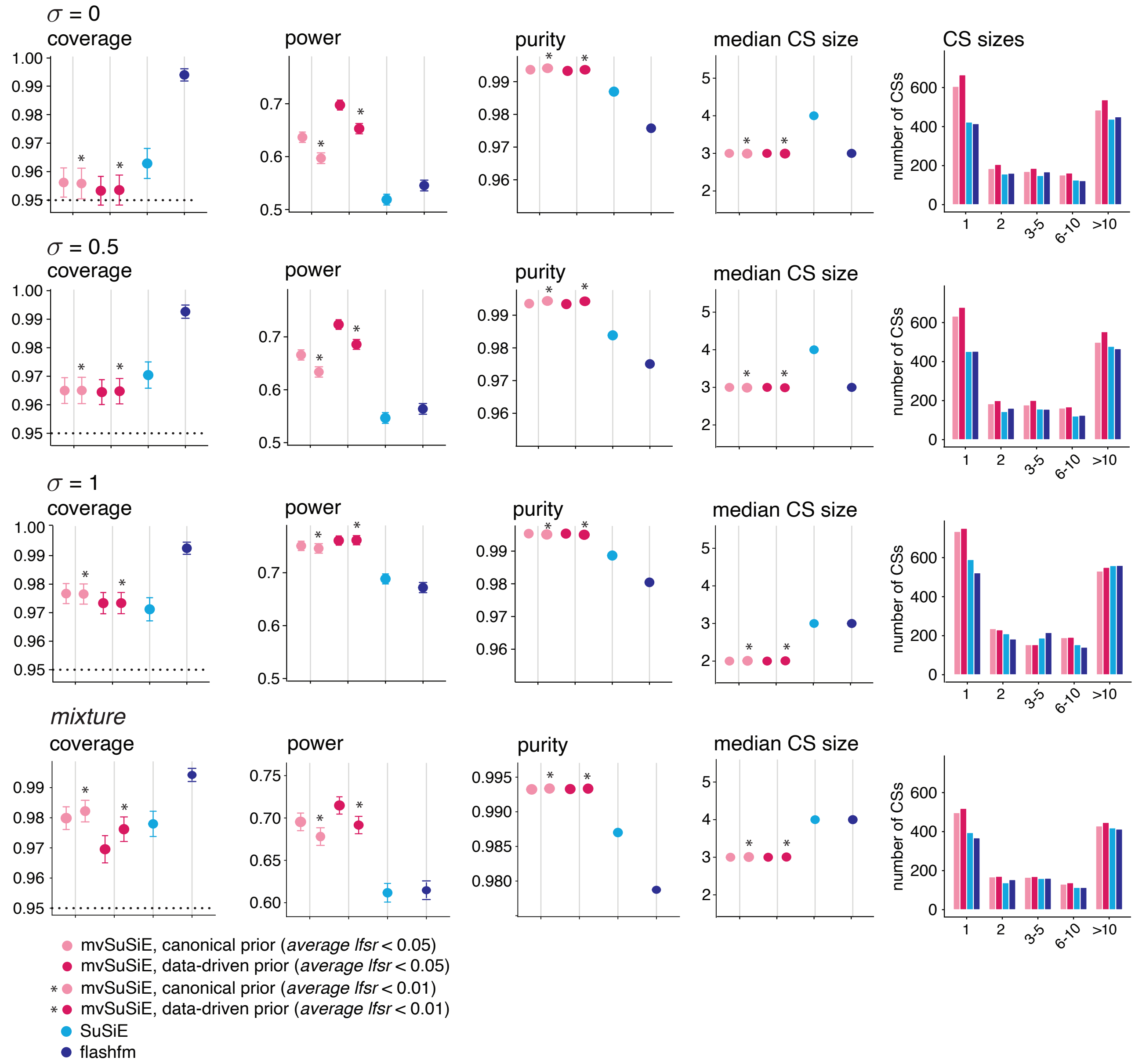

**Supplementary Figure 15. Comparison of fine-mapping methods in simulations with two independent traits and correlated effects, Part C: detection of trait-wise causal SNPs (trait-wise significant CSs).** The dotted horizontal lines show the target coverage (95%). See Supplementary Fig. 13 for Part A of these results, and for further explanations. Note that CAFEH does not provide trait-wise significant CSs so was not included in in these plots. Also note that the plots in the top row are the same as the $\sigma = 0$ plots in Supplementary Fig. 12.

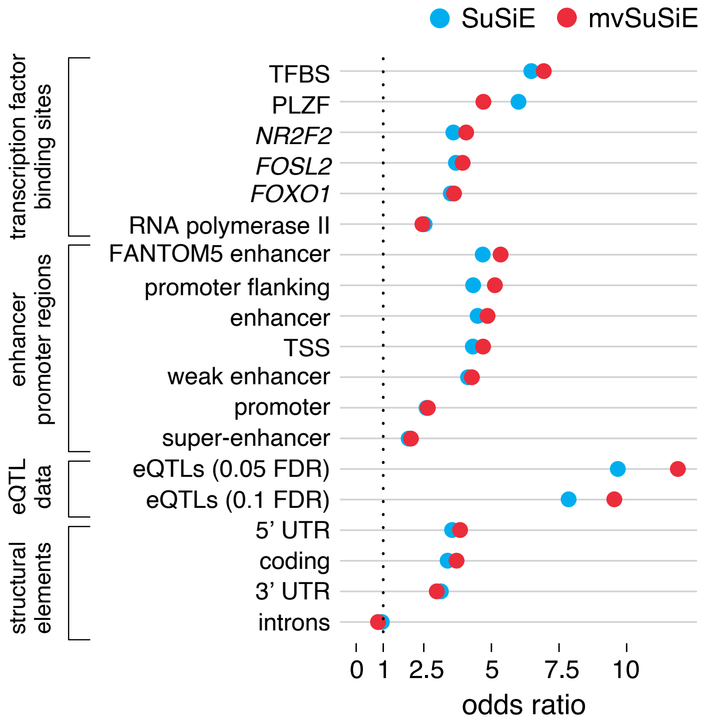

**Supplementary Figure 16. Regulatory enrichment from SuSiE and mvSuSiE fine-mapping of blood cell traits.** The plot shows enrichment odds ratios for SuSiE and mvSuSiE cross-trait fine-mapping results in non-cell-type-specific genomic regulatory annotations [39–43].

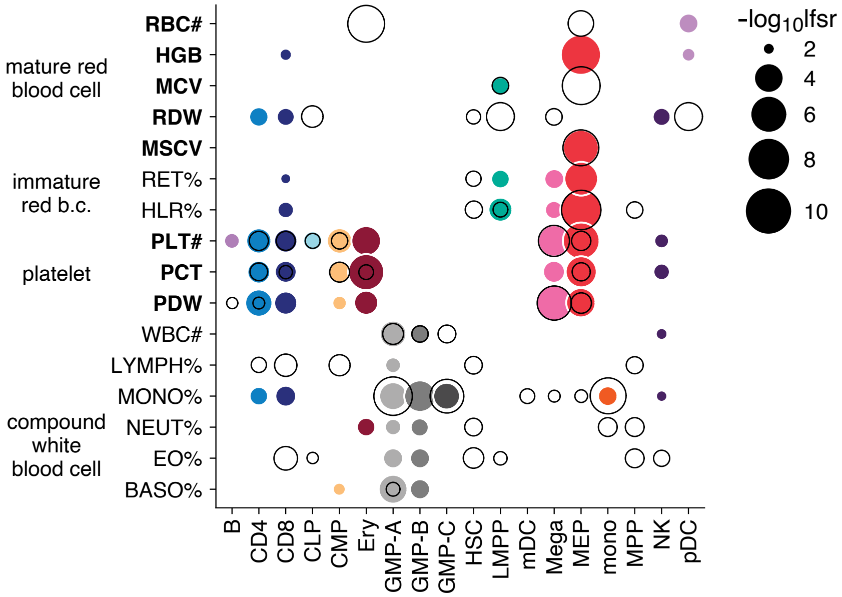

**Supplementary Figure 17. Hematopoietic cell-type enrichment from SuSiE and mvSuSiE fine-mapping of blood cell traits.** The show shows enrichment analysis results for accessible chromatin in hematopoietic cell populations [44]. SuSiE-based enrichments are shown as open circles, and mvSuSiE-based enrichments are colored according to the hematopoietic cell types, similar to [44]. Only enrichments with *lfsr* $< 0.01$ are shown. TFBS = transcription factor binding site; PLZF = promyelocytic leukemia zinc finger protein; mono = monocyte; gran = granulocyte; ery = erythroid; mega = megakaryocyte; CD4 = CD4+ T cell; CD8 = CD8+ T cell; B = B cell; NK = natural killer cell; mDC = myeloid dendritic cell; pDC, = plasmacytoid dendritic cell; MPP = multipotent progenitor; LMPP = lymphoid-primed multipotent progenitor; CMP = common myeloid progenitor; CLP = common lymphoid progenitor; GMP = granulocyte-macrophage progenitor; MEP = megakaryocyte-erythroid progenitor.

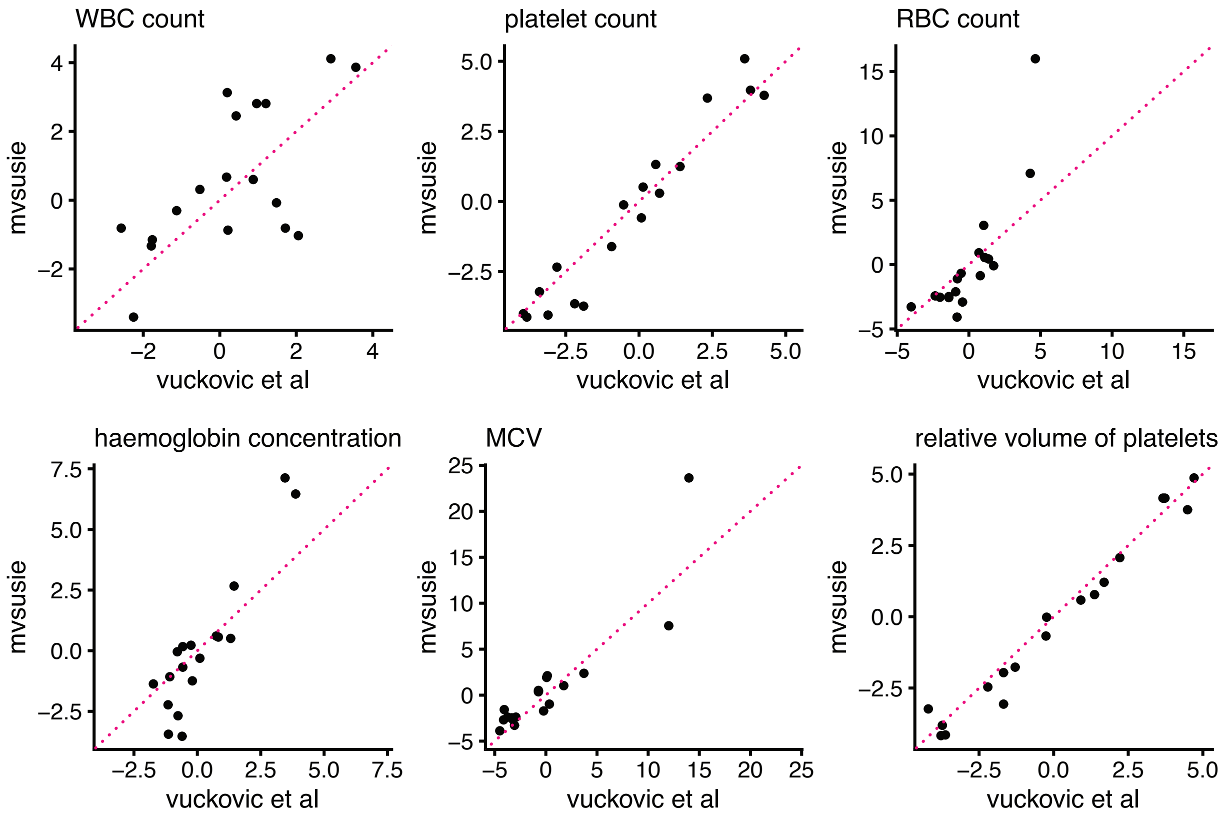

**Supplementary Figure 18. Comparison of gchromVAR enrichments from mvSuSiE vs. Vuckovic et al.** Here we compare the 6 blood cell traits that were included in our analyses and in the fine-mapping analyses of Vuckovic *et al.* [45]. Each plot shows the posterior $z$-scores (posterior means divided by posterior standard deviations) computed from the Vuckovic *et al* enrichment results against the posterior $z$-scores from our enrichment analysis. The Vuckovic *et al* z-scores were downloaded from `https://github.com/bloodcellgwas/manuscript_code`, then posterior $z$-scores were computed using adaptive shrinkage [28].

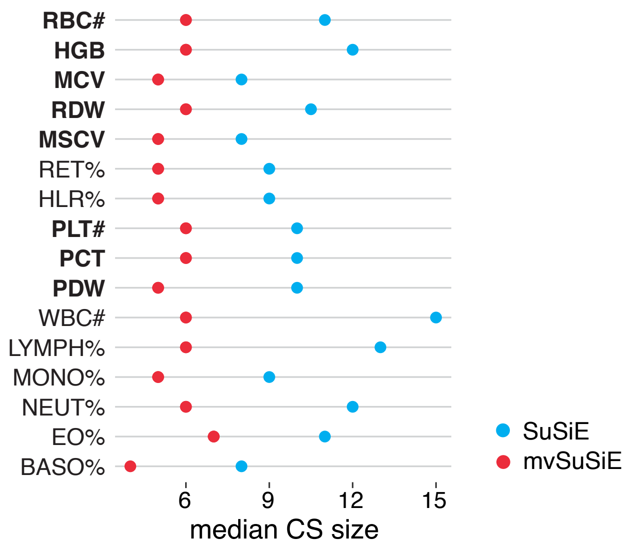

**Supplementary Figure 19. Sizes of trait-wise CSs in SuSiE and mvSuSiE fine-mapping of UK Biobank blood cell traits.** The plot compares the median sizes of the SuSiE CSs and mvSuSiE trait-wise significant CSs, after removing CSs with purity less than 0.5.
